# A pan-variant mRNA-LNP T cell vaccine protects HLA transgenic mice from mortality after infection with SARS-CoV-2 Beta

**DOI:** 10.1101/2022.09.23.509206

**Authors:** Brandon Carter, Pinghan Huang, Ge Liu, Yuejin Liang, Paulo J.C. Lin, Bi-Hung Peng, Lindsay McKay, Alexander Dimitrakakis, Jason Hsu, Vivian Tat, Panatda Saenkham-Huntsinger, Jinjin Chen, Clarety Kaseke, Gaurav D. Gaiha, Qiaobing Xu, Anthony Griffiths, Ying K. Tam, Chien-Te K. Tseng, David K. Gifford

## Abstract

Clinically licensed COVID-19 vaccines ameliorate viral infection by inducing vaccinee production of neutralizing antibodies that bind to the SARS-CoV-2 Spike protein to inhibit viral cellular entry (Walsh et al., 2020; Baden et al., 2021), however the clinical effectiveness of these vaccines is transitory as viral variants arise that escape antibody neutralization (Tregoning et al., 2021; Willett et al., 2022). Vaccines that solely rely upon a T cell response to combat viral infection could be transformational because they can be based on highly conserved short peptide epitopes that hold the potential for pan-variant immunity, but a mRNA-LNP T cell vaccine has not been shown to be sufficient for effective antiviral prophylaxis. Here we show that a mRNA-LNP vaccine based on highly conserved short peptide epitopes activates a CD8^+^ and CD4^+^ T cell response that prevents mortality in HLA-A*02:01 transgenic mice infected with the SARS-CoV-2 Beta variant of concern (B.1.351). In mice vaccinated with the T cell vaccine, 24% of the nucleated cells in lung were CD8^+^ T cells on day 7 post infection. This was 5.5 times more CD8^+^ T cell infiltration of the lungs in response to infection compared to the Pfizer-BioNTech Comirnaty® vaccine. Between days 2 and 7 post infection, the number of CD8^+^ T cells in the lung increased in mice vaccinated with the T cell vaccine and decreased in mice vaccinated with Comirnaty®. The T cell vaccine did not produce neutralizing antibodies, and thus our results demonstrate that SARS-CoV-2 viral infection can be controlled by a T cell response alone. Our results suggest that further study is merited for pan-variant T cell vaccines, and that T cell vaccines may be relevant for individuals that cannot produce neutralizing antibodies or to help mitigate Long COVID.

## Main

Current strategies for COVID-19 vaccine design utilize one or more SARS-CoV-2 Spike protein subunits to primarily activate the humoral arm of the adaptive immune response to produce neutralizing antibodies to the Spike receptor binding domain (RBD) (Walsh et al., 2020; Baden et al., 2021). Vaccination to produce neutralizing antibodies is a natural objective, as neutralizing antibodies present an effective barrier to the viral infection of permissive cells by binding to the RBD and thus blocking cellular entry via the ACE2 receptor. However, the strategy of focusing on Spike as the sole vaccine target has proven problematic as Spike rapidly evolves to produce structural variants that evade antibody-based acquired immunity from vaccination or infection with previous viral variants (Tregoning et al., 2021; Willett et al., 2022). Compared to the original Wuhan variant, novel viral variants are arising that are more contagious (Zhang et al., 2021), and infectious to a broader range of host species (Shuai et al., 2021). Thus, vaccine designers are pursuing a stream of novel Spike variant vaccines.

Multivalent Spike vaccines and bivalent booster vaccines provide protection against multiple known variants of concern (VOC) of SARS-CoV-2 but are not necessarily protective against unknown future variants (Martinez et al., 2021). Mosaic RBD nanoparticles that display disparate SARS-CoV RBDs have been found to produce effective neutralizing antibodies against both SARS-CoV-1 and SARS-CoV-2 (Cohen et al., 2022), but the robustness of mosaic RBD protection against possible future Spike mutations depends upon conserved Spike structural epitopes. In contrast, the vaccine approach we present here depends upon conserved T cell epitopes drawn from the entire viral proteome for protection against future variants.

We show in this study that an immunogenic T cell vaccine with 11 conserved short MHC displayed epitopes (“MIT-T-COVID vaccine”) provides effective prophylaxis against the onset of SARS-CoV-2-induced morbidity and mortality caused by SARS-CoV-2 Beta infection in transgenic mice carrying human HLA-A*02:01. We demonstrate the MIT-T-COVID vaccine causes significant infiltration of CD8^+^ and CD4^+^ T lymphocytes in the lungs. Multiple designs for T cell vaccines for SARS-CoV-2 have been proposed (Liu et al., 2020; Nathan et al., 2021; Heitmann et al., 2022; Pardieck et al., 2022) and are in clinical trials (NCT05113862, NCT0488536, NCT05069623, NCT04954469), but identifying the mechanisms behind the efficacy of pure T cell vaccines remains an open question. Substantial literature suggests that T cell responses are integral to the adaptive immunity to COVID-19 (Moss, 2022). For example, a study that ablated the B cell compartment of the immune system in Spike vaccinated mice found that CD8^+^ T cells alone can control viral infection (Israelow et al., 2021). Pardieck et al. (2022) found that vaccination with a single CD8^+^ T cell epitope conferred protection against mortality from the Leiden-0008/2020 SARS-CoV-2 variant in K18-hACE2 transgenic mice, but unlike our study, required three doses for efficacy, did not engage a CD4^+^ T cell response, and did not identify significant T cell infiltration of the lungs. In addition to specific antibody responses, COVID-19 vaccinees and convalescent patients possess SARS-CoV-2 specific CD8^+^ and CD4^+^ T cells, suggesting the contribution of the T cell compartment to the adaptive immunity to COVID-19 (Sekine et al., 2020), and clinical findings have revealed vaccine-induced T cell responses in B cell-deficient patients (Shree, 2022). It has also been reported that vaccination by WA (Wuhan) Spike in a mouse model failed to produce antibodies fully capable of neutralizing the SA (Beta) variant of SARS-CoV-2, yet immunized mice were protected against Beta strain challenge, leading to the conclusion that T cell responses were sufficient to control Beta SARS-CoV-2 (Kingstad-Bakke et al., 2022). In addition, vaccination with T cell epitope-rich Nucleocapsid protein produced specific T cell responses thought to be causally associated with viral control (Matchett et al., 2021). Intranasal vaccination of mice with SARS-CoV-1 Nucleocapsid followed by challenge with 10^4^ Plaque-forming units of SARS-CoV-1 prevented mortality in 75% of the mice (Zhao et al, 2016). Combined Spike and Nucleocapsid vaccination improved viral control compared to Spike vaccination alone in preclinical models, while CD8^+^ T cell depletion demonstrated the role of CD8^+^ T cells in viral control and protection from weight loss (Hajnik et al., 2022).

### Vaccine Design

We designed a COVID-19 T cell vaccine using T cell epitopes that are unchanged over 19 presently known SARS-CoV-2 variants of concern (VOC) (Methods). Since T cell epitopes can originate from any part of the viral proteome, they can be drawn from portions of the proteome that are likely to be evolutionarily stable. The prediction of stable epitopes can be accomplished by historical analysis of thousands of viral variants (Liu et al., 2020), structural analysis (Nathan et al., 2021), or the functional analysis of mutations lethal to the virus.

Beginning with stable epitopes as candidates, our T cell vaccine design proceeds by vaccine epitope selection that optimizes population coverage where every vaccinated individual is predicted to experience on average multiple immunogenic peptide-HLA hits (Liu et al., 2021; Liu et al., 2022).

The MIT-T-COVID vaccine realizes an *n-times* coverage objective by encoding multiple epitopes for each target MHC class I and II diplotype to (1) expand diverse sets of T cell clonotypes to fight viral infection, (2) accommodate variations in epitope immunogenicity between vaccinees, and (3) reduce the chances that viral evolution will lead to immune system escape (Liu et al., 2022). The MIT-T-COVID vaccines consists of eight MHC class I epitopes and three MHC class II epitopes (Figure 1A, Methods, Supplemental Table 1). The MHC class I and II vaccine peptides are encoded into a single mRNA construct for delivery with the same Acuitas LNP delivery platform that is used by the Pfizer-BioNTech Comirnaty® vaccine (Figure 1A). The eight MIT-T-COVID MHC class I epitopes are the HLA-A*02:01 subset of the MHC class I *de novo* MIRA only vaccine design of Liu et al. (2021) that used combinatorial optimization to select vaccine epitopes to maximize *n-times* population coverage over HLA haplotype frequencies (Liu et al., 2022). For inclusion in the assembled construct, the eight MHC class I vaccine peptides were randomly shuffled, and alternate peptides were flanked with five additional amino acids at each terminus as originally flanked in the SARS-CoV-2 proteome. Selected epitopes were flanked to test if flanking enhanced or impaired epitope presentation.

**Figure 1.**
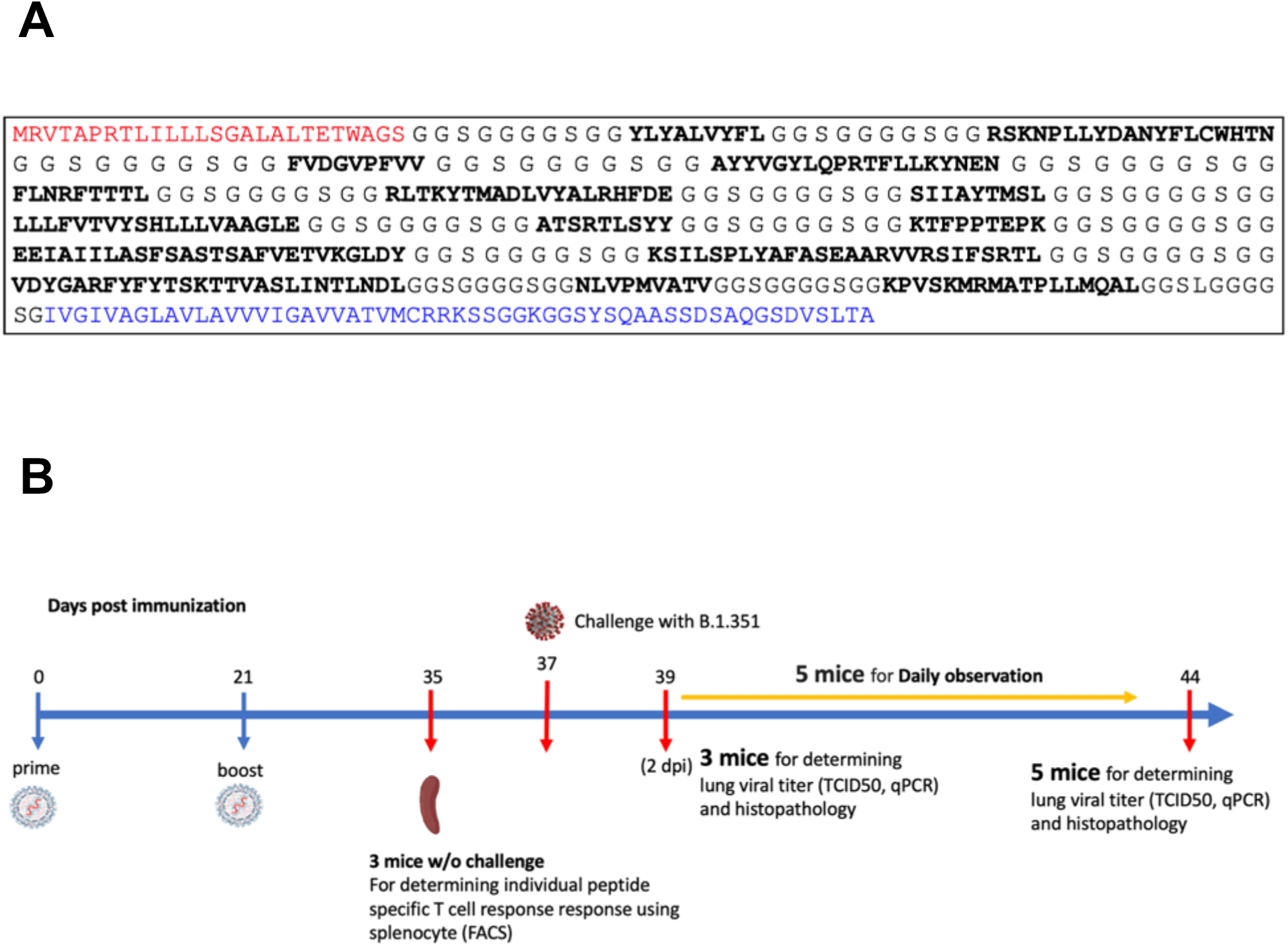
(A) Assembled vaccine construct containing a secretion signal sequence (red), peptides (bold) joined by non-immunogenic glycine/serine linkers, and an MHC class I trafficking signal (blue). (B) Study design.

The three MHC class II epitopes were selected by considering all SARS-CoV-2 proteome windows of length 13-25 and selecting conserved peptides that are predicted to be displayed by the mouse MHC class II allele H-2-IAb (Methods). The mRNA construct encodes a secretion signal sequence at the N-terminus and an MHC class I trafficking signal (MITD) at the C-terminus (Kreiter et al., 2008). Peptide sequences are joined by non-immunogenic glycine/serine linkers (Sahin et al., 2017). The construct also included control peptides for HLA-A*02:01 and H-2-IAb (CMV pp65: NLVPMVATV for HLA-A*02:01 and Human CD74: KPVSKMRMATPLLMQAL for H-2-IAb) (Vita et al., 2018).

### Challenge Study

We vaccinated with three test articles: a negative control injection (PBS), the Pfizer-BioNTech Comirnaty® vaccine (wastage that was refrozen and then thawed), and the MIT-T-COVID vaccine (Figure 1B). We vaccinated HLA-A*02:01 human transgenic mice with these three vaccines, vaccinating 11 age-matched male mice with each vaccine. Comirnaty® or MIT-T-COVID vaccinations contained of 10 μg of mRNA. The mice were vaccinated at Day 0 and boosted at Day 21 (Methods). At Day 35 three mice from each group were sacrificed for immunogenicity studies, and at Day 37 the remaining eight mice were challenged intranasally (I.N.) with 5 × 10^4^ TCID_50_ of the SARS-CoV-2 B.1.351 variant. Challenged mice were subjected to daily monitoring for the onset of morbidity (i.e., weight changes and other signs of illness) and any mortality. At Day 39 (2 days post infection) three mice were taken from each group and sacrificed to determine viral burdens and to perform lung histopathology. At Day 44 (7 days post infection) the remaining mice were sacrificed to determine viral burdens and to perform lung histopathology.

### Immunogenicity of vaccines

Vaccination by the MIT-T-COVID vaccine expanded CD8^+^ and CD4^+^ T cells that expressed interferon gamma (IFN-γ) or tumor necrosis factor alpha (TNF-α) when queried by vaccine epitopes (Figures 2A-B). The observed variability of immunogenicity of MIT-T-COVID epitopes in convalescent COVID patients (Supplemental Table 1) and in the present study supports our use of multiple epitopes per MHC diplotype for *n-times* coverage. Vaccination by Comirnaty® produced no significant T cell responses to vaccine epitopes when compared to PBS. CD8^+^ T cells that are activated by the CD8-4 epitope (YLQPRTFLL, Spike 269-277) are expanded in animals vaccinated with the MIT-T-COVID vaccine (IFN-γ 1.32% ± 0.53% of CD8^+^ T cells, *P* = 0.0087 vs. PBS; TNF-α 0.38% ± 0.09% of CD8^+^ T cells, *P* = 0.0149 vs. PBS; Figure 2A).

**Figure 2.**
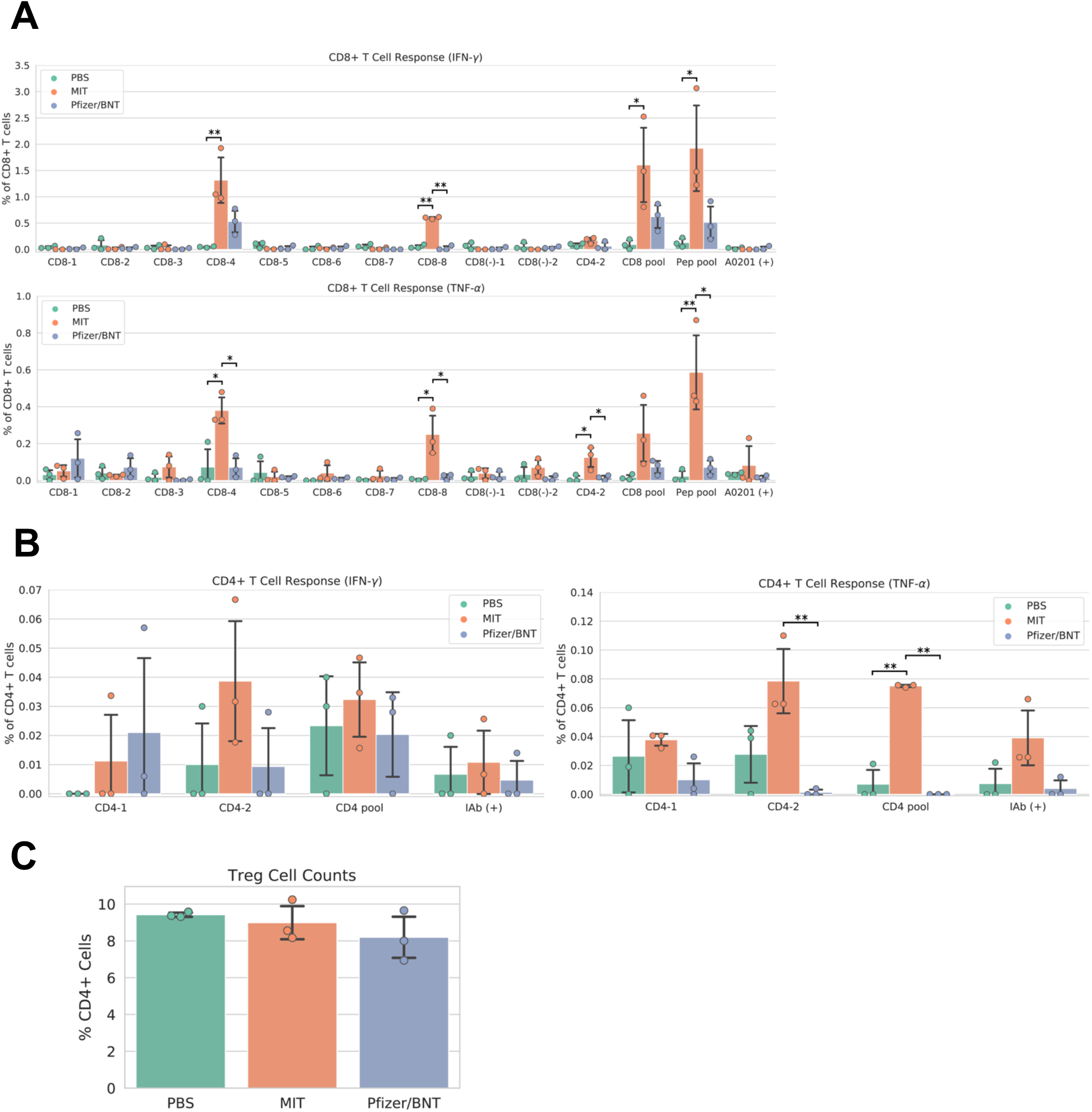
Vaccine immunogenicity. (A) CD8^+^ T cell responses, (B) CD4^+^ T cell responses, (C) Foxp3^+^ regulatory T cells (Tregs) as a percentage of all CD4^+^ cells. The CD8 pool includes MHC class I peptides CD8-1—CD8-8 (Supplemental Table 1). The CD4 pool includes MHC class II peptides CD4-1 and CD4-2. The Pep pool includes all query peptides in Supplemental Table 1 except CD4-3. Error bars indicate the standard deviation around each mean. *P* values were computed by one-way ANOVA with Tukey’s test. \**P* < 0.05, \*\**P* < 0.01.

Similarly, the CD8-8 epitope (TVYSHLLLV, ORF3a 89-97) activated CD8^+^ T cells that are expanded by the MIT-T-COVID vaccine (IFN-γ 0.60% ± 0.02% of CD8^+^ T cells, *P* = 0.001 vs. PBS; TNF-α 0.25% ± 0.12% of CD8^+^ T cells, *P* = 0.015 vs. PBS) and the CD4-2 epitope (RFYFYTSKTTVASLIN, ORF1ab 1421-1436) activated CD4^+^ T cells are expanded by the MIT-T-COVID vaccine (TNF-α 0.078% ± 0.027%, *P* = 0.058 vs. PBS, *P* = 0.010 vs. Comirnaty®, IFN-γ not significant, Figure 2B). The lack of HLA-A*02:01 transgenic animal response to certain CD8^+^ epitopes that were immunogenic in patients (Supplemental Table 1) is consistent with past studies of transgenic mouse models (Kotturi et al., 2009) and the variability of immunogenicity of epitopes between in-bred mice (Croft et al., 2019). We vaccinated an additional cohort of female HLA-A*02:01 transgenic mice (Methods), and results are shown in Supplemental Figure 8. We also vaccinated female mice with synthetic peptides mixed with poly IC adjuvant and found no significant T cell responses compared to PBS controls (Supplemental Figures 9A-B, Methods). We found a significant increase in the number of effector and memory CD8^+^ CD44^+^ T cells in MIT-T-COVID and Comirnaty® vaccinated mice compared to mice vaccinated with Peptide/poly IC or PBS (Supplemental Figure 9C).

T cells that lack IFN-γ responses can produce effective immune mediators (Nakiboneka et al., 2019). We found that the fraction of CD4^+^ T cells that are Foxp3^+^ were not expanded in Comirnaty® or MIT-T-COVID vaccinated animals, suggesting that fraction of T regulatory cells that could induce tolerance were not expanded by vaccination (Figure 2C).

### Vaccine efficacy against the onset of morbidity and mortality and the viral loads in challenged mice

Upon viral challenge, both PBS and MIT-T-COVID vaccinated animals exhibited a more than 10% weight loss by Day 3 (PBS mean weight reduction 11.386% ± 1.688%, MIT mean weight reduction 10.851% ± 0.641%), with the MIT-T-COVID vaccinated animals beginning to recover from Day 4 onward (Figure 3A). The weight phenotype was mirrored by the clinical score phenotype, with PBS animals not recovering and MIT-T-COVID vaccinated animals improving from Day 5 onward (Figure 3B, Methods). The Comirnaty® vaccine protected animals from significant weight loss and poor clinical scores.

**Figure 3.**
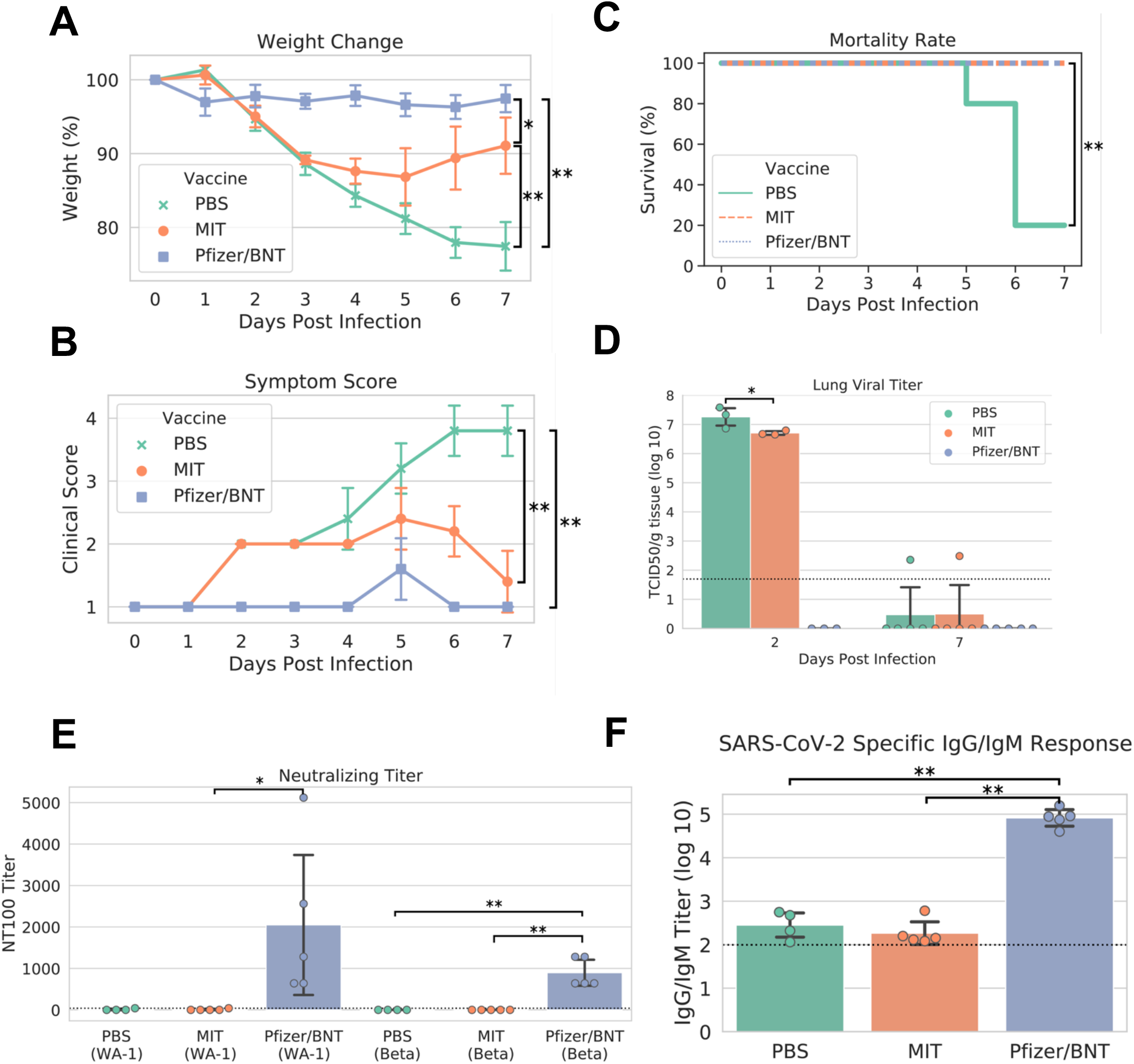
Study phenotypic data, lung viral titer, and vaccine antibody responses. (A) Weights vs. days post infection, (B) clinical scores vs. days post infection, (C) Kaplan-Meier mortality curve (mortality at 80% weight loss), (D) lung viral titer, (E) maximum serum dilution that provided 100% neutralization of viral infection *in vitro*, (F) IgG/IgM titer measured by ELISA against cell lysate infected with WA-1 SARS-CoV-2. Dotted lines in (D)-(F) indicate assay limits of detection. Error bars indicate the standard deviation around each mean. *P* values were computed by one-way ANOVA with Tukey’s test except (C) *P* values were computed using the logrank test. \**P* < 0.05 and \*\**P* < 0.01.

When the yields of infectious progeny virus were measured at 2 days post infection (dpi), we noted that mice immunized with MIT-T-COVID vaccine had 6.706 ± 0.076 log10 TCID_50_/g, compared to 7.258 ± 0.367 log10 TCID_50_/g in the PBS control, representing a moderate 3.6-fold reduction in viral replication (*P* = 0.046; Figure 3D). In contrast, mice immunized with Comirnaty® had infectious viral titers that were below the detection limits at 2 dpi. Infectious viral titer was not significant for all test articles at 7 dpi (Figure 3D). Lung viral mRNA levels measured by qPCR followed the same trends as infectious virus progeny in the lungs, with Comirnaty® showing no significant levels. Despite a reduced content of total and sub-genomic viral RNAs associated with MIT-T-COVID vaccine samples as assessed by qPCR, the reduction was insignificant, compared to those of PBS control (Supplemental Figure 1).

All the Comirnaty® and MIT-T-COVID vaccine animals survived to 7 dpi, when the study was terminated, with the difference in clinical scores of these two vaccine groups becoming insignificant (Figure 3C). In contrast, four of five PBS control animals had been euthanized because of weight loss > 20%. The survival of all five animals vaccinated with Comirnaty® and the MIT-T-COVID vaccine was significant vs. PBS (*P* = 0.0053, logrank test).

We noted that mice immunization with the Comirnaty® vaccine elicited substantial specific antibodies capable of completely neutralizing Beta and WA-1 variants of SARS-CoV-2 with 100% neutralizing titers (NT_100_) of 896 ± 350 and fold 2048 ± 1887, respectively (Figure 3E). The higher neutralizing titer against WA-1 is expected as it is variant matched to Comirnaty®. As expected for a peptide-based vaccine, no neutralizing titer could be readily detected in the serum from mice immunized with MIT-T-COVID vaccine or PBS. Total specific IgG/IgM antibodies were also measured by ELISA against a cell lysate prepared from WA-1-infected Vero E6 cells. Comirnaty® has a log10 IgG/IgM titer of 4.915 ± 0.213, while the log10 titer produced by PBS was 2.452 ± 0.321 and the MIT-T-COVID vaccine produced a log10 titer of 2.266 ± 0.291 (Figure 3F). The low but detectable titers in PBS and MIT-T-COVID vaccine-immunized mice may represent an early IgM response to viral infection.

All lung samples were subjected to immunohistochemistry (IHC) staining for SARS-CoV-2 spike protein (Supplemental Figure 2). We found specimens vaccinated with PBS exhibited extensive staining indicative of viral infection throughout the epithelium of both the bronchioles and the alveolar sacs, with the viral infection appearing more intense at 2 dpi. Although viral infection is significantly reduced by 7 dpi, viral antigen was still readily detectable throughout alveoli. In comparison, specimens vaccinated with the MIT-T-COVID vaccine exhibited similarly extensive viral infection at 2 dpi throughout the bronchiolar and alveolar epithelia, albeit somewhat reduced in intensity. However, by 7 dpi, viral infection was significantly reduced in both extent and intensity, with brown puncta being detected only in a few alveoli scattered throughout the tissue. Contrasted with both PBS and MIT-T-COVID vaccinated specimens, the Comirnaty®-vaccinated specimens exhibited significantly reduced viral infections at both 2 and 7 dpi. Apart from a single area at 7 dpi (see Supplemental Figure 3), viral antigen was undetected at both timepoints in Comirnaty® vaccinated animals.

Paraffin-embedded and H&E-stained lung specimens of differentially immunized mice, harvested at 2 and 7 dpi, were subjected to histopathological examination (Supplemental Figure 4). We found that at 7 dpi mice immunized with MIT-T-COVID vaccine exhibited extensive lymphocytic infiltrations in perivascular regions and spaces from around bronchi, bronchioles, to alveoli. Fewer infiltrations were found in mice immunized with either Comirnaty® or PBS. Additionally, these infiltrations only localized at perivascular regions around bronchi and large bronchioles. Despite the less intensive and localized inflammatory infiltrates, we also noted widespread congestion, hemorrhage, and few foci of thromboembolism were exclusively observed within the lungs of Comirnaty®-immunized mice but not others (Supplemental Figure 4). We also noted the lung histopathology was milder but the same pattern at 2 dpi than those of 7 dpi (data not shown).

**Figure 4.**
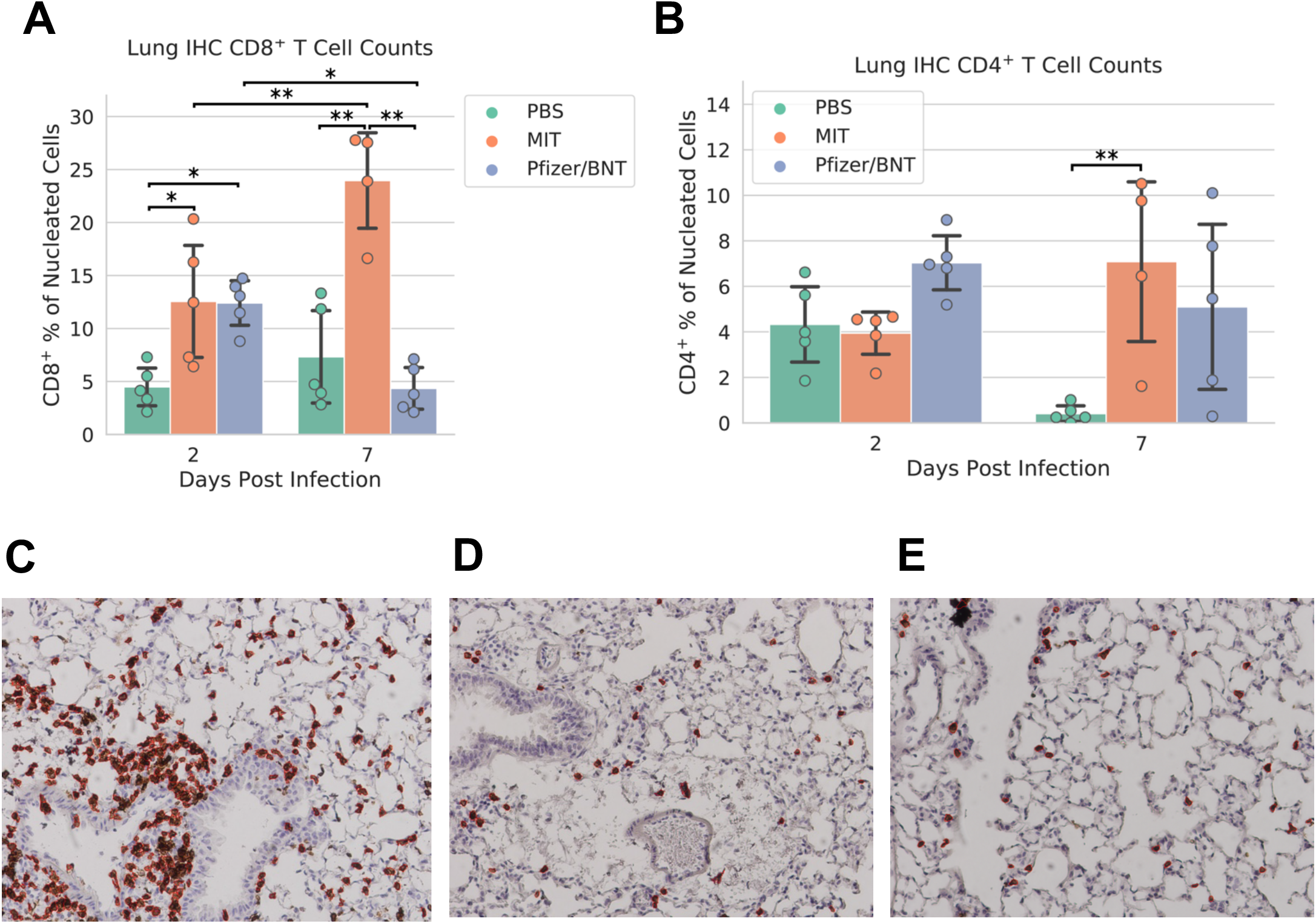
Lung immunohistochemistry for CD8^+^ and CD4^+^ cells. Counts of (A) CD8^+^ and (B) CD4^+^ T cells expressed as a percentage of all nucleated cells visible in each field from lung tissue. Example CD8^+^ stain images at 7 dpi for (C) MIT-T-COVID, (D) Pfizer/BNT, and (E) PBS vaccinated animals. Lung samples were subjected to IHC staining for CD8 (brown) with hematoxylin counterstain (blue). Images were taken at 10x magnification. Red outlines in (C)-(E) indicate CD8^+^ cells identified and counted by CellProfiler software (Methods). Error bars indicate the standard deviation around each mean. *P* values were computed by two-way ANOVA with Tukey’s test. \**P* < 0.05 and \*\**P* < 0.01. See also Supplemental Figures 5-7.

Lung specimens were subjected to IHC staining for CD8^+^ and CD4^+^ cells at both 2 and 7 dpi (Figure 4; Supplemental Figures 5-7). At 2 dpi, we found a significant increase in CD8^+^ T cells infiltrating the lungs in mice immunized with MIT-T-COVID (12.6% ± 5.91% of all nucleated cells were CD8^+^) or Comirnaty® (12.4% ± 2.35%) compared to mice immunized with PBS (PBS mean 4.48% ± 1.99%; *P* = 0.044 vs. MIT-T-COVID, *P* = 0.050 vs. Comirnaty®; Figure 4A).

At 7 dpi, we found a significant increase in CD8^+^ T cells infiltrating the lungs in mice immunized with MIT-T-COVID (24.0% ± 5.21% of all nucleated cells were CD8^+^) compared to mice immunized with Comirnaty® (Comirnaty® mean 4.35% ± 2.20%, *P* = 0.001) or PBS (PBS mean 7.32% ± 4.88%, *P* = 0.001; Figure 4A). We observed a significant increase in CD8^+^ T cells infiltrating the lungs in mice immunized with MIT-T-COVID between 2 and 7 dpi (*P* = 0.0039), and a significant decrease in CD8^+^ T cells infiltrating the lungs in mice immunized with Comirnaty® between 2 and 7 dpi (*P* = 0.044; Figure 4A). At 7 dpi, we also found a significant increase in CD4^+^ T cells infiltrating the lungs in mice immunized with MIT-T-COVID (7.09% ± 4.05%) compared to mice immunized with PBS (PBS mean 0.41% ± 0.39%; *P* = 0.0062; Figure 4B). In the unchallenged cohort of female mice, we found an increase in CD8^+^ T cells infiltrating the lungs in mice immunized with MIT-T-COVID (1.12% ± 0.59% of all nucleated cells were CD8^+^) or Comirnaty® (0.87% ± 0.56%) compared to mice immunized with PBS (0.09% ± 0.10%; *P* = 0.001 vs. MIT-T-COVID, *P* = 0.003 vs. Comirnaty®; Supplemental Figures 10-11).

## Discussion

Existing Spike vaccines produce a T cell response that is thought to be important for vaccine effectiveness and durability (Moss, 2022). However, the T cell response induced by a Spike vaccine may be less effective in promoting durable immune responses than the T cell response induced by a pure T cell vaccine. Spike based T cell responses may contain epitopes that are subject to evolutionary change (de Silva et al., 2021; Redd et al., 2021), or Spike epitopes may not be as well presented or immunogenic as epitopes drawn from a more diverse set of SARS-CoV-2 proteins. Thus, T cell augmentation strategies for Spike vaccines (Liu et al., 2021) and pure T cell vaccines will require further exploration to unravel the hierarchy of immune responses to their components, and how to structure a vaccine for optimal prophylaxis. In patients with impaired antibody responses, T cell vaccines would eliminate the burden of non-immunogenic B cell epitopes (Benjamini et al., 2022) and provide immune protection, at least to a certain extent, against infection.

The marginal IgG/IgM antibody titers that were not neutralizing elicited by mice immunized with the MIT-T-COVID vaccine indicates that the protective mechanism of the vaccine was likely based on a T cell response to the virus. The reduction in viral titer on day 2 shows that T cell responses were present early in infection. In the absence of neutralizing antibodies these responses were sufficient to rescue the vaccinated mice from the onset of mortality.

We expect that a T cell vaccine would provide prophylaxis similar to what we have described against future SARS-CoV variants and strains that conserve the vaccine’s epitopes. We chose Beta (B.1.351) for our challenge study as it has a severe phenotype in a wild-type mouse background (Shuai et al., 2021). Transgenic human ACE2 (hACE2) mice exhibit encephalitis post COVID infection that does not represent human pathology (Kumari et al., 2021) and thus may be a poor model for T cell vaccine effectiveness. Instead, we chose to evaluate the HLA-A*02:01 component of our vaccine design in a HLA-A*02:01 transgenic animal model given the predominance of HLA-A02 in the human population (Ellis et al., 2000).

The MIT-T-COVID vaccine induced a response where circulating T cells migrated rapidly and efficiently into the lung upon viral challenge, as evidenced by more intense and widespread lymphocytic infiltrations than other groups. Further, the degree of CD8^+^ T cell infiltration of the lungs significantly increased in mice vaccinated with the MIT-T-COVID vaccine between days 2 and 7 post infection, and significantly decreased in mice vaccinated with the Comirnaty® vaccine. The lack of this T cell response in Comirnaty® vaccinated animals merits further study.

Another finding is that widespread congestion, hemorrhage, and few foci of thromboembolism were exclusively observed within the lungs of Comirnaty®-immunized mice. Although this finding is unrelated to the MIT-T-COVID vaccine, it suggests the possibility of immunization-induced side-effects or immunopathology of mRNA-based SARS-CoV-2 vaccines. The exact mechanism of pulmonary embolism in this case remains unknown for now and should be explored.

Cytotoxic T cells (CD8^+^ T cells) have the ability to kill virally infected cells, and thus T cell vaccines might provide long-term immunity and be effective therapeutics for Long COVID.

Post-acute sequelae of COVID-19 (PASC, “Long COVID”) causes persistent symptoms in ∼10% of people past twelve weeks of COVID infection (WHO, 2021). Long COVID has been associated with the continued presence of Spike in the blood, suggesting that a tissue based viral reservoir remains in Long COVID patients (Swank et al., 2022).

Our results suggest that if one goal of a vaccine is protection against novel viral strains it may be appropriate to develop vaccines that permit symptomatic infection but protect against severe illness. Current regulatory criteria for licensing vaccines are solely based upon the prevention of symptomatic illness for vaccine matched viral strains. Further research on T cell vaccines may reveal novel vaccine designs that prevent severe illness while providing protection against viral variants as an additional tool in the fight against global pandemics.

## Methods

### Mice

A total of 33 male HLA-A*02:01 human transgenic mice (Taconic 9659) were used that were between 12 and 16 weeks old when they received their initial vaccination. These CB6F1 background mice carry the MHC class I alleles HLA-A*02:01, H-2-Kb, H-2-Db, H-2-Kd, and H-2-Dd, and H-2-Ld. The mice carry the MHC class II alleles H-2-IAd, H-2-IEd, and H-2-IAb. We also vaccinated a female cohort of the same transgenic mouse strain (Taconic 9659, 3 mice per vaccine) for immunogenicity measurement, and results are shown in Supplemental Figures 8-11.

### Tissue culture and virus

Vero E6 cells (ATCC, CRL:1586) were grown in minimum essential medium (EMEM, Gibco) supplemented with penicillin (100 units/mL), streptomycin (100 μg/mL) and 10% fetal bovine serum (FBS). Two strains of SARS-CoV-2 were used in this study, SARS-CoV-2 (US_WA-1/2020 isolate) and Beta (B.1.351/SA, Strain: hCoV-19/USA/MD-HP01542/2021). Both viruses were propagated and quantified in Vero E6 cells and stored at -80 °C until needed.

### MIT-T-COVID vaccine design

#### MHC Class I epitopes

Eight peptides from SARS-CoV-2 were selected as a subset of the MHC class I *de novo* MIRA only vaccine design of Liu et al., 2021. We filtered this set of 36 peptides to the 8 peptides predicted to be displayed by HLA-A*02:01 by a combined MIRA and machine learning model of peptide-HLA immunogenicity (Liu et al., 2021). The combined model predicts which HLA molecule displayed a peptide that was observed to be immunogenic in a MIRA experiment, and uses machine learning predictions of peptide display for HLA alleles not observed or peptides not tested in MIRA data. Thus, all eight MHC class I peptides in our vaccine were previously observed to be immunogenic in data from convalescent COVID-19 patients (Snyder et al., 2021). We further validated that all peptides are predicted to bind HLA-A*02:01 with high (less than or equal to 50 nM) affinity using the NetMHCpan-4.1 (Reynisson et al., 2020) and MHCflurry 2.0 (O’Donnell et al., 2020) machine learning models. For inclusion in the assembled construct, the eight vaccine peptides were randomly shuffled, and alternate peptides were flanked with five additional amino acids at each terminus as originally flanked in the SARS-CoV-2 proteome. The CD4-2 vaccine epitope produced a CD8^+^ response, and this may be a consequence of a contained MHC class I epitope, with candidates including SLINTLNDL (HLA-A*02:01 predicted binding 55 nM), ASLINTLNDL (H-2-Db predicted binding 289 nM), FYTSKTTVASL (H-2-Kd predicted binding 293 nM), FYFYTSKTTV (H-2-Kd predicted binding 40 nM), and YFYTSKTTV (H-2-Kd predicted binding 56 nM).

#### MHC Class II epitopes

Three peptides from SARS-CoV-2 were optimized for predicted binding to H-2-IAb. We scored all SARS-CoV-2 peptides of length 13-25 using the sliding window approach of Liu et al. (2020) and a machine learning ensemble that outputs the mean predicted binding affinity (IC50) of NetMHCIIpan-4.0 (Reynisson et al., 2020) and PUFFIN (Zeng et al., 2019). We selected the top three peptides by predicted binding affinity using a greedy selection strategy with a minimum edit distance constraint of 5 between peptides to avoid selecting overlapping windows. All three peptides were flanked with an additional five amino acids per terminus from the SARS-CoV-2 proteome.

The start and end amino acid positions of each vaccine peptide in its origin gene is shown in Supplemental Table 1. For SARS-CoV-2, peptides are aligned to reference proteins in UniProt (UniProt Consortium, 2019) (UniProt IDs: P0DTC2 (S), P0DTC3 (ORF3a), P0DTC5 (M), P0DTC9 (N), P0DTD1 (ORF1ab)). All of the epitopes are conserved over these variants of concern: Alpha (B.1.1.7), Beta (B.1.351), Gamma (P.1), Delta (21A, 21J, B.1.617.2), Kappa (B.1.617.1), Epsilon (B.1.427, B.1.429), Iota (B.1.526), Lambda (C.37), Mu (B.1.621), Omicron (BA.1, BA.2, BA.4, B.5, BA.2.12.1, BA.2.75), EU1 (B.1.177).

### MIT-T-COVID vaccine formulation

Codon optimization for mouse expression of the MIT-T-COVID vaccine construct was performed using the IDT Codon Optimization Tool (Integrated DNA Technologies). The resulting nucleic acid sequence is provided in Supplemental Table 2. RNA was synthesized by TriLink BioTechnologies as a modified mRNA transcript with full substitution of 5-Methoxy-U, capped (Cap 1) using CleanCap® AG and polyadenylated (120A). RNA containing lipid nanoparticles were prepared as previously described (Pardi et al., 2017). Briefly, an ethanolic solution of ALC-0315 (Patent WO2017075531), cholesterol, distearoylphosphatidylcholine (DSPC), and 2-[(polyethylene glycol)-2000] N,N ditetradecylacetamide (ALC-0159, Patent Application US14732218) was rapidly mixed with an solution of RNA in citrate buffer at pH 4.0 (composition described in Patent WO2018081480). Physical properties of the LNP such as size and polydispersity were assessed by Malvern Zetasizer and encapsulation efficiency by Ribogreen assay (Life Technologies).

### Animal vaccination

Thirty-three HLA-A*02:01 human transgenic mice were randomly divided into three groups and immunized twice at three-week intervals with vehicle (PBS/300 mM sucrose), 10 μg of Comirnaty® vaccine or 10 μg of MIT-T-COVID vaccine. All vaccines were administered as a 50 μl intramuscular injection. The Comirnaty® vaccine was wastage vaccine that was diluted for human administration (0.9% NaCl diluent), and remaining unusable wastage vaccine in vials was flash frozen at -80 °C. This wastage vaccine was later thawed and immediately administered without dilution (50 μL is 10 μg of mRNA). No Comirnaty® was used that could have been administered to humans. Since Comirnaty® was thawed twice, our results may not be representative of its best performance. The MIT-T-COVID vaccine was diluted to 10 μg in 50 μL with PBS with 300 mM sucrose and then administered. In the female unchallenged cohort direct peptide vaccination was performed to test the importance of mRNA-LNP delivery. Three mice were vaccinated with an injection of 14 short synthetic peptides in the MIT-T-COVID vaccine (15 μg per peptide, 210 μg in total) adjuvanted with 50 μg of high molecular weight polyinosine-polycytidylic acid (Poly(I:C), InvivoGen, tlrl-pic) in a volume of 150 μL via intramuscular injection. These peptides include 11 MHC class I epitopes and 3 MHC class II epitopes present in the MIT mRNA vaccine (all Supplemental Table 1 epitopes except CD4-3). CD4-3 was not included during peptide/poly IC vaccination and was used as a negative control for immunogenicity measurement.

### Viral challenge

At two weeks post booster immunization, eight mice of each group were challenged with 5 × 10^4^ TCID_50_/60 μL of SARS-CoV-2 (B.1.351/SA, Strain: hCoV-19/USA/MD-HP01542/2021) via intranasal (IN) route. Mice were weighted daily and clinically observed at least once daily and scored based on a 1–4 grading system that describes the clinical wellbeing. Three mice in each group were euthanized at 2 dpi for assessing viral loads and histopathology of the lung. The remaining five mice were continued monitored for weight changes, other signs of clinical illness, and mortality (if any) for up to 7 dpi before euthanasia for assessing antibody responses within the blood and viral loads and histopathology of the lung. Animal studies were conducted at Galveston National Laboratory at University of Texas Medical Branch at Galveston, Texas, based on a protocol approved by the Institutional Animal Care and Use Committee at UTMB at Galveston.

### Assessment of mortality and morbidity

Differentially immunized and challenged mice were monitored at least once each day for the morbidity and mortality and assigned the clinical scores based on the following: 1: Healthy, 2: ruffled fur, lethargic. 3: hunched posture, orbital tightening, increased respiratory rate, and/or > 15% weight loss, and 4: dyspnea and/or cyanosis, reluctance to move when stimulated or > 20% weight loss.

### Immunogenicity measurements

At 14 days post booster immunization, three mice of each group were sacrificed for harvesting splenocytes in 2 mL R10 medium (RPMI, 10% FBS, 1%P/S, 10 mM HEPES). Briefly, spleens were homogenized and subjected to filtration onto 40 μm cell strainers, followed by a wash of strainers with 10 mL PBS and centrifuged at 500 g for 5 min at 4 °C. Cell pellet was resuspended by using 2 mL of 1x red blood cell lysis buffer (eBioscience) for 2-3 min, followed by a supplement of 20 mL PBS. Resulting cell suspensions were centrifuged at 500 g for 5 min, resuspended in 2 mL R10 medium before counting the numbers under a microscope. For a brief in vitro stimulation, aliquots of 10^6^ cells were incubated with indicated peptide at a final concentration of 1 μg/mL in each well of a 96-well plate. GolgiPlug (5 μg/mL, BD Bioscience) was added into the culture at 1 hr post stimulation and followed by an additional 4 hrs incubation. For cell surface staining, cultured splenocytes were resuspended in 40 μL FACS buffer containing fluorochrome-conjugated antibodies and incubated 1 hr at 4 °C followed by cell fixation using Cytofix/Cytoperm buffer (BD Bioscience) for 20 min at 4 °C. Cells were further incubated with fluorochrome-conjugated cytokine antibodies overnight. The next day, cells were washed twice using 1 × Perm/Wash buffer and resuspended in 300 μL FACS buffer for analysis. For Foxp3 staining, cells were fixed and permeabilized by Foxp3/Transcription Factor Staining Buffer Set (Thermo Fisher Scientific) according to the instruction. Fixable Viability Dye eFluor506 (Thermo Fisher Scientific) was also used in all sample staining to exclude dead cells from our data analysis. The fluorochrome-conjugated anti-mouse antibodies included: FITC-conjugated CD3 (17A2, Biolegend), efluor450-conjugated CD4 (GK1.5, eBioscience), PE-Cy7-conjugated CD8 (53-6.7, eBioscience), PE-conjugated IFN-γ (XMG1.2, eBioscience), PerCP-eFlour710-conjugated TNF-α (MP6-XV22, eBioscience). For the Foxp3 staining: Pacific Blue-conjugated CD4 (GK1.5), FITC-conjugated CD25 (PC61), Percp-Cy5.5-conjugated CTLA4 (UC-10-4B9), PE-conjugated Foxp3 (FJK-16s). For the CD44+ T cell analysis (female cohort only): FITC-conjugated CD3 (17A2, Biolegend), efluor450-conjugated CD4 (GK1.5, eBioscience), PE-Cy7-conjugated CD8 (53-6.7, eBioscience), APC-conjugated mouse/human CD44 (IM7, BioLegend). Cell acquisition was performed a BD LSR Fortessa and data were analyzed using BD FACSDiva 9.0 and FlowJo 10 (FlowJo, LLC). Lymphocytes were defined by SSC-A vs. FSC-A plots. Singlet cells were defined by FSC-H vs. FSC-A plots. Dead cells were excluded by positive staining with viability dye. CD4^+^ and CD8^+^ T cells were gated from CD3^+^ cells. The cytokines secreting CD4^+^ and CD8^+^ T cells were then identified with IFN-γ and TNF-α expression. Boundaries between positive and negative cells for the given marker were defined by the fluorescence minus one (FMO) control and adjusted according to the unstimulated splenocyte group. For the Treg cell gating strategy, CD4^+^ T cells were gated from CD3^+^ cells and identified by the positivity of Foxp3 and CD25 staining.

### Viral titer assay

For virus quantitation, the frozen lung specimens were weighed before homogenization in PBS/2% FBS solution using the TissueLyser (Qiagen), as previously described (Tseng et al., 2007). The homogenates were centrifuged to remove cellular debris. Cell debris-free homogenates were used to quantifying infectious viruses in the standard Vero E6 cell-based infectivity assays in 96-well microtiter plates, as we routinely used in the lab (Tseng et al., 2012). Titers of virus were expressed as 50% tissue culture infectious dose per gram of tissue (TCID_50_/g).

### Antibody neutralization assay

Sera of mice collected at 7 dpi were used for measuring specific antibody responses. Briefly, sera were heat-inactivated (56 °C) for 30 min, were stored at -80 °C until needed. For determining the SARS-CoV-2 neutralizing antibody titers, serially two-fold (starting from 1:40) and duplicate dilutions of heat-inactivated sera were incubated with 100 TCID of SARS-CoV-2 (US_WA-1/2020 isolate) or Beta (B.1.351/SA, Strain: hCoV-19/USA/MD-HP01542/2021) at 37 °C for 1 h before transferring into designated wells of confluent Vero E6 cells grown in 96-well microtiter plates. Vero E6 cells cultured with medium with or without virus were included as positive and negative controls, respectively. After incubation at 37 °C for 3 days, individual wells were observed under the microscopy for the status of virus-induced formation of cytopathic effect. The 100% neutralizing titers (NT_100_) of sera were expressed as the lowest dilution folds capable of completely preventing the formation of viral infection-induced cytopathic effect in 100% of the wells.

### Serum IgG/IgM response by ELISA

ELISA was applied to verified serum IgG and IgM responses with SARS-CoV-2 (US_WA-1/2020 isolate) infected Vero-E6 cell lysate. In brief, SARS-CoV-2 infected Vero-E6 cell lysate was coated on 96-well plate (Corning) at 1 μg/well in PBS for overnight at 4 °C. The plates were blocked using 1% BSA/PBST for 1 h at room temperature. The 5-fold serial-dilution serum from each mouse (starting at 1:100) was then added into antigen-coated plates and incubated for 1 h at 37 °C. The plates were then washed three times with PBST (PBS/0.1% Tween-20) followed by incubation with 100 μL of anti-mouse IgG and IgM HRP conjugated secondary antibody (Jackson immunoresearch) (1:2000) for 1 h at 37 °C. After three times wash using PBST, 100 μL of ABTS substrate (Seracare) was added to the plates and incubated for 30 min in the dark. After stopped by adding 1% SDS. Then absorbance at 405 nm (OD 405 nm) was measured with the plate reader (Molecular Devices) and analyzed with GraphPad Prism Version 9.1.2.

### RNA extraction and quantitative RT-PCR

Lung tissues were weighted and homogenized in 1 mL of Trizol reagent (Invitrogen) using TissueLyser (Qiagen). The RNA was then extracted using Direct-zol RNA miniprep kits (Zymo research) according to the manufacturer’s instructions. 500 ng total RNA was then applied to cDNA synthesis using iScript cDNA Synthesis kit (Biorad) according to the manufacturer’s instructions. The viral genomic RNA, subgenomic RNA and mice 18s rRNA has then been amplified using iQ SYBR green supermix (Biorad) and performed using CFX96 real time system (Biorad). The samples were run in duplicate using the following conditions: 95 °C for 3 min then 45 cycles of 95 °C for 15 s and 58 °C for 30 s. The level of expression was then normalized with 18s rRNA and calculated using 2^-Δ*Ct*^ method, as we have previously described (Tseng et al., 2007; Agrawal et al., 2015)

The primer set for SARS-CoV-2 RNA amplification is nCoV-F (ACAGGTACGTTAATAGTTAATAGCGT) and nCoV-R (ATATTGCAGCAGTACGCACACA). SgLeadSARS2-F (CGATCTCTTGTAGATCTGTTCTC) and nCoV-R (ATATTGCAGCAGTACGCACACA) were used for subgenomic SARS-CoV-2 RNA amplification. Mouse 18s rRNA was served as the internal control and amplified using 18s-F(GGACCAGAGCGAAAGCATTTGCC) and 18s-R (TCAATCTCGGGTGGCTGAACGC).

### Immunohistochemistry

All slides were prepared by the Histopathology Core (UTMB) into 5 μm paraffin-embedded sections for immunohistochemistry (IHC). IHC staining and analysis were performed by UTMB according to previously published protocols (Tseng et al., 2007; Yoshikawa et al., 2009). In brief, a standard IHC sequential incubation staining protocol was followed to detect the SARS-CoV-2 spike (S) protein, CD4^+^ cells, or CD8^+^ cells using a rabbit-raised anti-SARS-CoV-2 S protein antibody (1:5000 dilution, ab272504, Abcam plc, Cambridge UK), a rabbit monoclonal anti-CD4 antibody (1:250 dilution, ab183685, ab183685, Abcam plc, Cambridge, UK), and a rabbit monoclonal anti-CD8 antibody (1:500 dilution, ab217344, Abcam plc, Cambridge, UK) followed by peroxidase-conjugated secondary antibody and 3,3’-Diaminobenzidine (DAB) substrate kit (MP-7802, Vector Laboratories, Burlingame, CA). Slides were counterstained with hematoxylin (MHS16-500ML, Sigma-Aldrich Inc., St. Louis, MO) and antigen expression was examined under 10X and 40X magnifications using an Olympus IX71 microscope.

CD4^+^ and CD8^+^ cell counts were quantified using CellProfiler 4.2.4 (Stirling et al., 2021). The image analysis pipeline included the following modules: (1) UnmixColors (for each of DAB and hematoxylin stains), (2) ImageMath (applied to DAB stain images only; subtract 0.5 from all intensity values and set values less than 0 equal to 0), and (3) IdentifyPrimaryObjects (require object diameter 20-60 pixels [DAB stain] or 15-60 pixels [hematoxylin stain]; Global threshold strategy; Otsu thresholding method with two-class thresholding, threshold smoothing scale 1.3488, threshold correction factor 1.0, lower threshold bound 0.0, upper threshold bound 1.0, no log transform before thresholding; Shape method to distinguish clumped objects and draw dividing lines between clumped objects; automatically calculate size of smoothing filter for declumping; automatically calculate minimum allowed distance between local maxima; speed up by using lower-resolution image to find local maxima; fill holes in identified objects after declumping only; continue handling objects if excessive number of objects identified).

### Statistical Analysis

One-way ANOVA tests were performed in Python using the SciPy package (Virtanen et al., 2020). Two-way ANOVA and Tukey’s tests were performed in Python using the statsmodels package (Seabold and Perktold, 2010). Logrank tests were performed using GraphPad Prism.

## Acknowledgements

This work was supported in part by Schmidt Futures and a C3.ai grant to DG. We benefited from thoughtful comments from Michael Birnbaum.

## Supplemental Data

**Supplemental Table 1.**
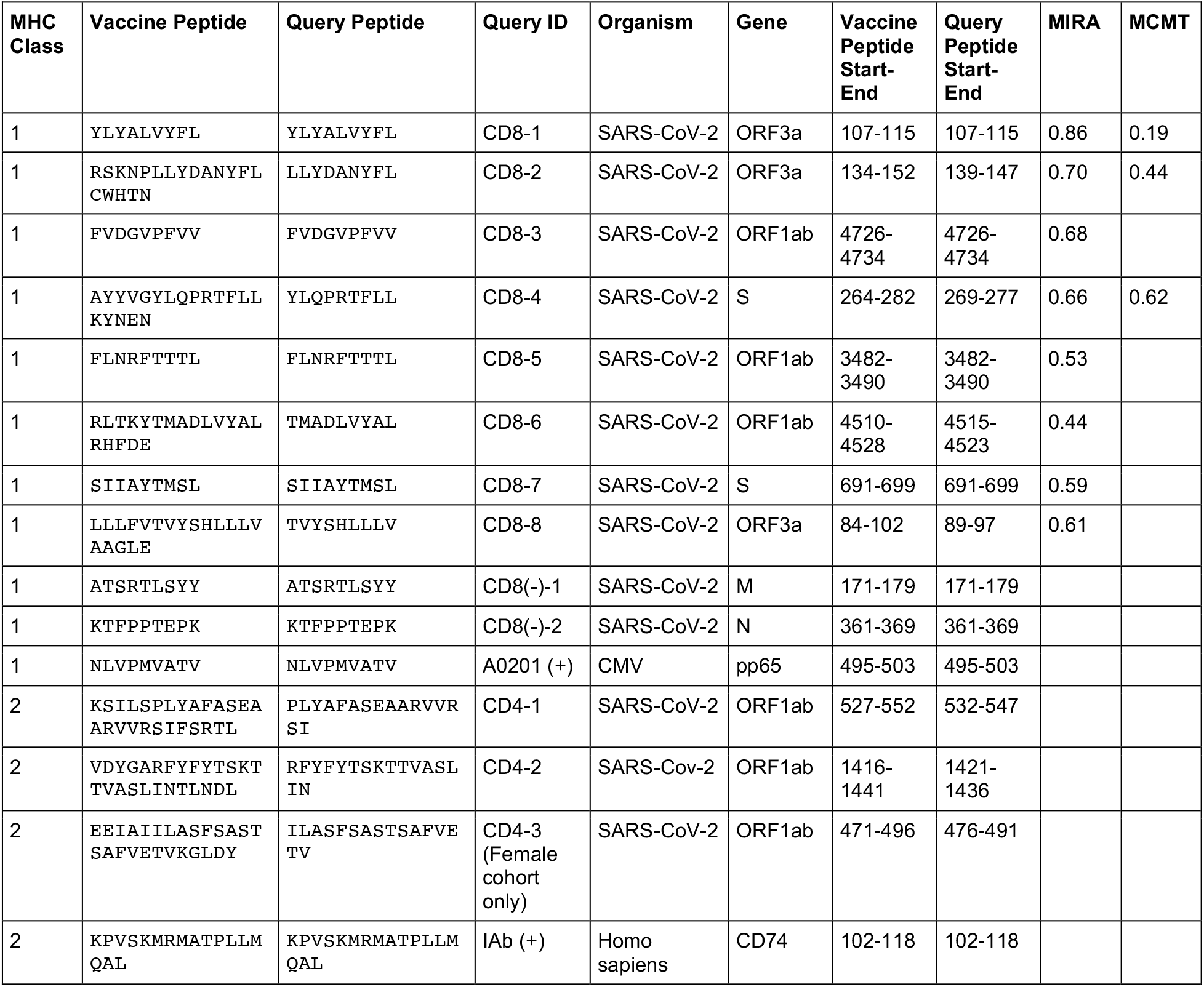
MIT-T-COVID vaccine peptides, query peptides (vaccine immunogenicity), peptide origin, and appearance probability in HLA-A*02:01 convalescent COVID-19 patients in Snyder et al. (2020) (MIRA column) and Kared et al. (2021) (MCMT). Selected peptides were tested by other studies for their immunogenicity in convalescent COVID-19 patients whose HLA type included HLA-A*02:01. The study by Snyder et al. (2020) included 80 HLA-A*02:01 convalescent COVID-19 patients and tested peptides individually or in small pools with the Multiplex Identification of T-cell Receptor Antigen Specificity (MIRA) assay. Query peptides were first filtered to only consider those with predicted HLA-A*02:01 binding affinity less than or equal to 25 nM. The MIRA fraction in Supplemental Table 1 is the number of individuals positive for a pool containing a query peptide divided by 80. The Kared et al. (2021) study evaluated 16 HLA-A*02:01 convalescent COVID-19 patient by mass cytometry– based multiplexed tetramer (MCMT) staining. The MCMT fraction in Supplemental Table 1 is the number of individuals positive for a query peptide divided by 16.

**Supplemental Table 2.**
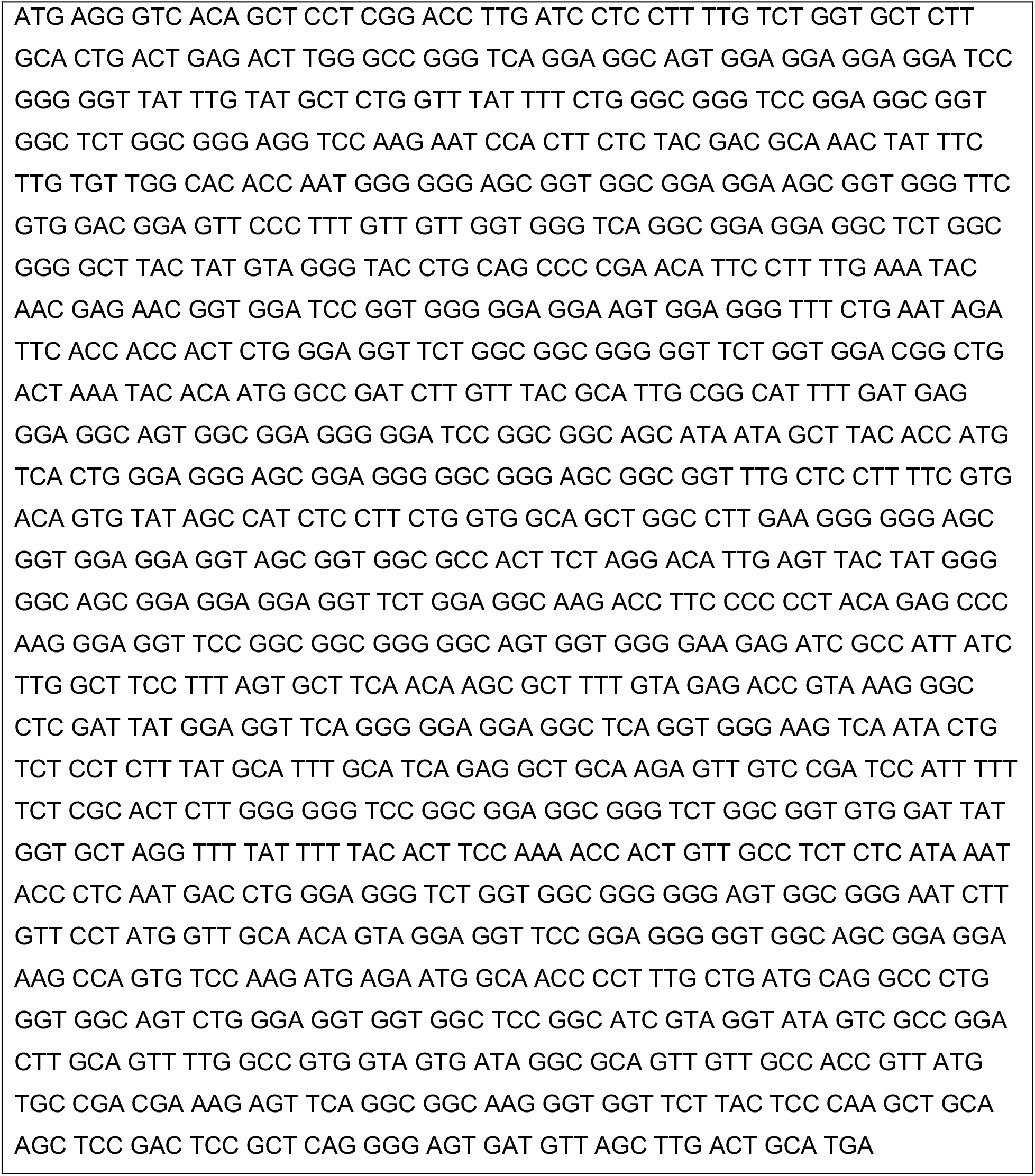
Nucleic acid sequence for assembled vaccine construct of Figure 1A.

**Supplemental Figure 1.**
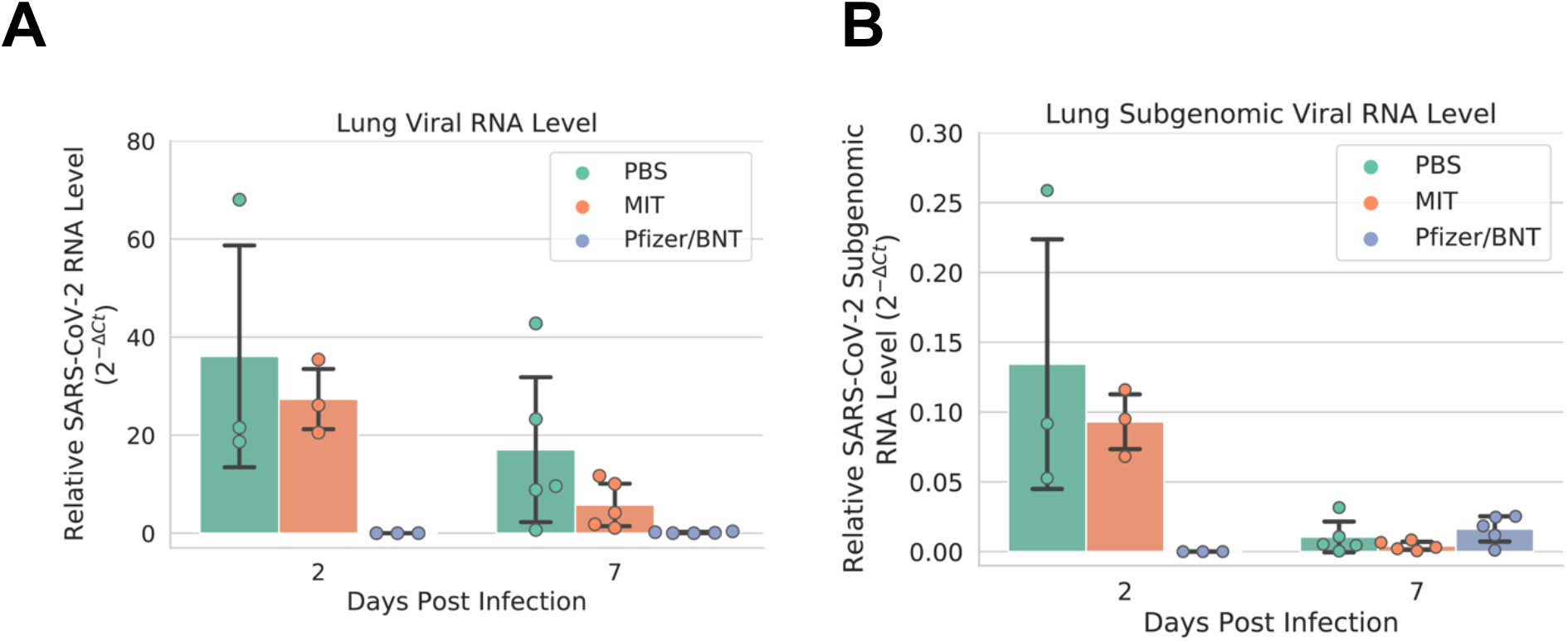
(A) Lung viral RNA level, and (B) lung subgenomic viral RNA level. Error bars indicate the standard deviation around each mean.

**Supplemental Figure 2.**
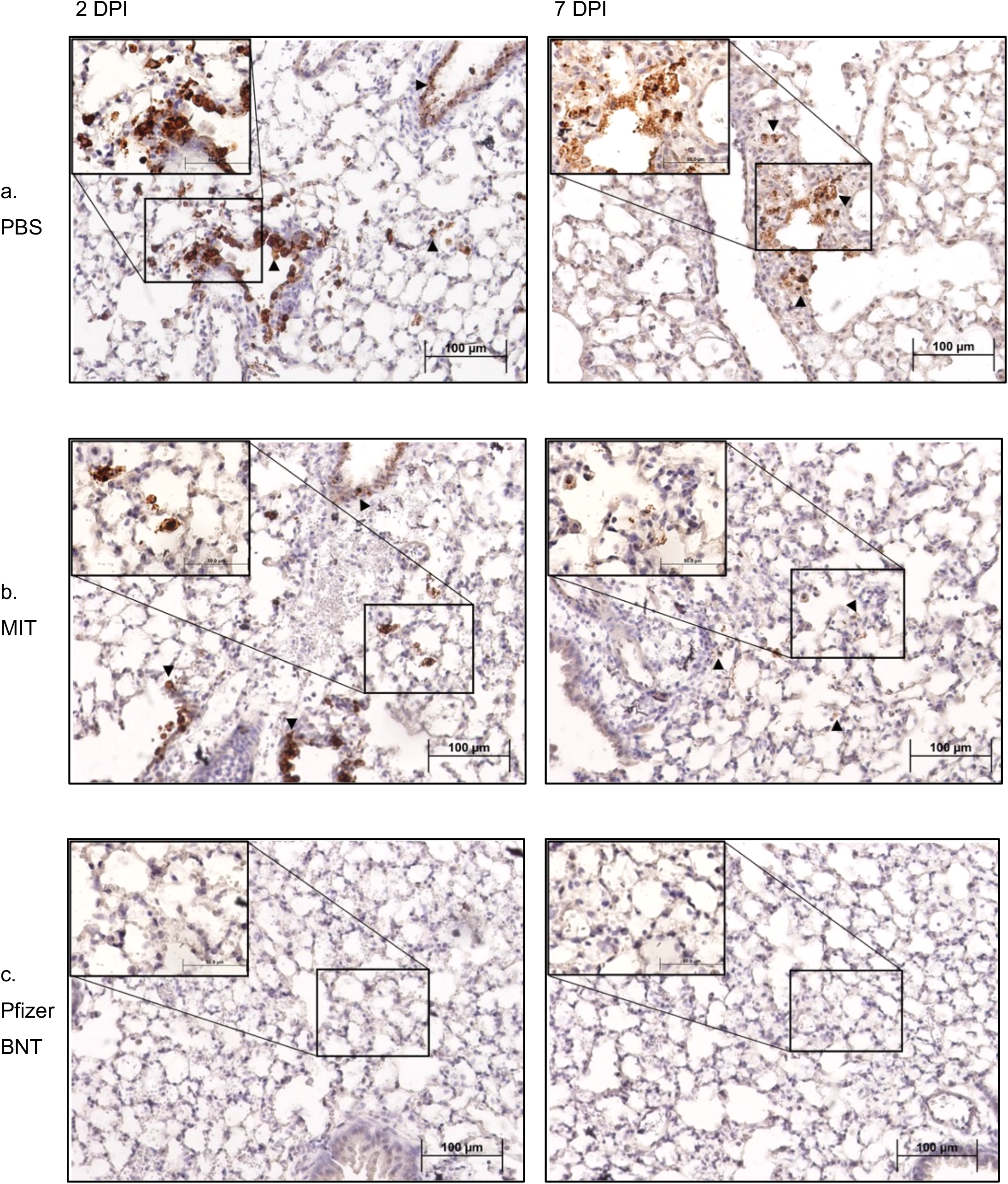
All lung samples were subjected to IHC staining for SARS-CoV-2 spike protein (brown) with hematoxylin counterstain (blue), representative images of which are shown here. Inset images taken at 40X magnification. Black arrowheads indicate selected areas of viral infection. (a) Specimens vaccinated with PBS exhibited extensive staining indicative of viral infection throughout the epithelium of both the bronchioles and the alveolar sacs, with the viral infection appearing more intense at 2 days post-infection (dpi, left) than at 7 dpi (right). Although viral infection is significantly reduced by 7 dpi, viral antigen was still readily detectable throughout alveoli. (b) In comparison, specimens vaccinated with the MIT-T-COVID vaccine exhibited similarly extensive viral infection at 2 dpi (left) throughout the bronchiolar and alveolar epithelia, albeit somewhat reduced in intensity. However, by 7 dpi (right), viral infection was significantly reduced in both extent and intensity, with brown puncta being detected only in a few alveoli scattered throughout the tissue. (c) Contrasted with both PBS and MIT-T-COVID vaccinated specimens, the Pfizer/BNT-vaccinated specimens exhibited significantly reduced viral infections at both 2 (left) and 7 dpi (right). With the exception of a single area at 7 dpi (see Supplemental Figure 3), viral antigen was undetected at both timepoints.

**Supplemental Figure 3.**
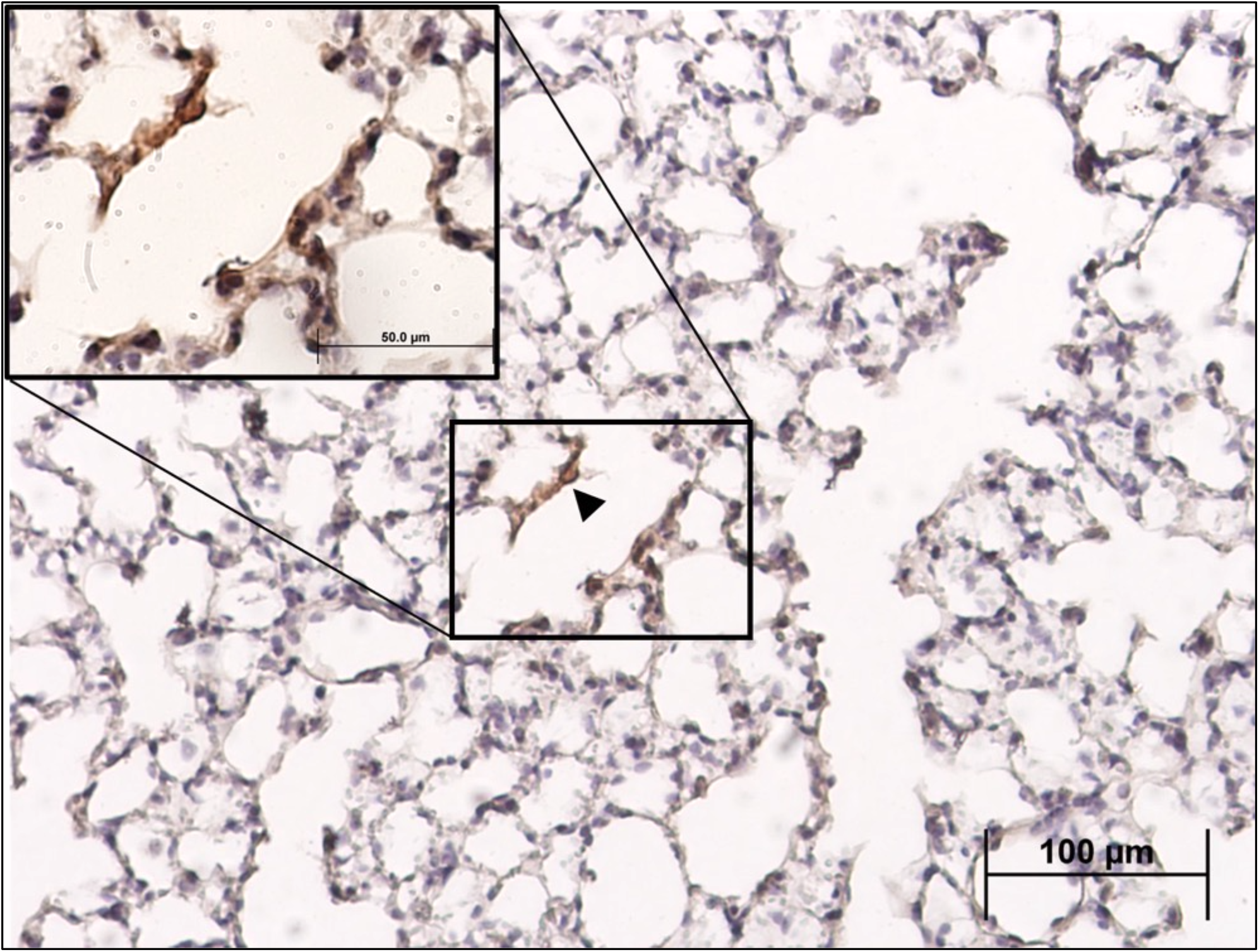
Sole region of detectable SARS-CoV-2 spike antigen in alveoli of Comirnaty® vaccinated lung specimens at 7 days post-infection. Black arrowhead indicates infected cell.

**Supplemental Figure 4.**
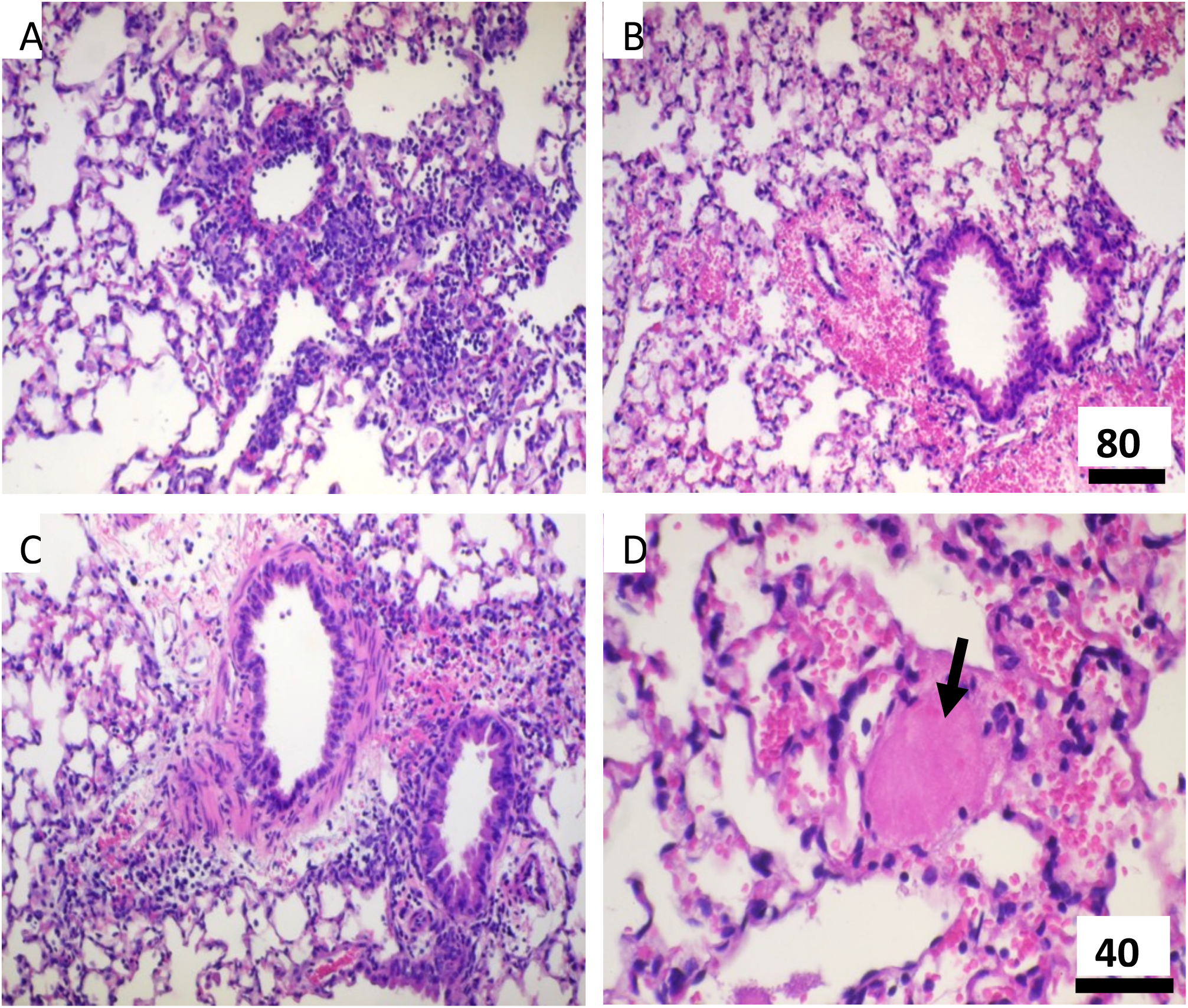
Lung histopathology. Lungs of mice vaccinated with MIT-T-COVID (A) or Pfizer/BNT (B) are compared with those with PBS (C). At 7 dpi, the MIT-T-COVID vaccinated group showed extensive lymphocytic infiltrations in perivascular regions and spaces around bronchi, bronchioles, and alveoli. There are fewer infiltrations found in the Pfizer/BNT or PBS groups and they are only localized at perivascular regions around bronchi and large bronchioles. There is widespread congestion along with hemorrhage and few foci of thromboembolism (arrow in D) seen in the Pfizer/BNT group but not others. Bar = 80 μm in A, B and C; Bar = 40 μm in D.

**Supplemental Figure 5.**
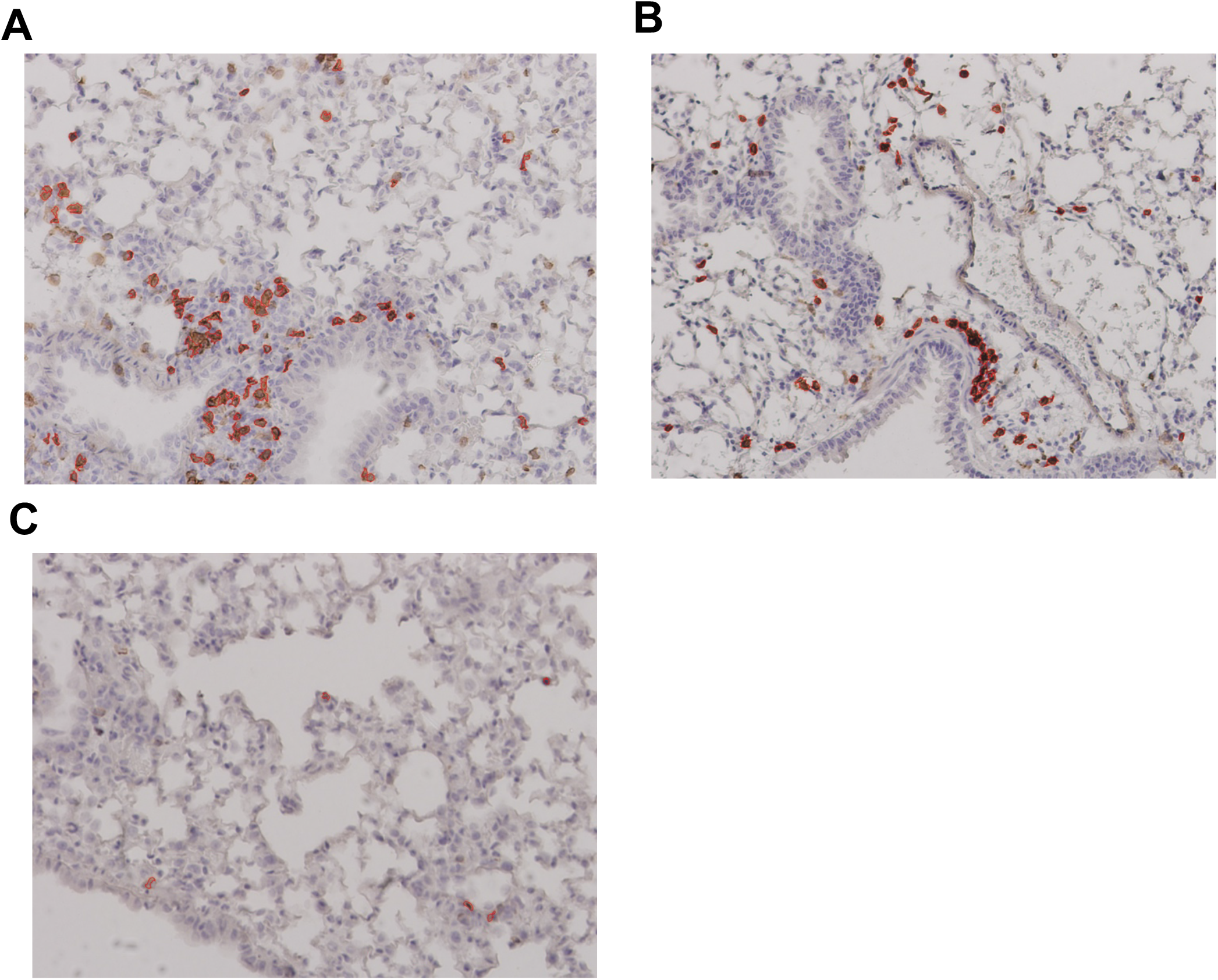
Lung immunohistochemistry for CD4^+^ cells at 7 dpi. Example CD4^+^ stain images for (A) MIT-T-COVID, (B) Pfizer/BNT, and (C) PBS vaccinated animals. Lung samples were subjected to IHC staining for CD4 (brown) with hematoxylin counterstain (blue). Images were taken at 10X magnification. Red outlines indicate CD4^+^ cells identified and counted by CellProfiler software (Methods). See also Figure 4 and Supplemental Figures 6-7.

**Supplemental Figure 6.**
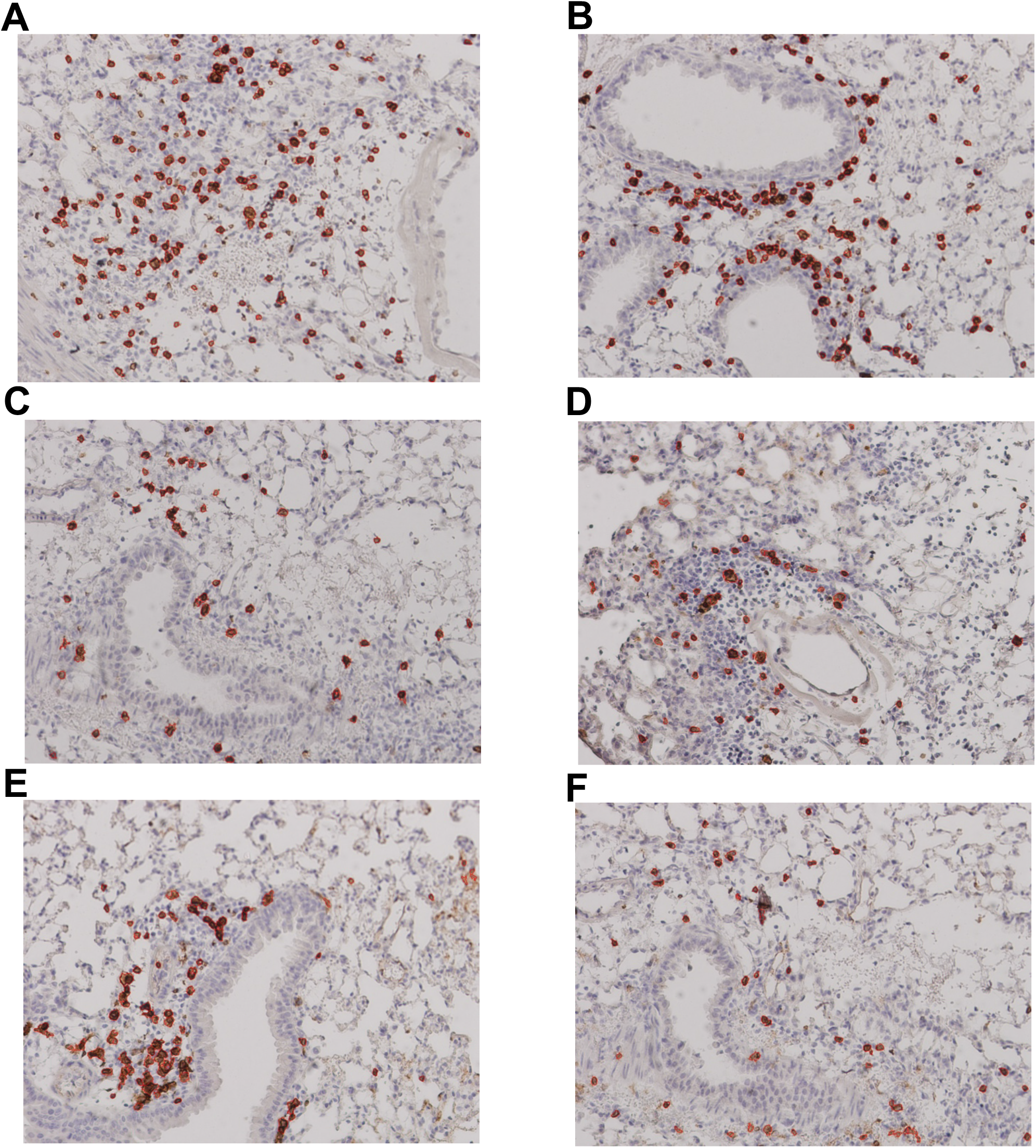
Lung immunohistochemistry for CD8^+^ and CD4^+^ cells at 2 dpi. Example CD8^+^ stain images for (A) MIT-T-COVID, (B) Pfizer/BNT, and (C) PBS vaccinated animals. Example CD4^+^ stain images for (D) MIT-T-COVID, (E) Pfizer/BNT, and (F) PBS vaccinated animals. Lung samples were subjected to IHC staining for CD4 (brown) with hematoxylin counterstain (blue). Images were taken at 10X magnification. Red outlines indicate cells identified and counted by CellProfiler software (Methods). See also Figure 4, Supplemental Figure 5, and Supplemental Figure 7.

**Supplemental Figure 7.** Complete set of cell counts in lung immunohistochemistry images for CD8^+^ and CD4^+^ cells at 2 and 7 dpi. Lung samples were subjected to IHC staining for CD4 or CD8 (brown) with hematoxylin counterstain (blue). Images were taken at 10X magnification. Red outlines indicate cells identified by CellProfiler software (Methods). Mouse and image field replicates are identified as Mouse # - Field #. See also Figure 4 and Supplemental Figures 5-6.

**Supplemental Figure 8.**
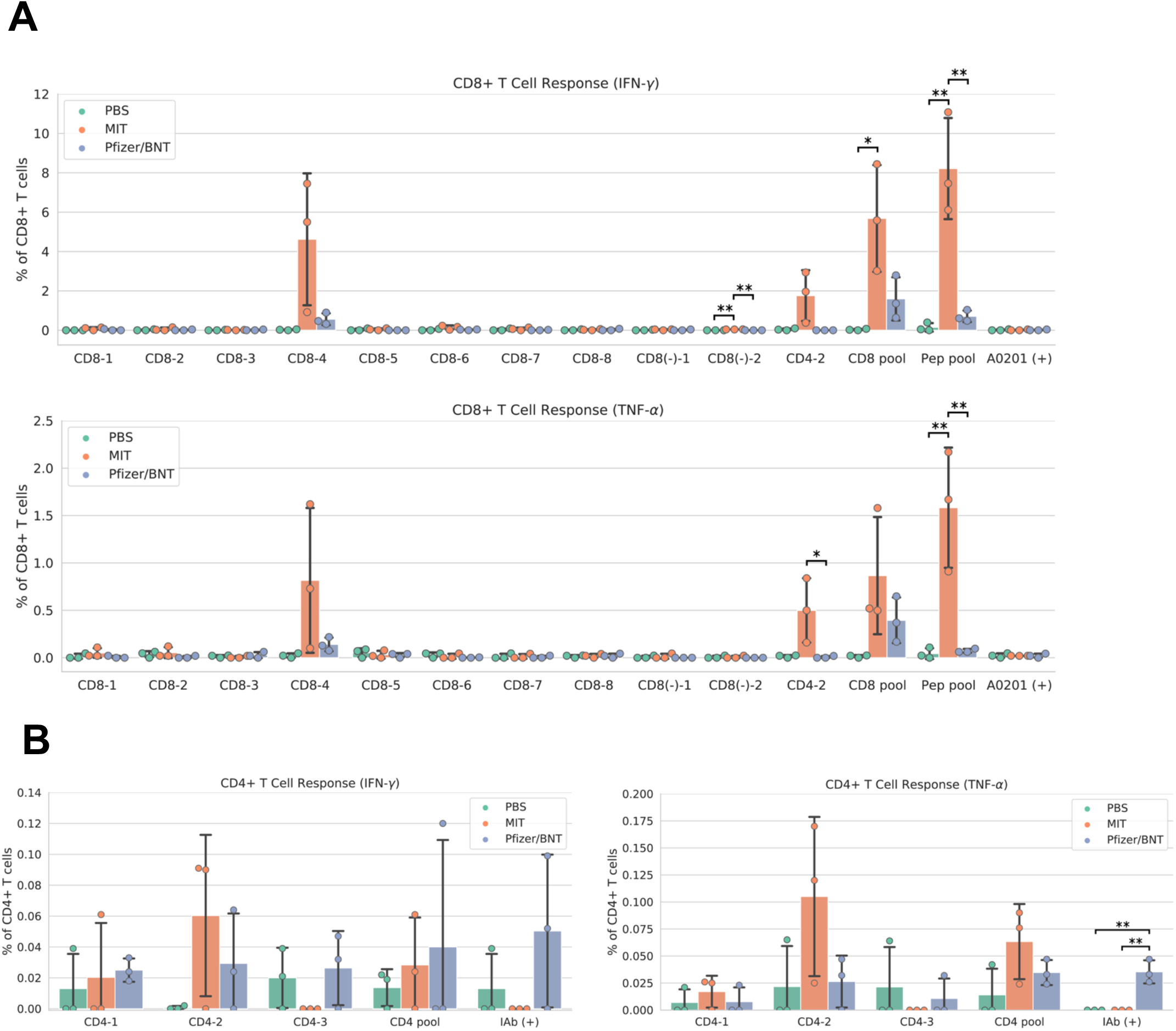
Vaccine immunogenicity in female mouse cohort (Methods). (A) CD8^+^ T cell responses, (B) CD4^+^ T cell responses. The CD8 pool includes MHC class I peptides CD8-1—CD8-8 (Supplemental Table 1). The CD4 pool includes MHC class II peptides CD4-1, CD4-2, and CD4-3. The Pep pool includes all query peptides in Supplemental Table 1. Error bars indicate the standard deviation around each mean. *P* values were computed by one-way ANOVA with Tukey’s test. \**P* < 0.05, \*\**P* < 0.01. See also Supplemental Figure 9.

**Supplemental Figure 9.**
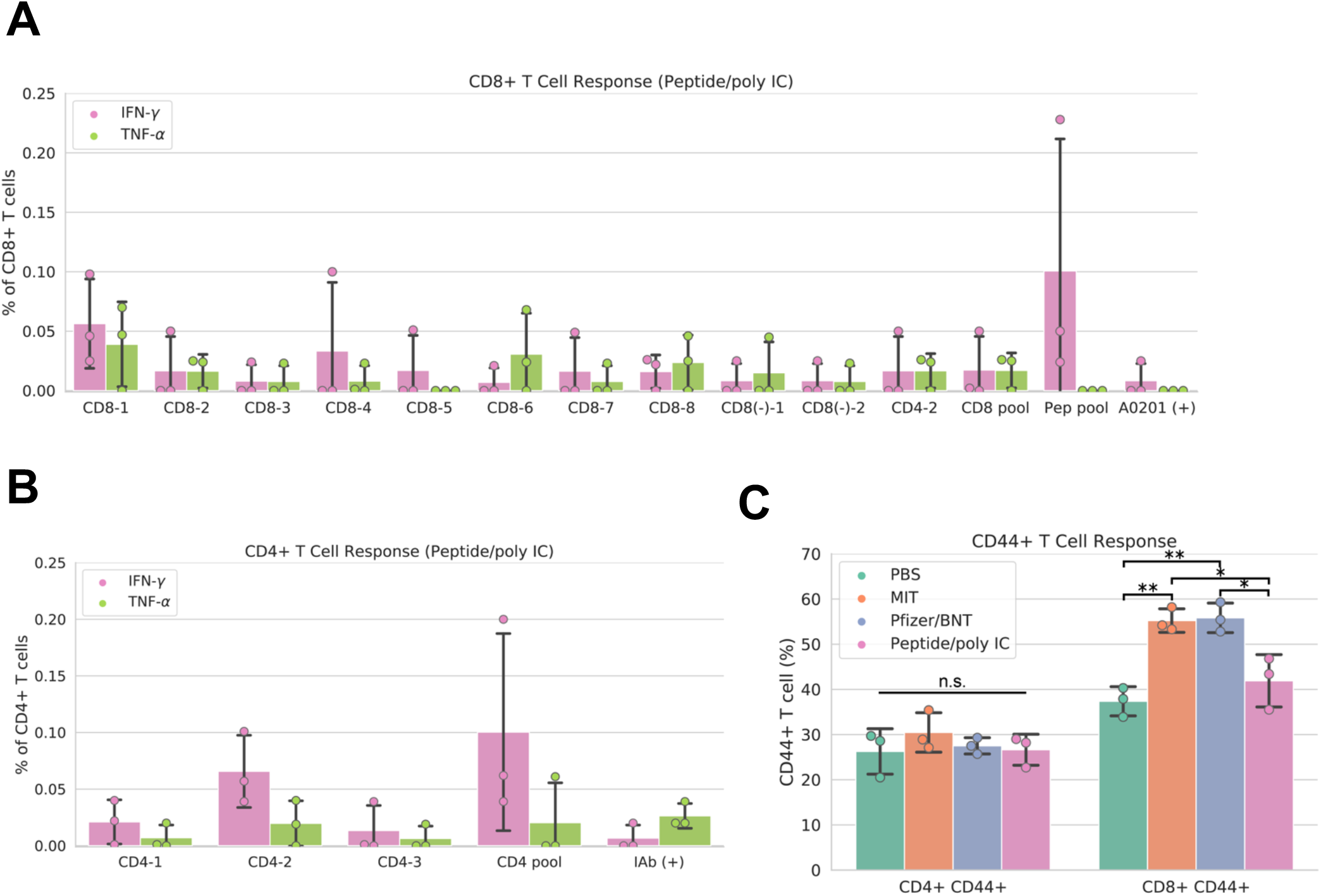
Immunogenicity of Peptide/poly IC vaccination in female mouse cohort (Methods). (A) CD8^+^ T cell responses, (B) CD4^+^ T cell responses, and (C) CD44^+^ T cell responses. The CD8 pool includes MHC class I peptides CD8-1—CD8-8 (Supplemental Table 1). Mice were vaccinated with all Supplemental Table 1 epitopes except CD4-3 (negative control). The CD4 pool includes MHC class II peptides CD4-1, CD4-2, and CD4-3. The Pep pool includes all query peptides in Supplemental Table 1. Error bars indicate the standard deviation around each mean. See also Supplemental Figure 8.

**Supplemental Figure 10.**
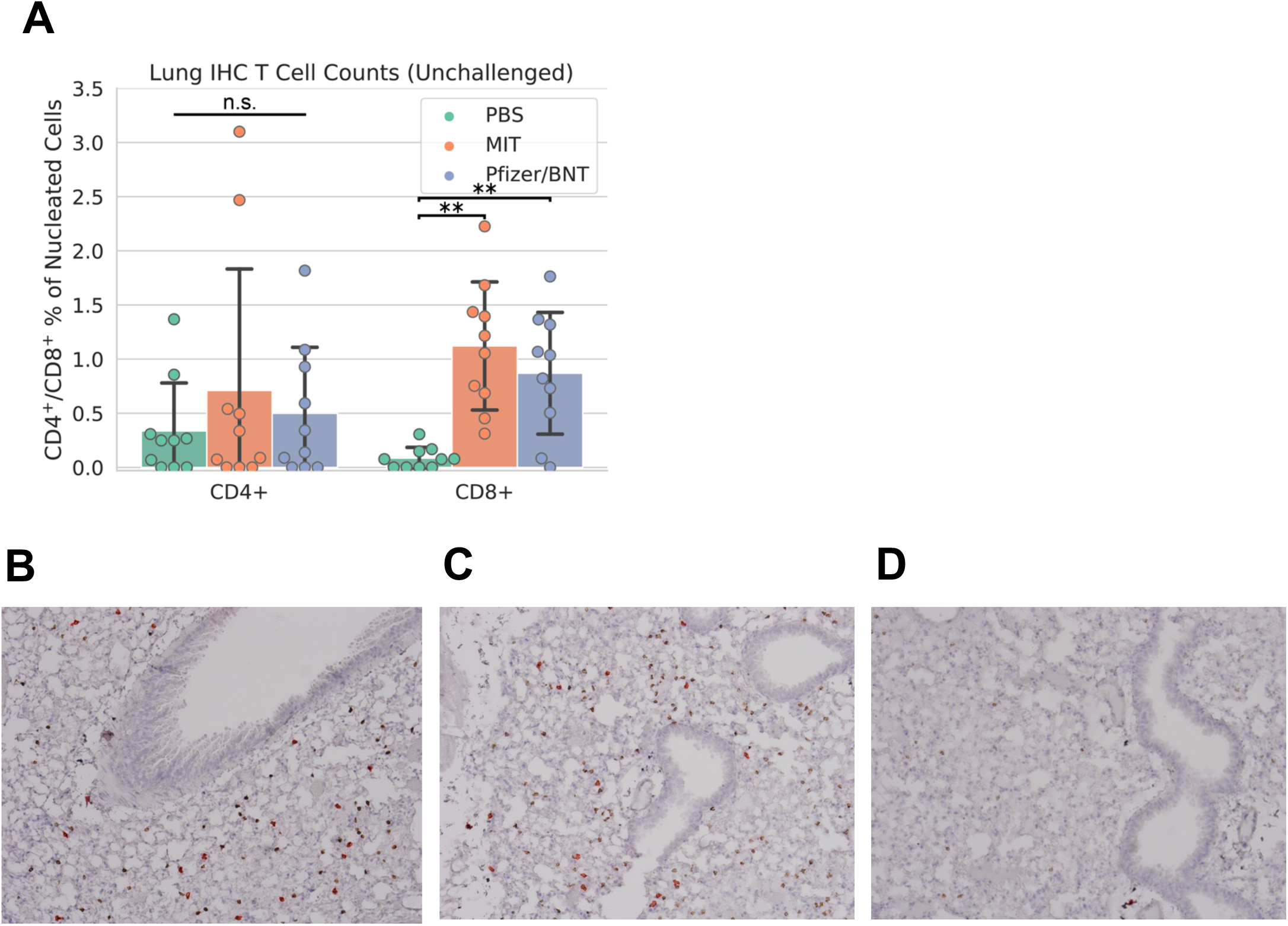
Lung immunohistochemistry for CD8^+^ and CD4^+^ cells in unchallenged female mouse cohort. (A) Counts of CD8^+^ and CD4^+^ T cells expressed as a percentage of all nucleated cells visible in each field from lung tissue. Example CD8^+^ stain images for (B) MIT-T-COVID, (C) Pfizer/BNT, and (D) PBS vaccinated animals. Lung samples were subjected to IHC staining for CD8 (brown) with hematoxylin counterstain (blue). Images were taken at 10x magnification. Red outlines in (B)-(D) indicate CD8^+^ cells identified and counted by CellProfiler software (Methods). Error bars indicate the standard deviation around each mean. *P* values were computed by one-way ANOVA with Tukey’s test. \*\**P* < 0.01, n.s. = not significant. See also Supplemental Figure 11.

**Supplemental Figure 11.** Complete set of cell counts in lung immunohistochemistry images for CD8^+^ and CD4^+^ cells from the unchallenged female mouse cohort. Lung samples were subjected to IHC staining for CD4 or CD8 (brown) with hematoxylin counterstain (blue). Images were taken at 10X magnification. Red outlines indicate cells identified by CellProfiler software (Methods). Mouse and image field replicates are identified as Mouse # - Field #. See also Supplemental Figure 10.

## References

1. Agrawal AS, Garron T, Tao X, Peng BH, Wakamiya M, Chan TS, Couch RB, Tseng CT. Generation of a transgenic mouse model of Middle East respiratory syndrome coronavirus infection and disease. J Virol. 2015 Apr;89(7):3659–70. doi: 10.1128/JVI.03427-14. Epub 2015 Jan 14. PMID: 25589660; PMCID: PMC4403411.

2. Baden LR, El Sahly HM, Essink B, Kotloff K, Frey S, Novak R, Diemert D, Spector SA, Rouphael N, Creech CB, McGettigan J, Khetan S, Segall N, Solis J, Brosz A, Fierro C, Schwartz H, Neuzil K, Corey L, Gilbert P, Janes H, Follmann D, Marovich M, Mascola J, Polakowski L, Ledgerwood J, Graham BS, Bennett H, Pajon R, Knightly C, Leav B, Deng W, Zhou H, Han S, Ivarsson M, Miller J, Zaks T; COVE Study Group. Efficacy and Safety of the mRNA-1273 SARS-CoV-2 Vaccine. N Engl J Med. 2021 Feb 4;384(5):403–416. doi: 10.1056/NEJMoa2035389. Epub 2020 Dec 30. PMID: 33378609; PMCID: PMC7787219.

3. Benjamini O, Rokach L, Itchaki G, Braester A, Shvidel L, Goldschmidt N, Shapira S, Dally N, Avigdor A, Rahav G, Lustig Y, Ben David SS, Fineman R, Paz A, Bairey O, Polliack A, Levy I, Tadmor T. Safety and efficacy of the BNT162b mRNA COVID-19 vaccine in patients with chronic lymphocytic leukemia. Haematologica. 2022 Mar 1;107(3):625–634. doi: 10.3324/haematol.2021.279196. PMID: 34320789; PMCID: PMC8883569.

4. Cohen AA, van Doremalen N, Greaney AJ, Andersen H, Sharma A, Starr TN, Keeffe JR, Fan C, Schulz JE, Gnanapragasam PNP, Kakutani LM, West AP Jr, Saturday G, Lee YE, Gao H, Jette CA, Lewis MG, Tan TK, Townsend AR, Bloom JD, Munster VJ, Bjorkman PJ. Mosaic RBD nanoparticles protect against challenge by diverse sarbecoviruses in animal models. Science. 2022 Aug 5;377(6606):eabq0839. doi: 10.1126/science.abq0839. Epub 2022 Aug 5. PMID: 35857620; PMCID: PMC9273039.

5. Croft NP, Smith SA, Pickering J, Sidney J, Peters B, Faridi P, Witney MJ, Sebastian P, Flesch IEA, Heading SL, Sette A, La Gruta NL, Purcell AW, Tscharke DC. Most viral peptides displayed by class I MHC on infected cells are immunogenic. Proc Natl Acad Sci U S A. 2019 Feb 19;116(8):3112–3117. doi: 10.1073/pnas.1815239116. Epub 2019 Feb 4. PMID: 30718433; PMCID: PMC6386720.

6. de Silva TI, Liu G, Lindsey BB, Dong D, Moore SC, Hsu NS, Shah D, Wellington D, Mentzer AJ, Angyal A, Brown R, Parker MD, Ying Z, Yao X, Turtle L, Dunachie S; COVID-19 Genomics UK (COG-UK) Consortium, Maini MK, Ogg G, Knight JC; ISARIC4C Investigators, Peng Y, Rowland-Jones SL, Dong T. The impact of viral mutations on recognition by SARS-CoV-2 specific T cells. iScience. 2021 Nov 19;24(11):103353. doi: 10.1016/j.isci.2021.103353. Epub 2021 Oct 28. PMID: 34729465; PMCID: PMC8552693.

7. Ellis JM, Henson V, Slack R, Ng J, Hartzman RJ, Katovich Hurley C. Frequencies of HLA-A2 alleles in five U.S. population groups. Predominance Of A*02011 and identification of HLA-A*0231. um Immunol. 2000 Mar;61(3):334–40. doi: 10.1016/s0198-8859(99)00155-x. PMID: 10689125.

8. Geers D, Shamier MC, Bogers S, den Hartog G, Gommers L, Nieuwkoop NN, Schmitz KS, Rijsbergen LC, van Osch JAT, Dijkhuizen E, Smits G, Comvalius A, van Mourik D, Caniels TG, van Gils MJ, Sanders RW, Oude Munnink BB, Molenkamp R, de Jager HJ, Haagmans BL, de Swart RL, Koopmans MPG, van Binnendijk RS, de Vries RD, GeurtsvanKessel CH. SARS-CoV-2 variants of concern partially escape humoral but not T-cell responses in COVID-19 convalescent donors and vaccinees. Sci Immunol. 2021 May 25;6(59):eabj1750. doi: 10.1126/sciimmunol.abj1750. PMID: 34035118; PMCID: PMC9268159.

9. Hajnik RL, Plante JA, Liang Y, Alameh MG, Tang J, Bonam SR, Zhong C, Adam A, Scharton D, Rafael GH, Liu Y, Hazell NC, Sun J, Soong L, Shi PY, Wang T, Walker DH, Sun J, Weissman D, Weaver SC, Plante KS, Hu H. Dual spike and nucleocapsid mRNA vaccination confer protection against SARS-CoV-2 Omicron and Delta variants in preclinical models. Sci Transl Med. 2022 Sep 14;14(662):eabq1945. doi: 10.1126/scitranslmed.abq1945. Epub 2022 Sep 14. PMID: 36103514.

10. Heitmann JS, Bilich T, Tandler C, Nelde A, Maringer Y, Marconato M, Reusch J, Jäger S, Denk M, Richter M, Anton L, Weber LM, Roerden M, Bauer J, Rieth J, Wacker M, Hörber S, Peter A, Meisner C, Fischer I, Löffler MW, Karbach J, Jäger E, Klein R, Rammensee HG, Salih HR, Walz JS. A COVID-19 peptide vaccine for the induction of SARS-CoV-2 T cell immunity. Nature. 2022 Jan;601(7894):617–622. doi: 10.1038/s41586-021-04232-5. Epub 2021 Nov 23. PMID: 34814158; PMCID: PMC8791831.

11. Israelow B, Mao T, Klein J, Song E, Menasche B, Omer SB, Iwasaki A. Adaptive immune determinants of viral clearance and protection in mouse models of SARS-CoV-2. Sci Immunol. 2021 Oct 15;6(64):eabl4509. doi: 10.1126/sciimmunol.abl4509. Epub 2021 Sep 2. PMID: 34623900; PMCID: PMC9047536.

12. Kared H, Redd AD, Bloch EM, Bonny TS, Sumatoh H, Kairi F, Carbajo D, Abel B, Newell EW, Bettinotti MP, Benner SE, Patel EU, Littlefield K, Laeyendecker O, Shoham S, Sullivan D, Casadevall A, Pekosz A, Nardin A, Fehlings M, Tobian AA, Quinn TC. SARS-CoV-2-specific CD8+ T cell responses in convalescent COVID-19 individuals. J Clin Invest. 2021 Mar 1;131(5):e145476. doi: 10.1172/JCI145476. PMID: 33427749; PMCID: PMC7919723.

13. Kingstad-Bakke B, Lee W, Chandrasekar SS, Gasper DJ, Salas-Quinchucua C, Cleven T, Sullivan JA, Talaat A, Osorio JE, Suresh M. Vaccine-induced systemic and mucosal T cell immunity to SARS-CoV-2 viral variants. Proc Natl Acad Sci U S A. 2022 May 17;119(20):e2118312119. doi: 10.1073/pnas.2118312119. Epub 2022 May 13. PMID: 35561224; PMCID: PMC9171754.

14. Kotturi MF, Assarsson E, Peters B, Grey H, Oseroff C, Pasquetto V, Sette A. Of mice and humans: how good are HLA transgenic mice as a model of human immune responses? Immunome Res. 2009 Jun 17;5:3. doi: 10.1186/1745-7580-5-3. PMID: 19534819; PMCID: PMC2702351.

15. Kreiter S, Selmi A, Diken M, Sebastian M, Osterloh P, Schild H, Huber C, Türeci O, Sahin U. Increased antigen presentation efficiency by coupling antigens to MHC class I trafficking signals. J Immunol. 2008 Jan 1;180(1):309–18. doi: 10.4049/jimmunol.180.1.309. Erratum in: J Immunol. 2012 Sep 1;189(5):2682. PMID: 18097032.

16. Kumari P, Rothan HA, Natekar JP, Stone S, Pathak H, Strate PG, Arora K, Brinton MA, Kumar M. Neuroinvasion and Encephalitis Following Intranasal Inoculation of SARS-CoV-2 in K18-hACE2 Mice. Viruses. 2021 Jan 19;13(1):132. doi: 10.3390/v13010132. PMID: 33477869; PMCID: PMC7832889.

17. Liu G, Carter B, Bricken T, Jain S, Viard M, Carrington M, Gifford DK. Computationally Optimized SARS-CoV-2 MHC Class I and II Vaccine Formulations Predicted to Target Human Haplotype Distributions. Cell Syst. 2020 Aug 26;11(2):131-144.e6. doi: 10.1016/j.cels.2020.06.009. Epub 2020 Jul 27. PMID: 32721383; PMCID: PMC7384425.

18. Liu G, Carter B, Gifford DK. Predicted Cellular Immunity Population Coverage Gaps for SARS-CoV-2 Subunit Vaccines and Their Augmentation by Compact Peptide Sets. Cell Syst. 2021 Jan 20;12(1):102-107.e4. doi: 10.1016/j.cels.2020.11.010. Epub 2020 Nov 27. PMID: 33321075; PMCID: PMC7691134.

19. Liu, G., Carter B., Dimitrakakis A., Gifford D. K. Maximum n-times Coverage for Vaccine Design, International Conference on Learning Representations, ICLR (2022), https://arxiv.org/abs/2101.10902

20. Martinez DR, Schäfer A, Leist SR, De la Cruz G, West A, Atochina-Vasserman EN, Lindesmith LC, Pardi N, Parks R, Barr M, Li D, Yount B, Saunders KO, Weissman D, Haynes BF, Montgomery SA, Baric RS. Chimeric spike mRNA vaccines protect against Sarbecovirus challenge in mice. Science. 2021 Aug 27;373(6558):991–998. doi: 10.1126/science.abi4506. Epub 2021 Jun 22. PMID: 34214046; PMCID: PMC8899822.

21. Matchett WE, Joag V, Stolley JM, Shepherd FK, Quarnstrom CF, Mickelson CK, Wijeyesinghe S, Soerens AG, Becker S, Thiede JM, Weyu E, O’Flanagan S, Walter JA, Vu MN, Menachery VD, Bold TD, Vezys V, Jenkins MK, Langlois RA, Masopust D. Nucleocapsid vaccine elicits spike-independent SARS-CoV-2 protective immunity. J Immunol. 2021 Jun 30;: PMID: 33948591; PMCID: PMC8095198.

22. Moss P. The T cell immune response against SARS-CoV-2. Nat Immunol. 2022 Feb;23(2):186–193. doi: 10.1038/s41590-021-01122-w. Epub 2022 Feb 1. PMID: 35105982.

23. Nakiboneka R, Mugaba S, Auma BO, Kintu C, Lindan C, Nanteza MB, Kaleebu P, Serwanga J. Interferon gamma (IFN-γ) negative CD4+ and CD8+ T-cells can produce immune mediators in response to viral antigens. Vaccine. 2019 Jan 3;37(1):113–122. doi: 10.1016/j.vaccine.2018.11.024. Epub 2018 Nov 17. PMID: 30459072; PMCID: PMC6290111.

24. Nathan A, Rossin EJ, Kaseke C, Park RJ, Khatri A, Koundakjian D, Urbach JM, Singh NK, Bashirova A, Tano-Menka R, Senjobe F, Waring MT, Piechocka-Trocha A, Garcia-Beltran WF, Iafrate AJ, Naranbhai V, Carrington M, Walker BD, Gaiha GD. Structure-guided T cell vaccine design for SARS-CoV-2 variants and sarbecoviruses. Cell. 2021 Aug 19;184(17):4401-4413.e10. doi: 10.1016/j.cell.2021.06.029. Epub 2021 Jun 30. PMID: 34265281; PMCID: PMC8241654.

25. O’Donnell TJ, Rubinsteyn A, Laserson U. MHCflurry 2.0: Improved Pan-Allele Prediction of MHC Class I-Presented Peptides by Incorporating Antigen Processing. Cell Syst. 2020 Jul 22;11(1):42-48.e7. doi: 10.1016/j.cels.2020.06.010. Epub 2020 Jul 14. Erratum in: Cell Syst. 2020 Oct 21;11(4):418-419. PMID: 32711842.

26. Pardi N, Hogan MJ, Pelc RS, Muramatsu H, Andersen H, DeMaso CR, Dowd KA, Sutherland LL, Scearce RM, Parks R, Wagner W, Granados A, Greenhouse J, Walker M, Willis E, Yu JS, McGee CE, Sempowski GD, Mui BL, Tam YK, Huang YJ, Vanlandingham D, Holmes VM, Balachandran H, Sahu S, Lifton M, Higgs S, Hensley SE, Madden TD, Hope MJ, Karikó K, Santra S, Graham BS, Lewis MG, Pierson TC, Haynes BF, Weissman D. Zika virus protection by a single low-dose nucleoside-modified mRNA vaccination. Nature. 2017 Mar 9;543(7644):248–251. doi: 10.1038/nature21428. Epub 2017 Feb 2. PMID: 28151488; PMCID: PMC5344708.

27. Pardieck IN, van der Sluis TC, van der Gracht ETI, Veerkamp DMB, Behr FM, van Duikeren S, Beyrend G, Rip J, Nadafi R, Beyranvand Nejad E, Mülling N, Brasem DJ, Camps MGM, Myeni SK, Bredenbeek PJ, Kikkert M, Kim Y, Cicin-Sain L, Abdelaal T, van Gisbergen KPJM, Franken KLMC, Drijfhout JW, Melief CJM, Zondag GCM, Ossendorp F, Arens R. A third vaccination with a single T cell epitope confers protection in a murine model of SARS-CoV-2 infection. Nat Commun. 2022 Jul 8;13(1):3966. doi: 10.1038/s41467-022-31721-6. PMID: 35803932; PMCID: PMC9267705.

28. Redd AD, Nardin A, Kared H, Bloch EM, Pekosz A, Laeyendecker O, Abel B, Fehlings M, Quinn TC, Tobian AA. CD8+ T cell responses in COVID-19 convalescent individuals target conserved epitopes from multiple prominent SARS-CoV-2 circulating variants. medRxiv [Preprint]. 2021 Feb 12:2021.02.11.21251585. doi: 10.1101/2021.02.11.21251585. Update in: Open Forum Infect Dis. 2021 Mar 30;8(7):ofab143. PMID: 33594378; PMCID: PMC7885937.

29. Reynisson B, Alvarez B, Paul S, Peters B, Nielsen M. NetMHCpan-4.1 and NetMHCIIpan-4.0: improved predictions of MHC antigen presentation by concurrent motif deconvolution and integration of MS MHC eluted ligand data. Nucleic Acids Res. 2020 Jul 2;48(W1):W449–W454. doi: 10.1093/nar/gkaa379. PMID: 32406916; PMCID: PMC7319546.

30. Sahin U, Derhovanessian E, Miller M, Kloke BP, Simon P, Löwer M, Bukur V, Tadmor AD, Luxemburger U, Schrörs B, Omokoko T, Vormehr M, Albrecht C, Paruzynski A, Kuhn AN, Buck J, Heesch S, Schreeb KH, Müller F, Ortseifer I, Vogler I, Godehardt E, Attig S, Rae R, Breitkreuz A, Tolliver C, Suchan M, Martic G, Hohberger A, Sorn P, Diekmann J, Ciesla J, Waksmann O, Brück AK, Witt M, Zillgen M, Rothermel A, Kasemann B, Langer D, Bolte S, Diken M, Kreiter S, Nemecek R, Gebhardt C, Grabbe S, Höller C, Utikal J, Huber C, Loquai C, Türeci Ö. Personalized RNA mutanome vaccines mobilize poly-specific therapeutic immunity against cancer. Nature. 2017 Jul 13;547(7662):222–226. doi: 10.1038/nature23003. Epub 2017 Jul 5. PMID: 28678784.

31. Seabold, S., Perktold, J. Statsmodels: Econometric and statistical modeling with python. Proceedings of the 9th Python in Science Conference. 2010; 57(61).

32. Sekine T, Perez-Potti A, Rivera-Ballesteros O, Strålin K, Gorin JB, Olsson A, Llewellyn-Lacey S, Kamal H, Bogdanovic G, Muschiol S, Wullimann DJ, Kammann T, Emgård J, Parrot T, Folkesson E; Karolinska COVID-19 Study Group, Rooyackers O, Eriksson LI, Henter JI, Sönnerborg A, Allander T, Albert J, Nielsen M, Klingström J, Gredmark-Russ S, Björkström NK, Sandberg JK, Price DA, Ljunggren HG, Aleman S, Buggert M. Robust T Cell Immunity in Convalescent Individuals with Asymptomatic or Mild COVID-19. Cell. 2020 Oct 1;183(1):158-168.e14. doi: 10.1016/j.cell.2020.08.017. Epub 2020 Aug 14. PMID: 32979941; PMCID: PMC7427556.

33. Shuai H, Chan JF, Yuen TT, Yoon C, Hu JC, Wen L, Hu B, Yang D, Wang Y, Hou Y, Huang X, Chai Y, Chan CC, Poon VK, Lu L, Zhang RQ, Chan WM, Ip JD, Chu AW, Hu YF, Cai JP, Chan KH, Zhou J, Sridhar S, Zhang BZ, Yuan S, Zhang AJ, Huang JD, To KK, Yuen KY, Chu H. Emerging SARS-CoV-2 variants expand species tropism to murines. EBioMedicine. 2021 Nov;73:103643. doi: 10.1016/j.ebiom.2021.103643. Epub 2021 Oct 21. PMID: 34689086; PMCID: PMC8530107.

34. Snyder TM, Gittelman RM, Klinger M, May DH, Osborne EJ, Taniguchi R, Zahid HJ, Kaplan IM, Dines JN, Noakes MT, Pandya R, Chen X, Elasady S, Svejnoha E, Ebert P, Pesesky MW, De Almeida P, O’Donnell H, DeGottardi Q, Keitany G, Lu J, Vong A, Elyanow R, Fields P, Greissl J, Baldo L, Semprini S, Cerchione C, Nicolini F, Mazza M, Delmonte OM, Dobbs K, Laguna-Goya R, Carreño-Tarragona G, Barrio S, Imberti L, Sottini A, Quiros-Roldan E, Rossi C, Biondi A, Bettini LR, D’Angio M, Bonfanti P, Tompkins MF, Alba C, Dalgard C, Sambri V, Martinelli G, Goldman JD, Heath JR, Su HC, Notarangelo LD, Paz-Artal E, Martinez-Lopez J, Carlson JM, Robins HS. Magnitude and Dynamics of the T-Cell Response to SARS-CoV-2 Infection at Both Individual and Population Levels. medRxiv [Preprint]. 2020 Sep 17:2020.07.31.20165647. doi: 10.1101/2020.07.31.20165647. PMID: 32793919; PMCID: PMC7418734.

35. Stirling DR, Swain-Bowden MJ, Lucas AM, Carpenter AE, Cimini BA, Goodman A. CellProfiler 4: improvements in speed, utility and usability. BMC Bioinformatics. 2021 Sep 10;22(1):433. doi: 10.1186/s12859-021-04344-9. PMID: 34507520; PMCID: PMC8431850.

36. Swank Z, Senussi Y, Manickas-Hill Z, Yu XG, Li JZ, Alter G, Walt DR. Persistent circulating SARS-CoV-2 spike is associated with post-acute COVID-19 sequelae. Clin Infect Dis. 2022 Sep 2:ciac722. doi: 10.1093/cid/ciac722. Epub ahead of print. PMID: 36052466.

37. Tregoning JS, Flight KE, Higham SL, Wang Z, Pierce BF. Progress of the COVID-19 vaccine effort: viruses, vaccines and variants versus efficacy, effectiveness and escape. Nat Rev Immunol. 2021 Oct;21(10):626–636. doi: 10.1038/s41577-021-00592-1. Epub 2021 Aug 9. PMID: 34373623; PMCID: PMC8351583.

38. Tseng CTK, Huang C, Newman P, Wang N, Narayanan K, Watts DM, Makino S, Packard M, Zaki SR, Chan TS, and Peters CJ. 2007. SARS Coronavirus Infection of Transgenic Mice bearing the Human Angiotensin Converting Enzyme 2 (hACE2) Virus Receptor. JVI 81: 1162–1173.

39. Tseng CT, Sbrana E, Iwata-Yoshikawa N, Newman PC, Garron T, Atmar RL, Peters CJ, Couch RB. Immunization with SARS coronavirus vaccines leads to pulmonary immunopathology on challenge with the SARS virus. PLoS One. 2012;7(4):e35421. doi: 10.1371/journal.pone.0035421. Epub 2012 Apr 20. Erratum in: PLoS One. 2012;7(8). doi:10.1371/annotation/2965cfae-b77d-4014-8b7b-236e01a35492. PMID: 22536382; PMCID: PMC3335060.

40. UniProt Consortium. UniProt: a worldwide hub of protein knowledge. Nucleic Acids Res. 2019 Jan 8;47(D1):D506–D515. doi: 10.1093/nar/gky1049. PMID: 30395287; PMCID: PMC6323992.

41. Virtanen P, Gommers R, Oliphant TE, Haberland M, Reddy T, Cournapeau D, Burovski E, Peterson P, Weckesser W, Bright J, van der Walt SJ, Brett M, Wilson J, Millman KJ, Mayorov N, Nelson ARJ, Jones E, Kern R, Larson E, Carey CJ, Polat İ, Feng Y, Moore EW, VanderPlas J, Laxalde D, Perktold J, Cimrman R, Henriksen I, Quintero EA, Harris CR, Archibald AM, Ribeiro AH, Pedregosa F, van Mulbregt P; SciPy 1.0 Contributors. SciPy 1.0: fundamental algorithms for scientific computing in Python. Nat Methods. 2020 Mar;17(3):261–272. doi: 10.1038/s41592-019-0686-2. Epub 2020 Feb 3. Erratum in: Nat Methods. 2020 Feb 24;: PMID: 32015543; PMCID: PMC7056644.

42. Vita R, Mahajan S, Overton JA, Dhanda SK, Martini S, Cantrell JR, Wheeler DK, Sette A, Peters B. The Immune Epitope Database (IEDB): 2018 update. Nucleic Acids Res. 2018 Oct 24. doi: 10.1093/nar/gky1006. [Epub ahead of print] PubMed PMID: 30357391.

43. Walsh EE, Frenck RW Jr, Falsey AR, Kitchin N, Absalon J, Gurtman A, Lockhart S, Neuzil K, Mulligan MJ, Bailey R, Swanson KA, Li P, Koury K, Kalina W, Cooper D, Fontes-Garfias C, Shi PY, Türeci Ö, Tompkins KR, Lyke KE, Raabe V, Dormitzer PR, Jansen KU, Şahin U, Gruber WC. Safety and Immunogenicity of Two RNA-Based Covid-19 Vaccine Candidates. N Engl J Med. 2020 Dec 17;383(25):2439–2450. doi: 10.1056/NEJMoa2027906. Epub 2020 Oct 14. PMID: 33053279; PMCID: PMC7583697.

44. Willett BJ, Grove J, MacLean OA, Wilkie C, De Lorenzo G, Furnon W, Cantoni D, Scott S, Logan N, Ashraf S, Manali M, Szemiel A, Cowton V, Vink E, Harvey WT, Davis C, Asamaphan P, Smollett K, Tong L, Orton R, Hughes J, Holland P, Silva V, Pascall DJ, Puxty K, da Silva Filipe A, Yebra G, Shaaban S, Holden MTG, Pinto RM, Gunson R, Templeton K, Murcia PR, Patel AH, Klenerman P, Dunachie S; PITCH Consortium; COVID-19 Genomics UK (COG-UK) Consortium, Haughney J, Robertson DL, Palmarini M, Ray S, Thomson EC. SARS-CoV-2 Omicron is an immune escape variant with an altered cell entry pathway. Nat Microbiol. 2022 Aug;7(8):1161–1179. doi: 10.1038/s41564-022-01143-7. Epub 2022 Jul 7. PMID: 35798890; PMCID: PMC9352574.

45. World Health Organization (WHO). In the wake of the pandemic: preparing for Long COVID (2021). Policy Brief 39. Accessed August 14, 2022. https://apps.who.int/iris/bitstream/handle/10665/339629/Policy-brief-39-1997-8073-eng.pdf

46. Zeng H, Gifford DK. Quantification of Uncertainty in Peptide-MHC Binding Prediction Improves High-Affinity Peptide Selection for Therapeutic Design. Cell Syst. 2019 Aug 28;9(2):159-166.e3. doi: 10.1016/j.cels.2019.05.004. Epub 2019 Jun 5. PMID: 31176619; PMCID: PMC6715517.

47. Zhang X, Wu S, Wu B, Yang Q, Chen A, Li Y, Zhang Y, Pan T, Zhang H, He X. SARS-CoV-2 Omicron strain exhibits potent capabilities for immune evasion and viral entrance. Signal Transduct Target Ther. 2021 Dec 17;6(1):430. doi: 10.1038/s41392-021-00852-5. PMID: 34921135; PMCID: PMC8678971.

48. Zhao J, Zhao J, Mangalam AK, Channappanavar R, Fett C, Meyerholz DK, Agnihothram S, Baric RS, David CS, Perlman S. Airway Memory CD4(+) T Cells Mediate Protective Immunity against Emerging Respiratory Coronaviruses. Immunity. 2016 Jun 21;44(6):1379–91. doi: 10.1016/j.immuni.2016.05.006. Epub 2016 Jun 7. PMID: 27287409; PMCID: PMC4917442.

