## Supplemental Figure 7 for "A pan-variant mRNA-LNP T cell vaccine protects HLA transgenic mice from mortality after infection with SARS-CoV-2 Beta"

Supplemental Figure 7
Page 1/29

MIT-T-COVID Lung 1-1
2 dpi

CD8$^+$/CD4$^+$ Cell Annotations

Nucleated Cell Annotations

CD8$^+$

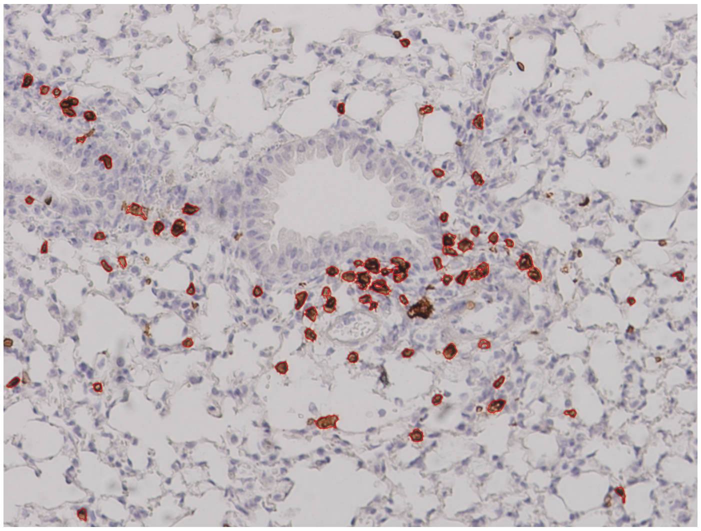

75 CD8$^+$ cells

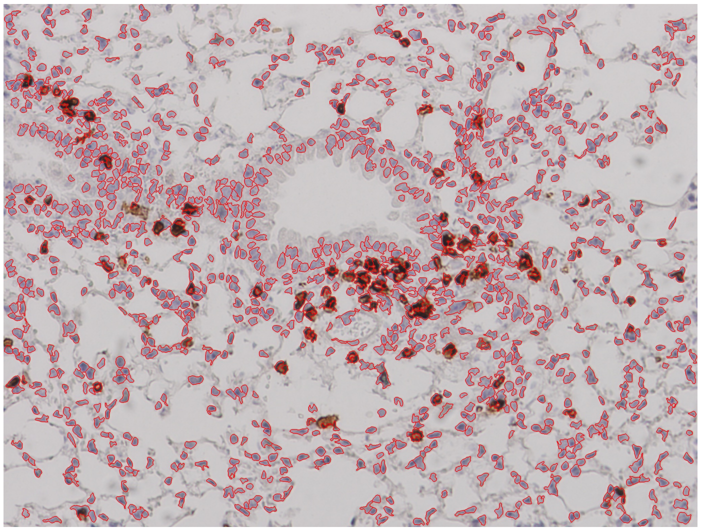

1170 nucleated cells

CD4$^+$

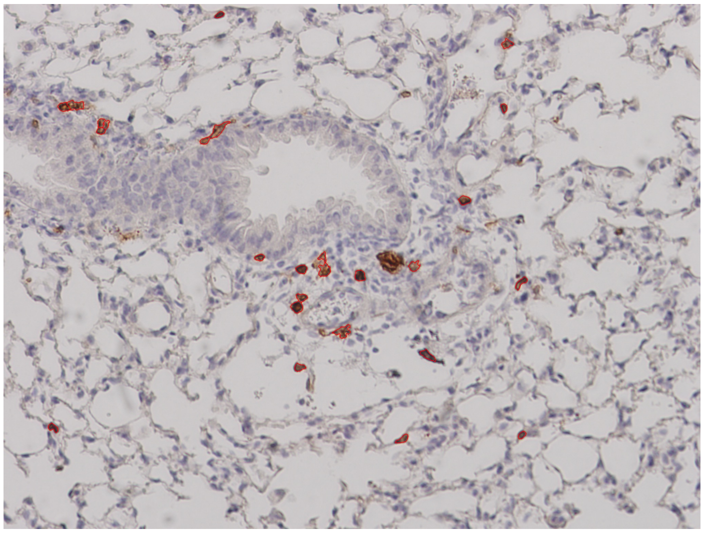

26 CD4$^+$ cells

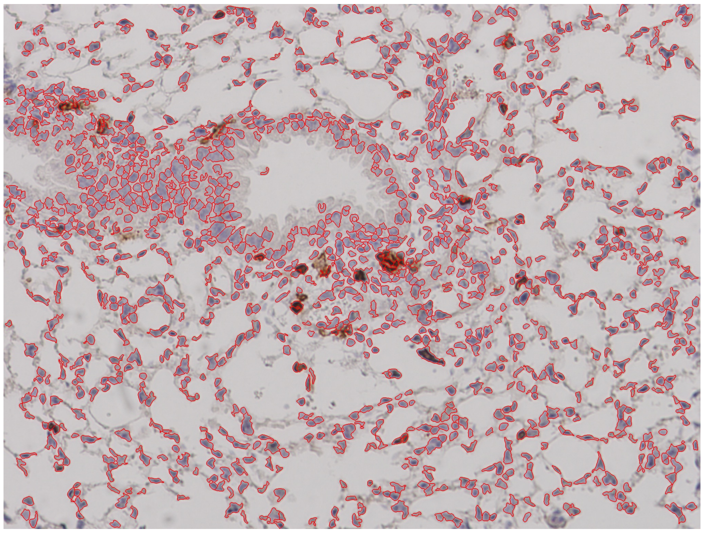

1194 nucleated cells

Supplemental Figure 7
Page 2/29

MIT-T-COVID Lung 1-2
2 dpi

CD8$^+$/CD4$^+$ Cell Annotations | Nucleated Cell Annotations

CD8$^+$

CD4$^+$

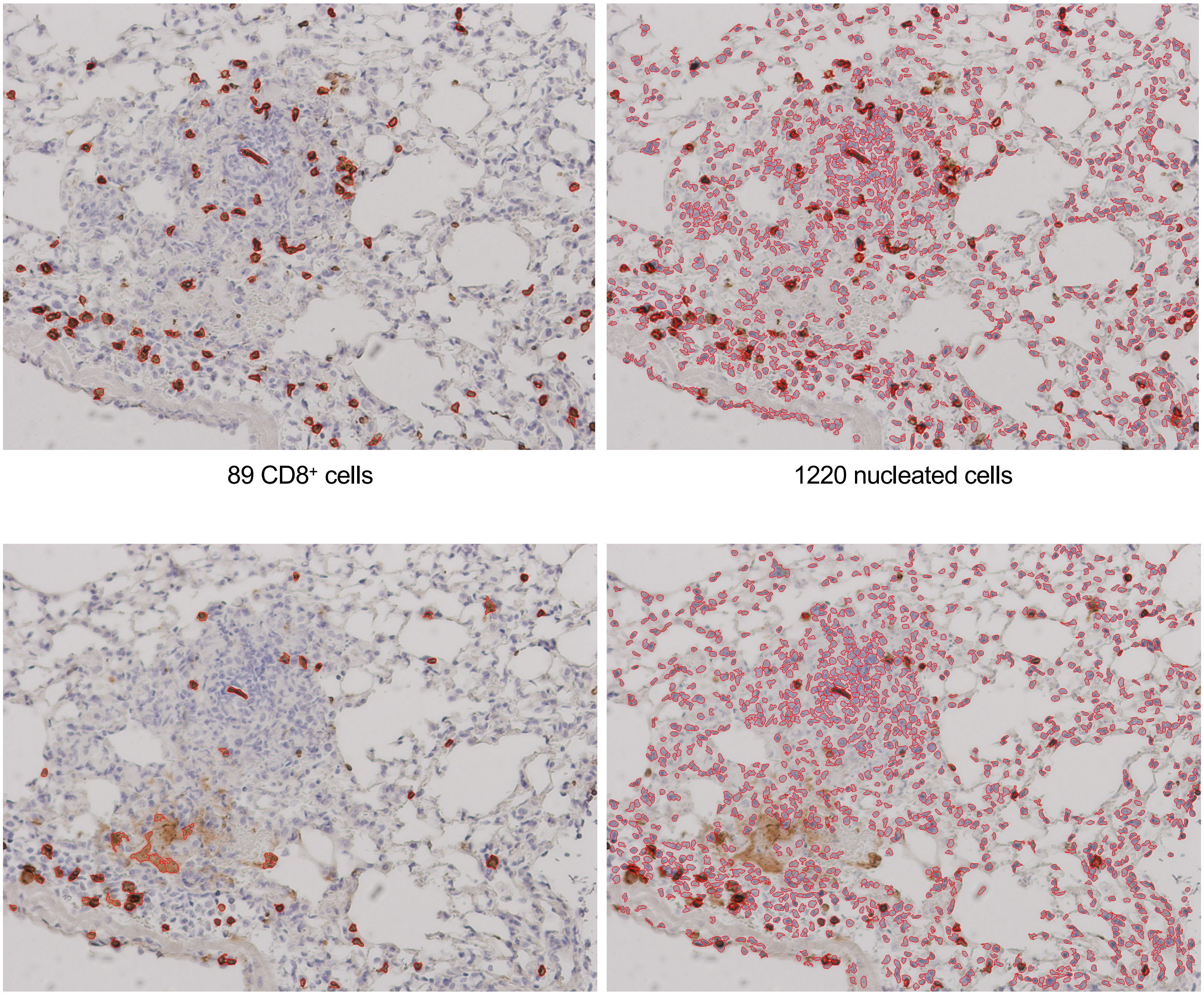

89 CD8$^+$ cells

1220 nucleated cells

56 CD4$^+$ cells

1248 nucleated cells

### MIT-T-COVID Lung 1-3
#### 2 dpi

CD8$^+$/CD4$^+$ Cell Annotations | Nucleated Cell Annotations

CD8$^+$

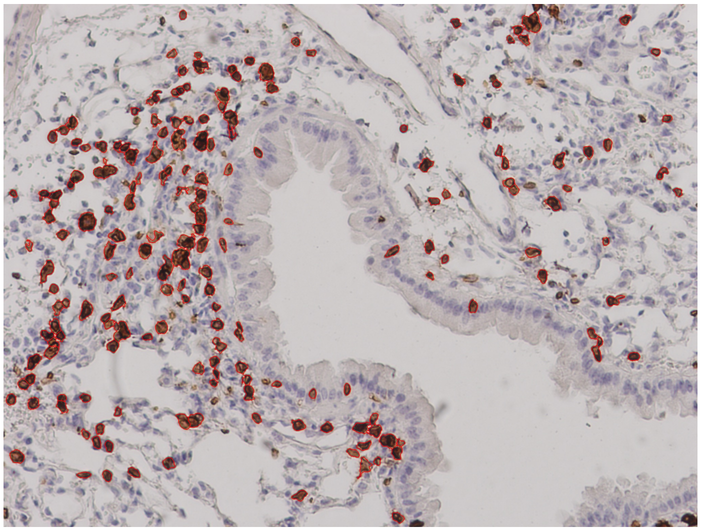

188 CD8$^+$ cells

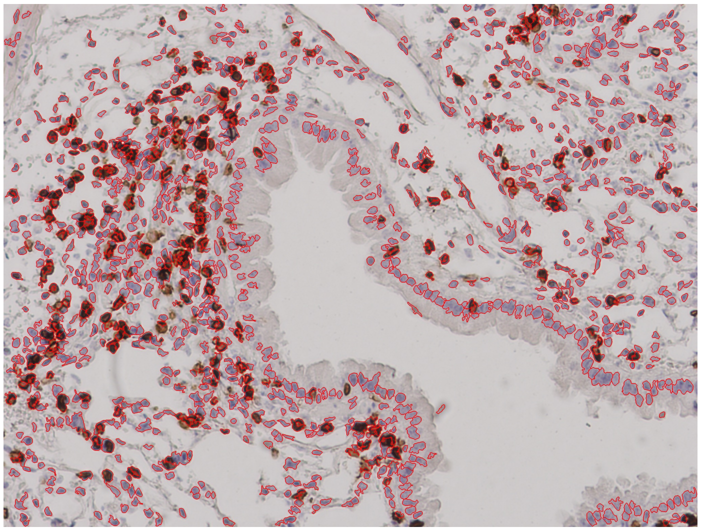

1156 nucleated cells

CD4$^+$

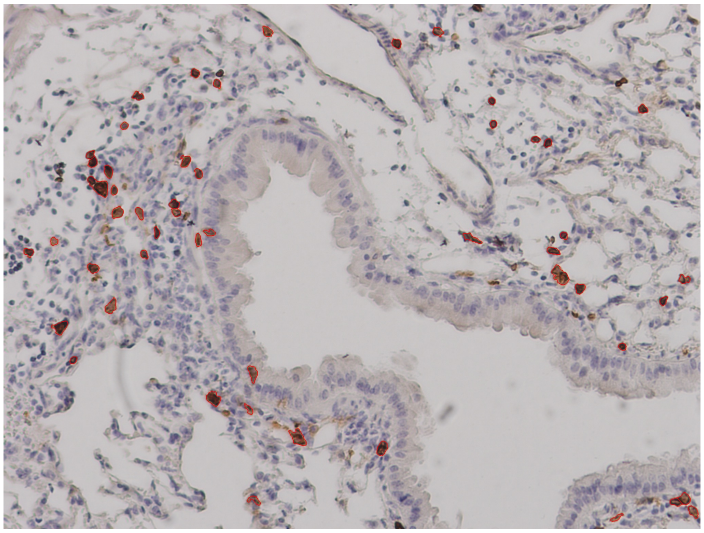

51 CD4$^+$ cells

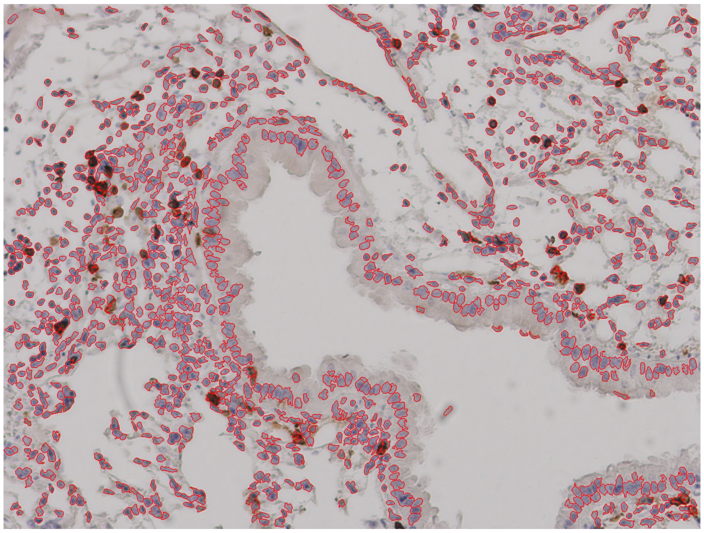

1121 nucleated cells

Supplemental Figure 7
Page 4/29

MIT-T-COVID Lung 2-1
2 dpi

CD8$^+$/CD4$^+$ Cell Annotations

Nucleated Cell Annotations

CD8$^+$

CD4$^+$

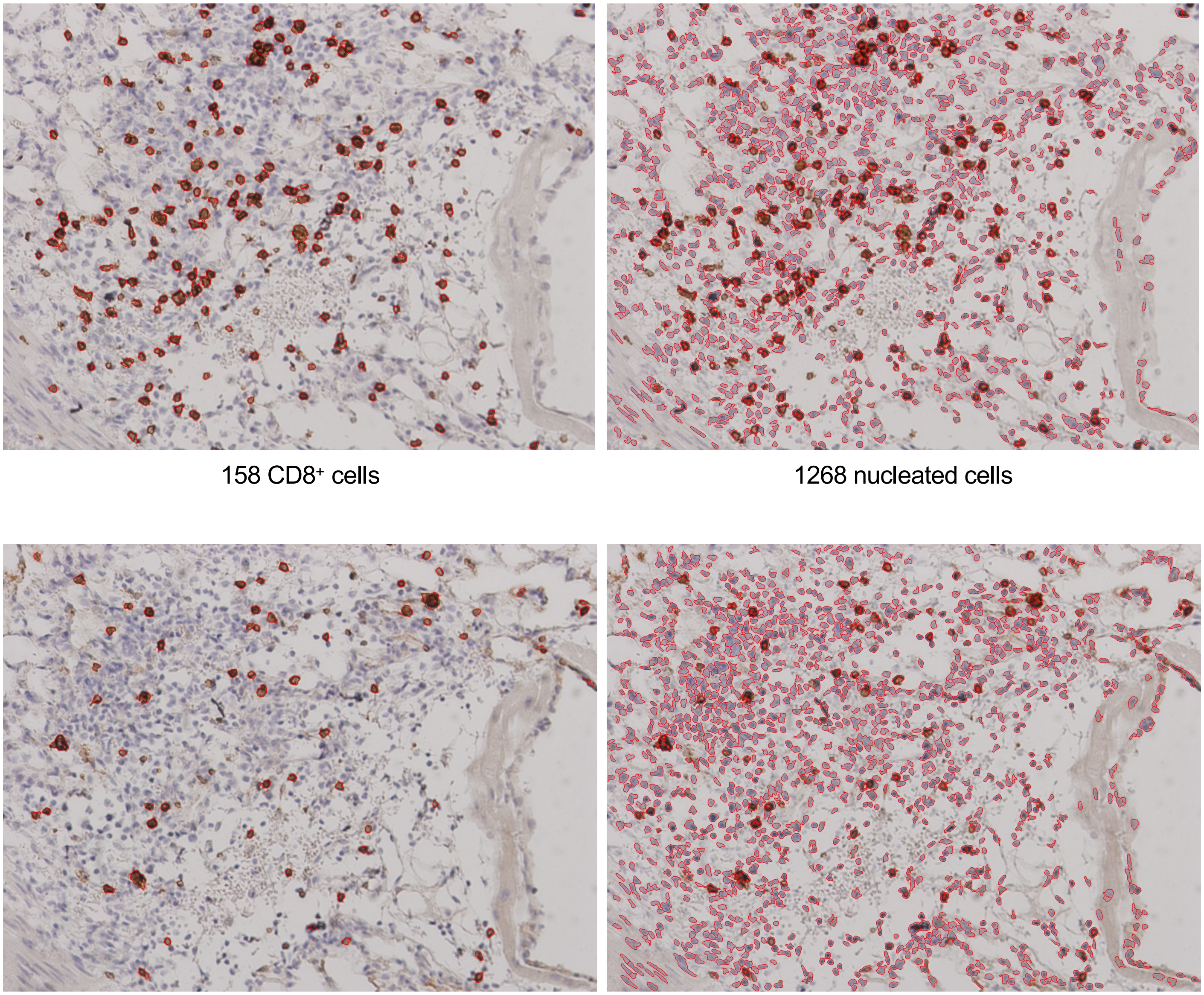

158 CD8$^+$ cells

1268 nucleated cells

61 CD4$^+$ cells

1309 nucleated cells

Supplemental Figure 7


MIT-T-COVID Lung 3-1
2 dpi

$CD8^+$/$CD4^+$ Cell Annotations | Nucleated Cell Annotations

$CD8^+$

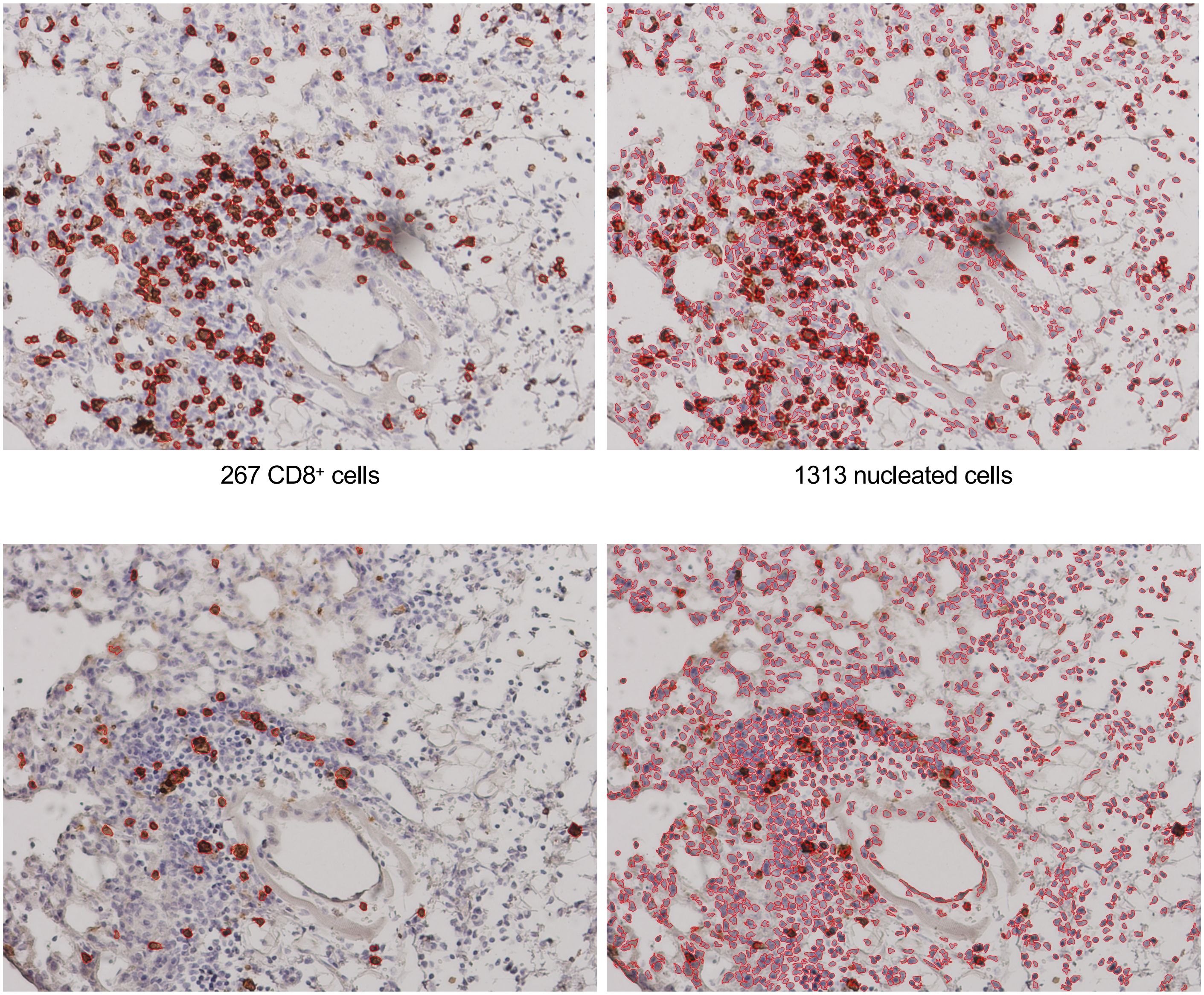

267 $CD8^+$ cells | 1313 nucleated cells

$CD4^+$

56 $CD4^+$ cells | 1455 nucleated cells

### Supplemental Figure 7


#### Pfizer/BNT Lung 1-1
2 dpi

CD8$^+$/CD4$^+$ Cell Annotations

Nucleated Cell Annotations

CD8$^+$

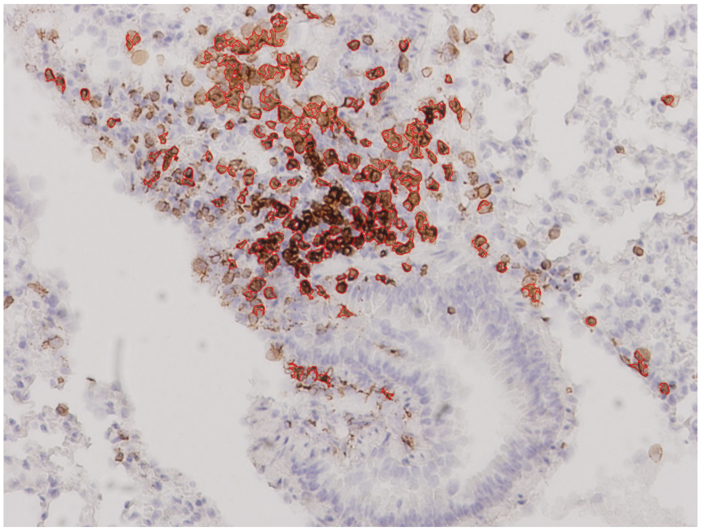

152 CD8$^+$ cells

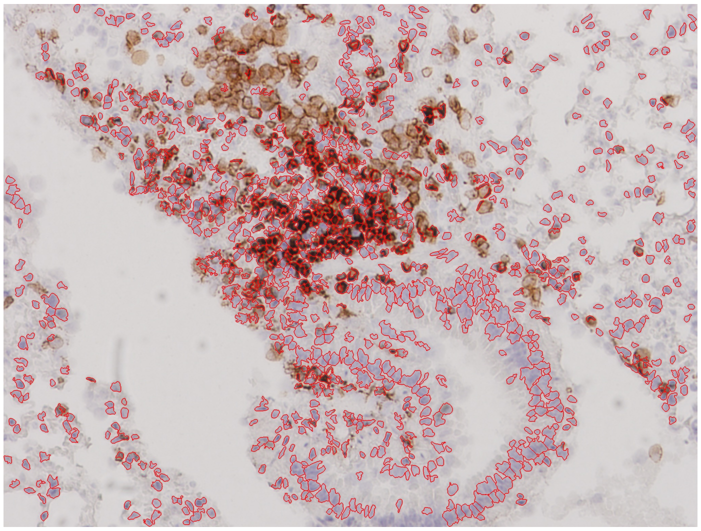

1032 nucleated cells

CD4$^+$

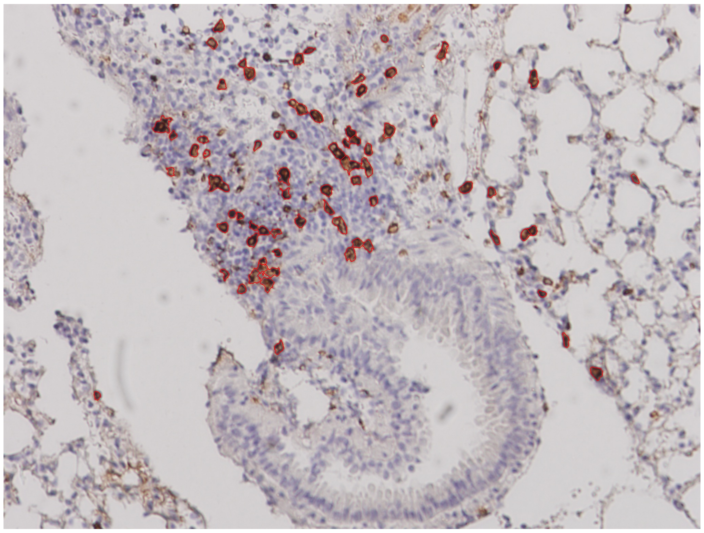

72 CD4$^+$ cells

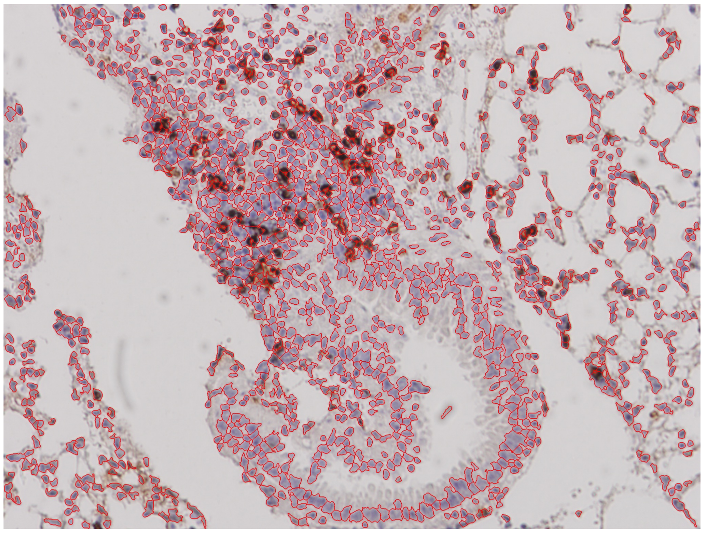

1386 nucleated cells

### Supplemental Figure 7



#### Pfizer/BNT Lung 2-1
#### 2 dpi

| | CD8$^+$/CD4$^+$ Cell Annotations | Nucleated Cell Annotations |
|---|---|---|
| CD8$^+$ | 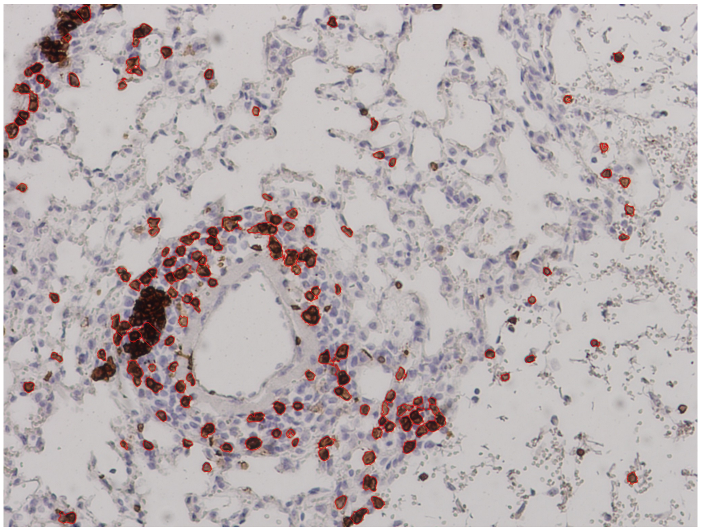 | 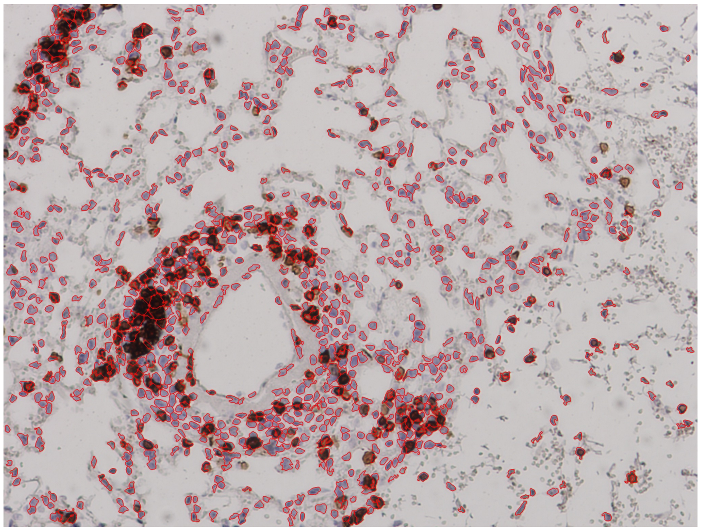 |
| | 138 CD8$^+$ cells | 989 nucleated cells |
| CD4$^+$ | 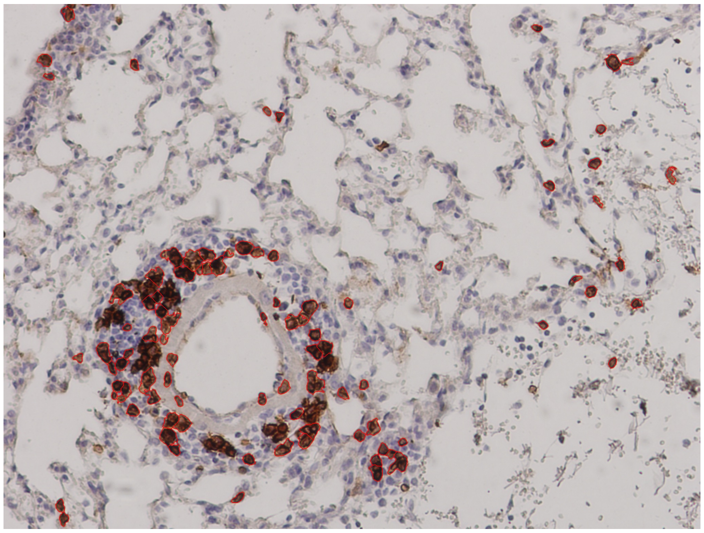 | 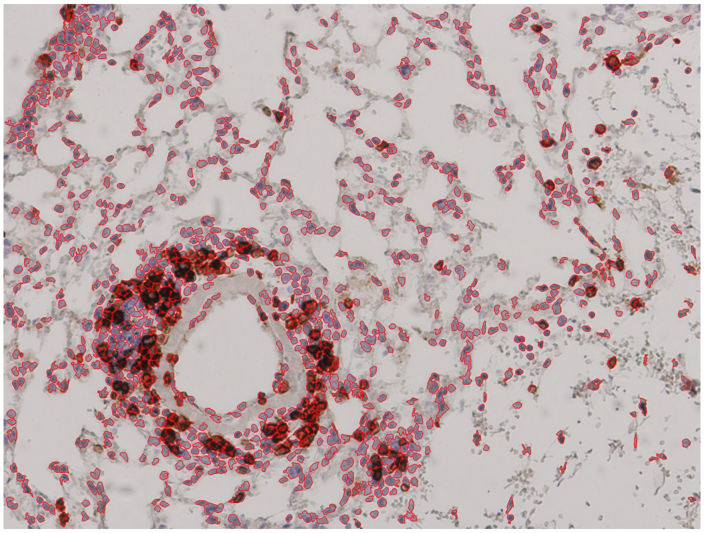 |
| | 90 CD4$^+$ cells | 1009 nucleated cells |

Supplemental Figure 7


Pfizer/BNT Lung 2-2
2 dpi

CD8$^+$/CD4$^+$ Cell Annotations | Nucleated Cell Annotations

CD8$^+$

CD4$^+$

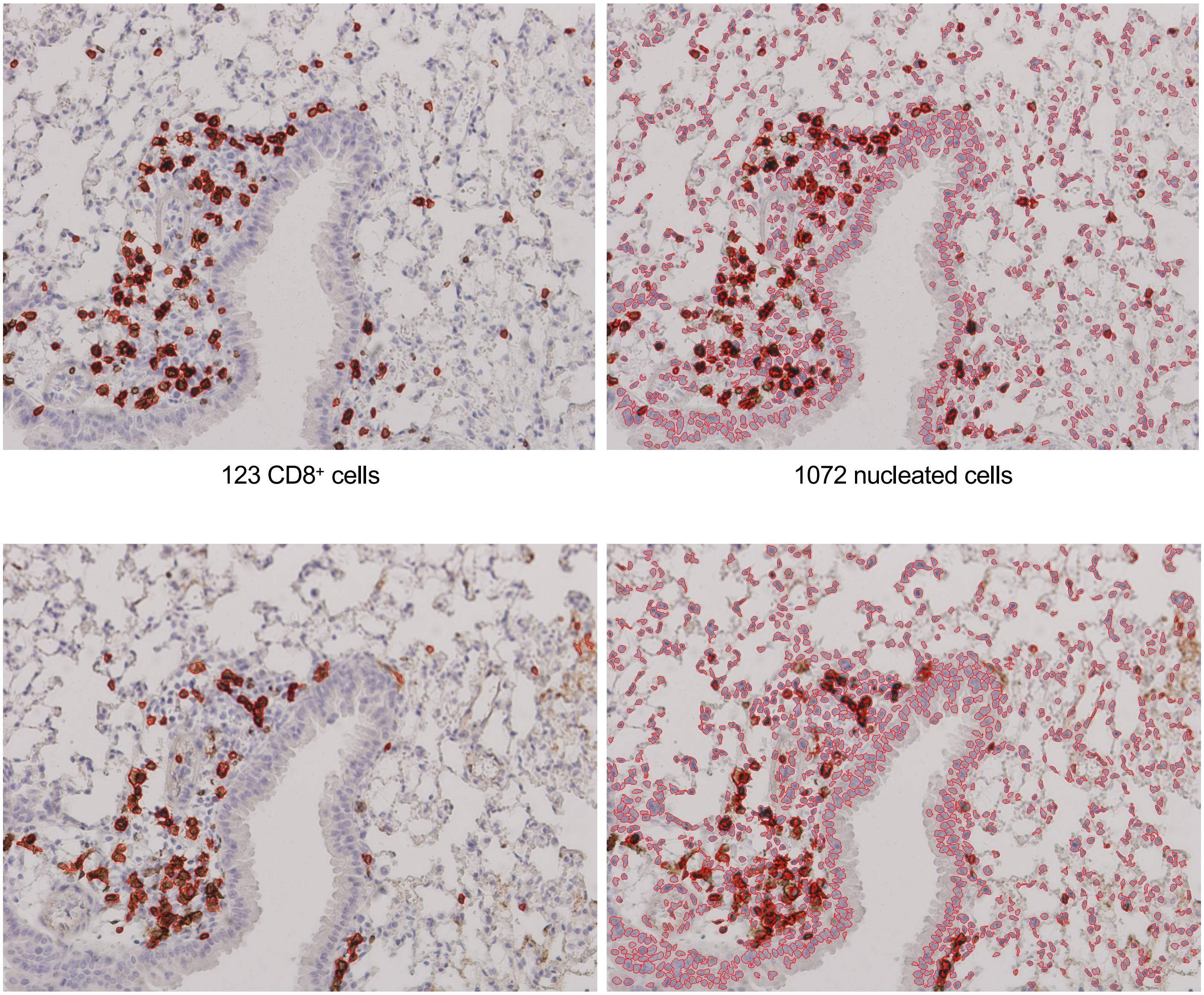

123 CD8$^+$ cells | 1072 nucleated cells

82 CD4$^+$ cells | 1178 nucleated cells

Supplemental Figure 7


Pfizer/BNT Lung 3-1
2 dpi

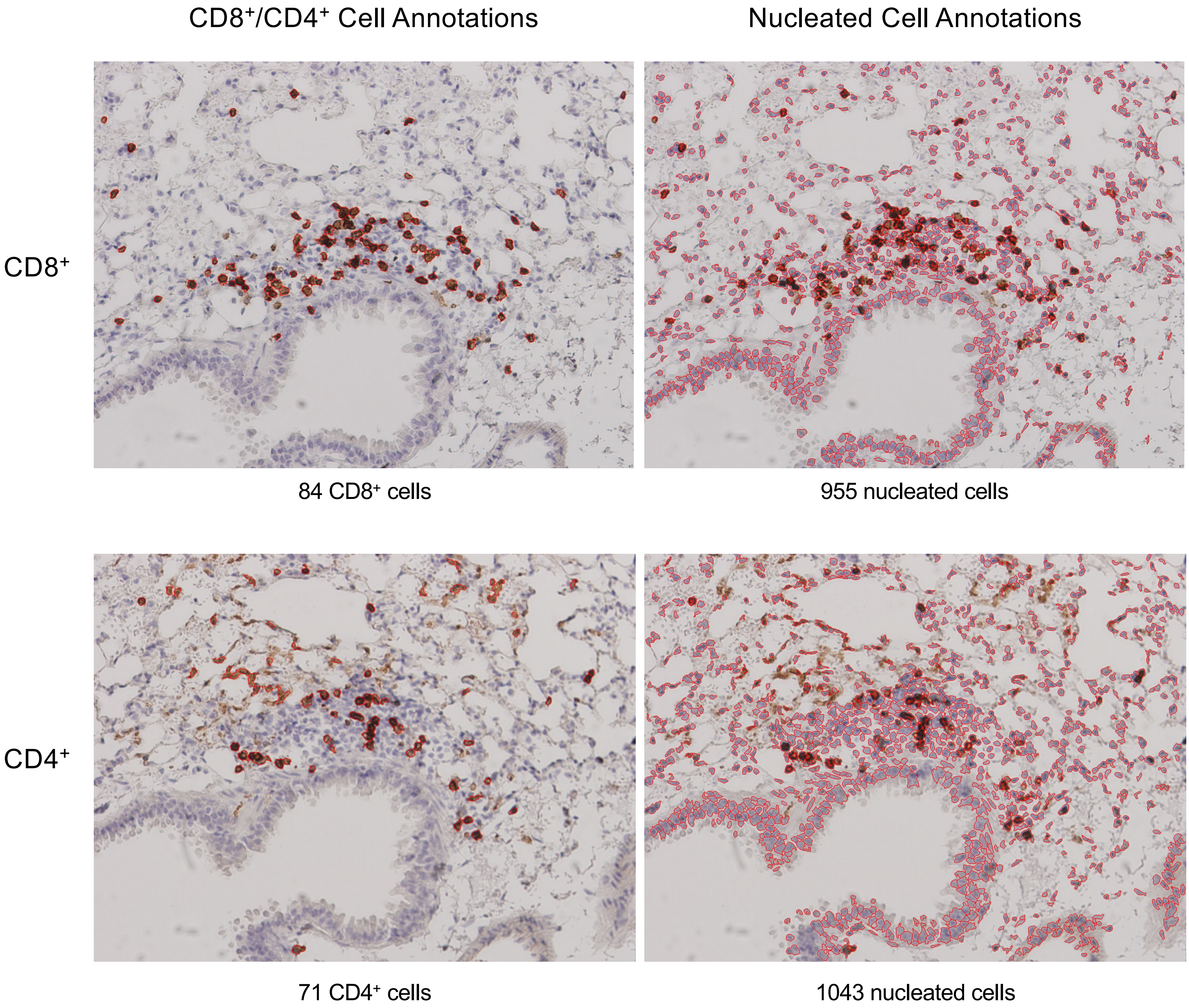

Supplemental Figure 7


Pfizer/BNT Lung 3-2
2 dpi

CD8$^{+}$/CD4$^{+}$ Cell Annotations | Nucleated Cell Annotations

CD8$^{+}$

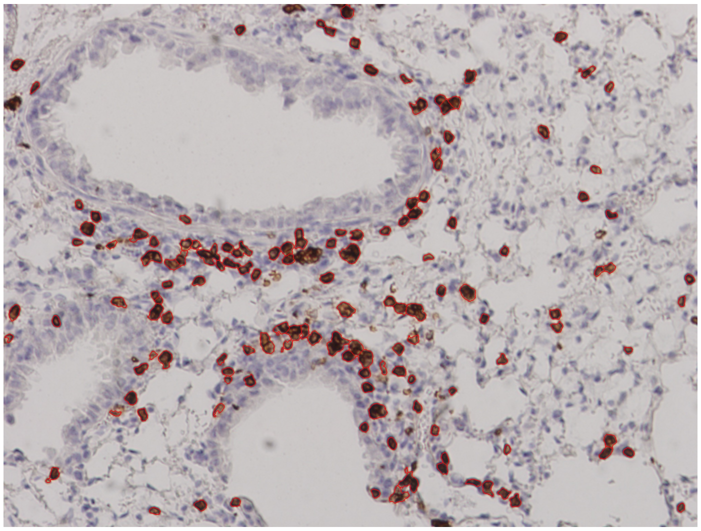

139 CD8$^{+}$ cells

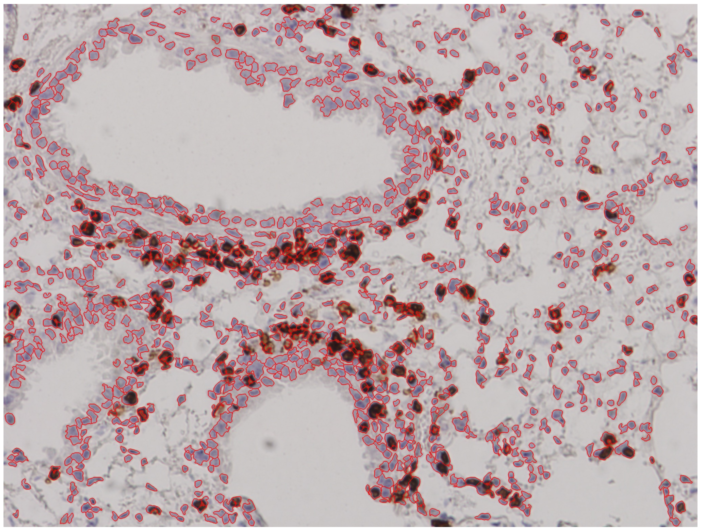

1064 nucleated cells

CD4$^{+}$

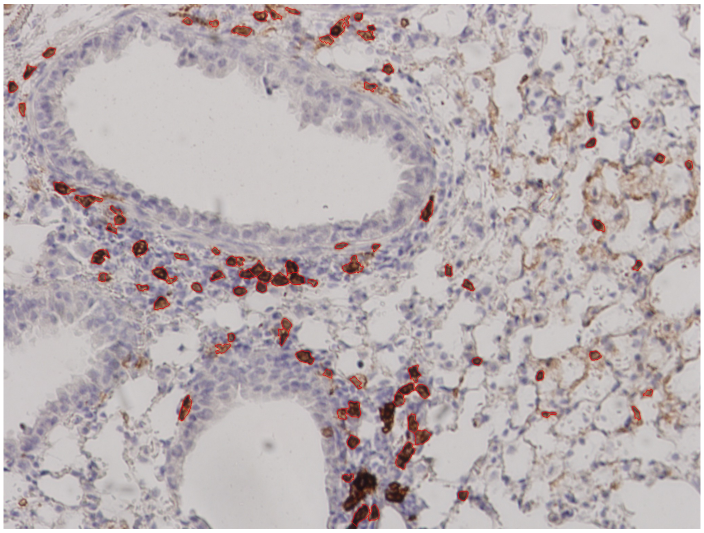

85 CD4$^{+}$ cells

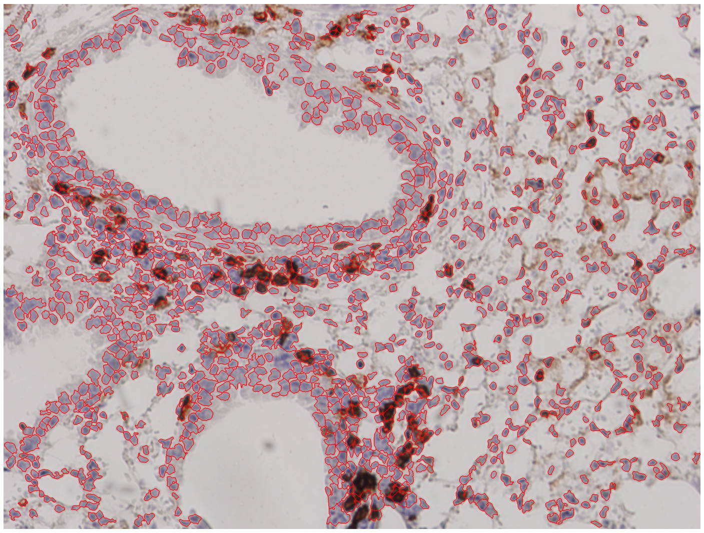

1165 nucleated cells

Supplemental Figure 7


PBS Lung 1-1
2 dpi

CD8$^+$/CD4$^+$ Cell Annotations

Nucleated Cell Annotations

CD8$^+$

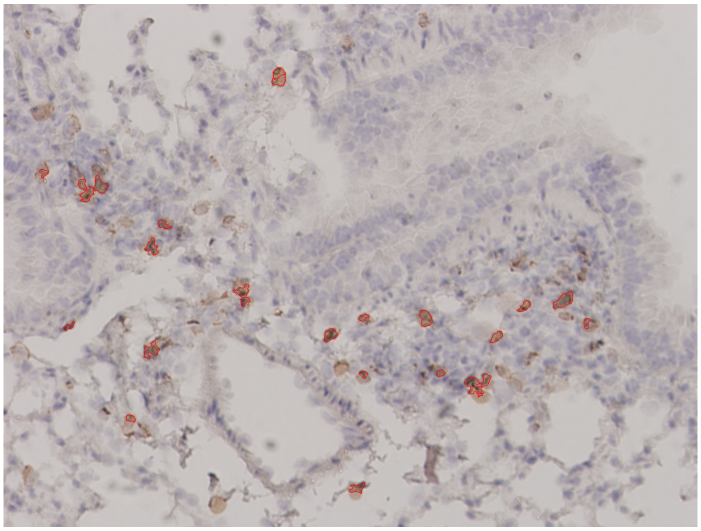

25 CD8$^+$ cells

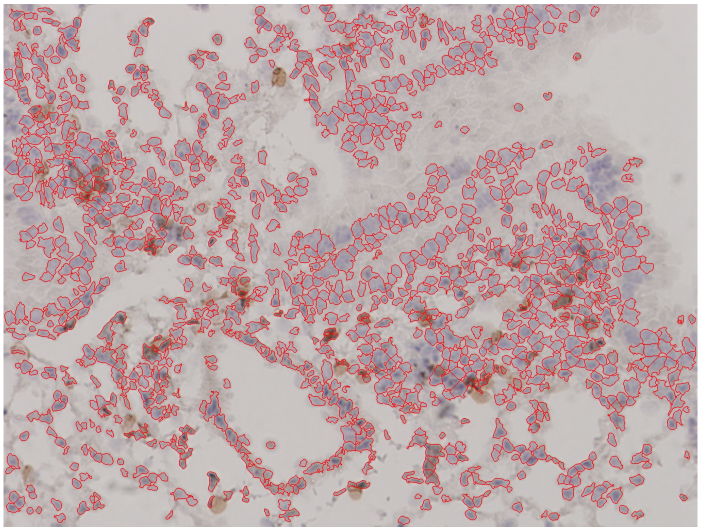

1165 nucleated cells

CD4$^+$

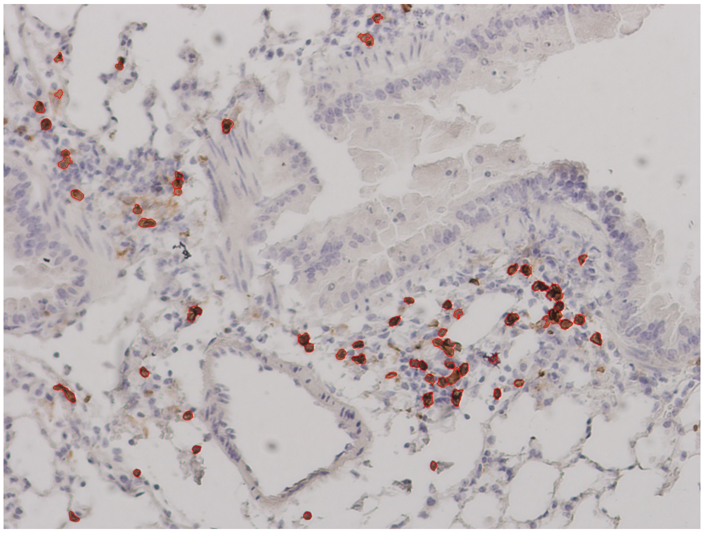

63 CD4$^+$ cells

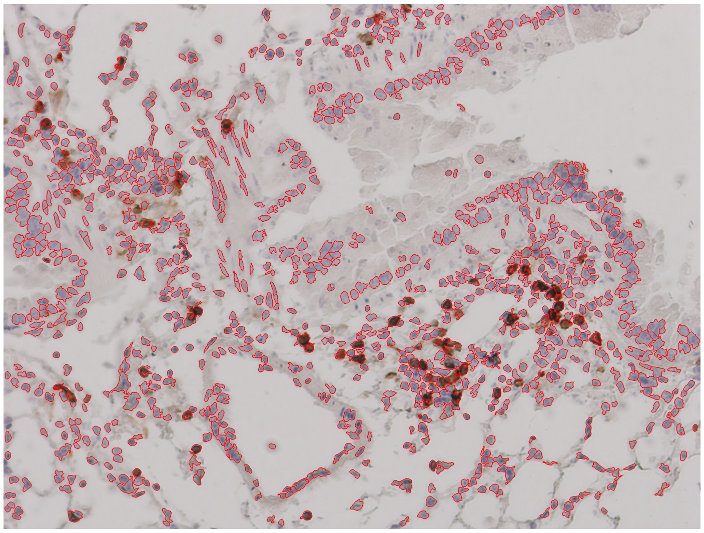

952 nucleated cells

PBS Lung 2-1
2 dpi

CD8$^+$/CD4$^+$ Cell Annotations | Nucleated Cell Annotations

CD8$^+$

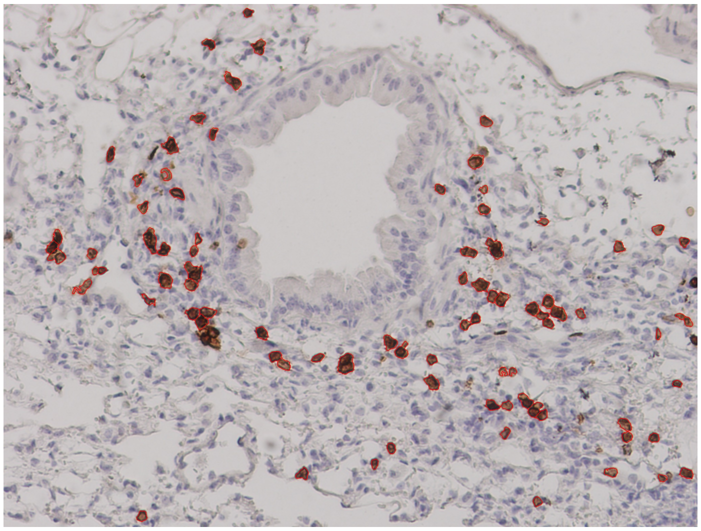

85 CD8$^+$ cells

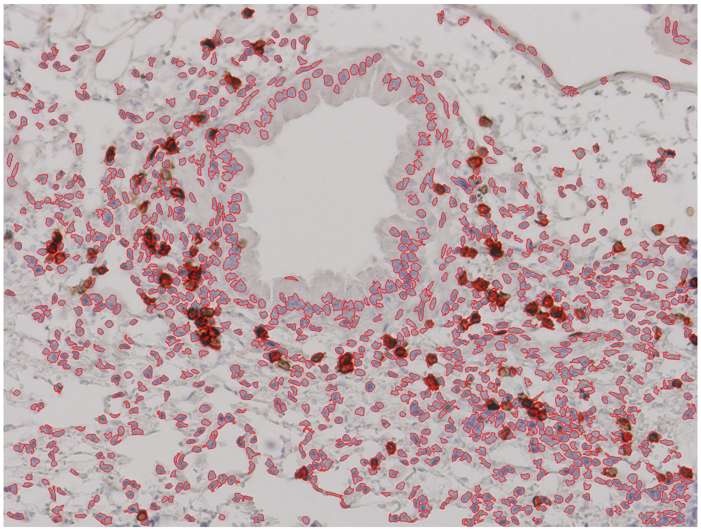

1166 nucleated cells

CD4$^+$

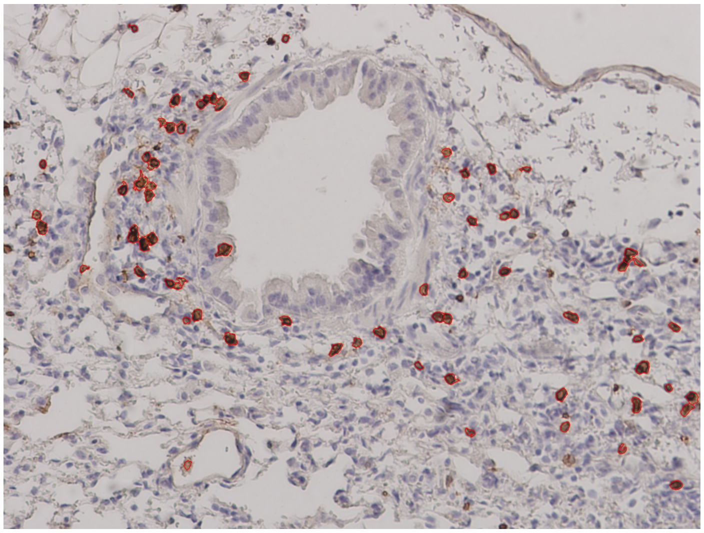

69 CD4$^+$ cells

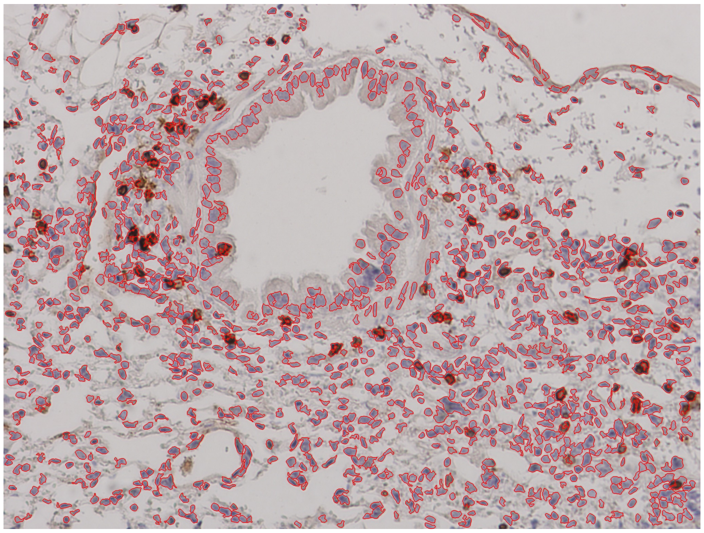

1229 nucleated cells

PBS Lung 2-2
2 dpi

CD8$^+$/CD4$^+$ Cell Annotations | Nucleated Cell Annotations

CD8$^+$

CD4$^+$

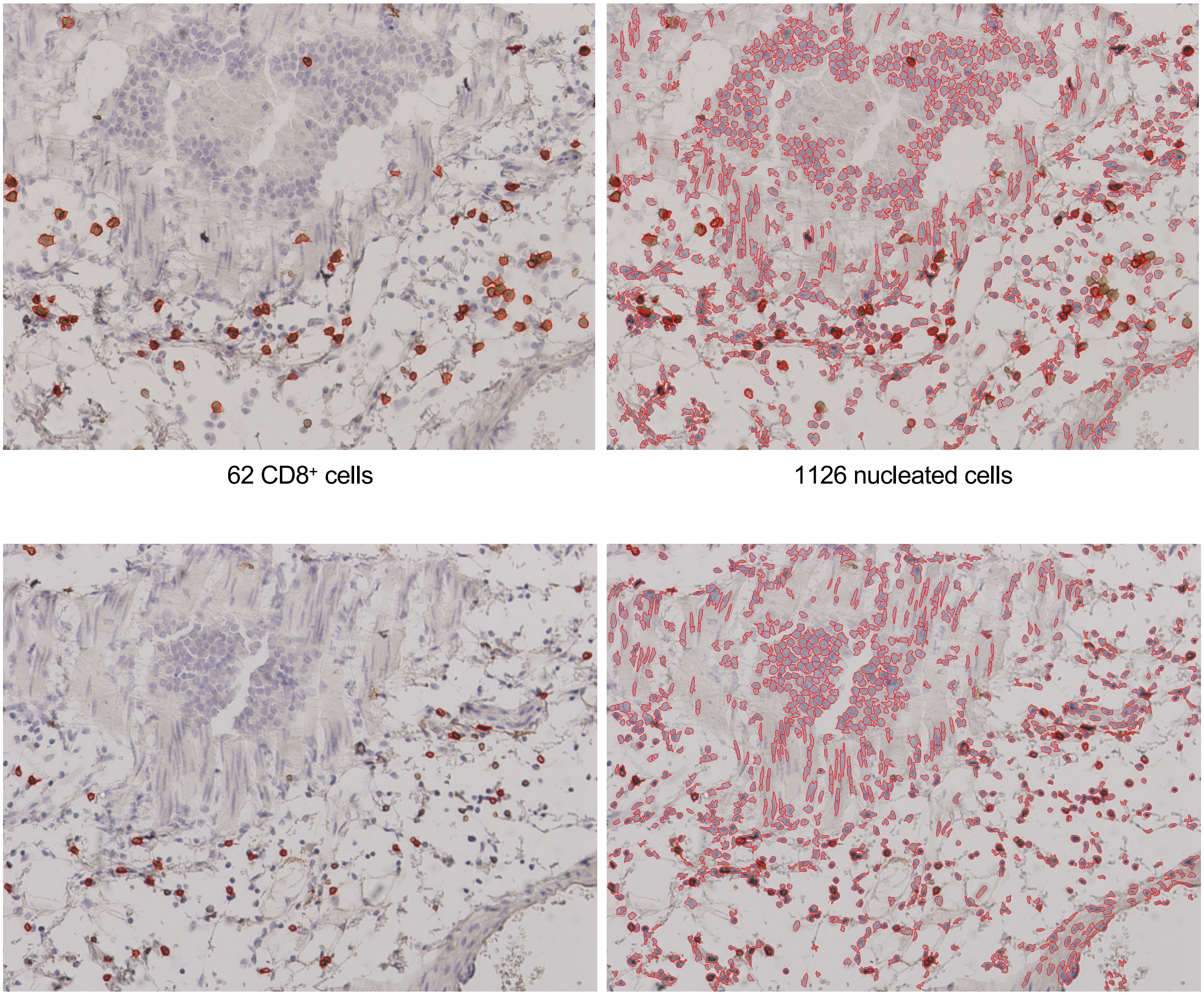

62 CD8$^+$ cells | 1126 nucleated cells

37 CD4$^+$ cells | 1034 nucleated cells

PBS Lung 3-1
2 dpi

CD8$^+$/CD4$^+$ Cell Annotations | Nucleated Cell Annotations

CD8$^+$

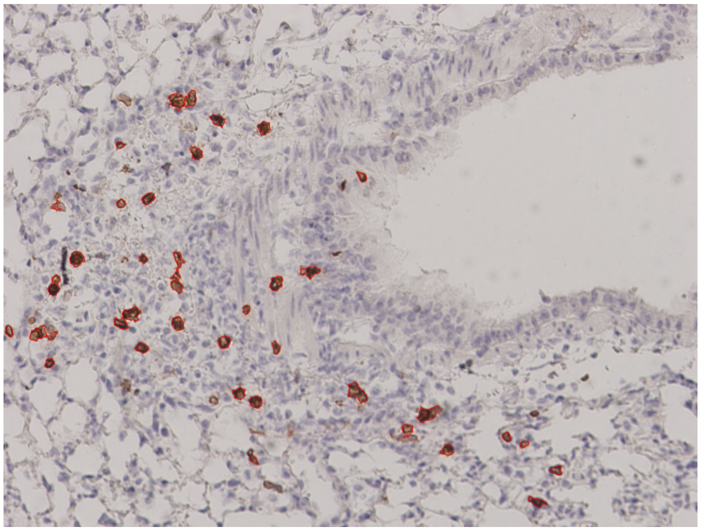

42 CD8$^+$ cells

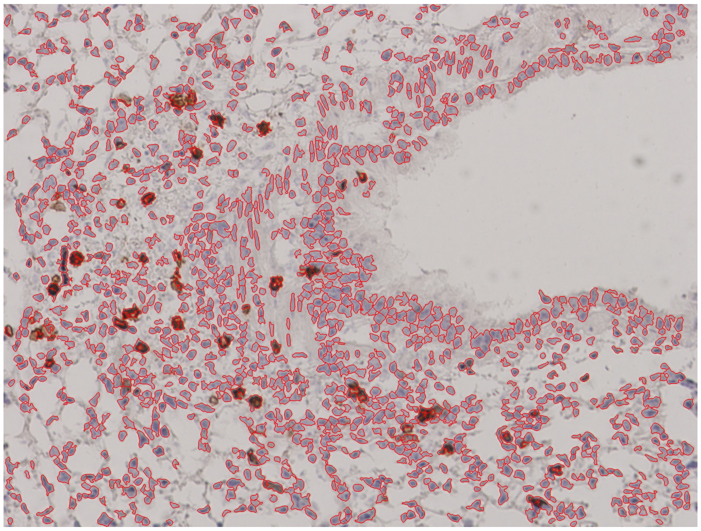

1259 nucleated cells

CD4$^+$

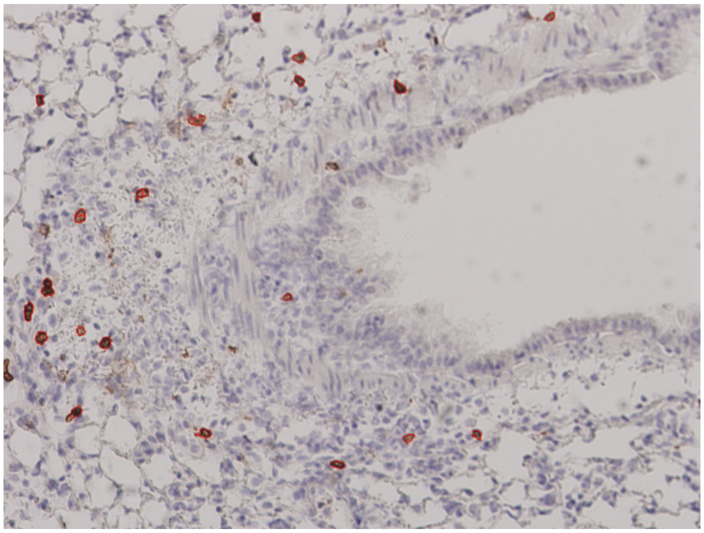

23 CD4$^+$ cells

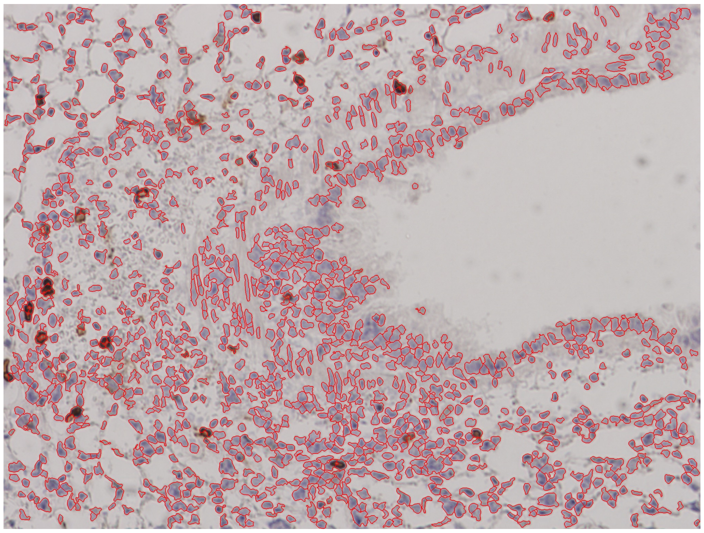

1241 nucleated cells

Supplemental Figure 7
Page 15/29

PBS Lung 3-2
2 dpi

CD8$^+$/CD4$^+$ Cell Annotations | Nucleated Cell Annotations

CD8$^+$

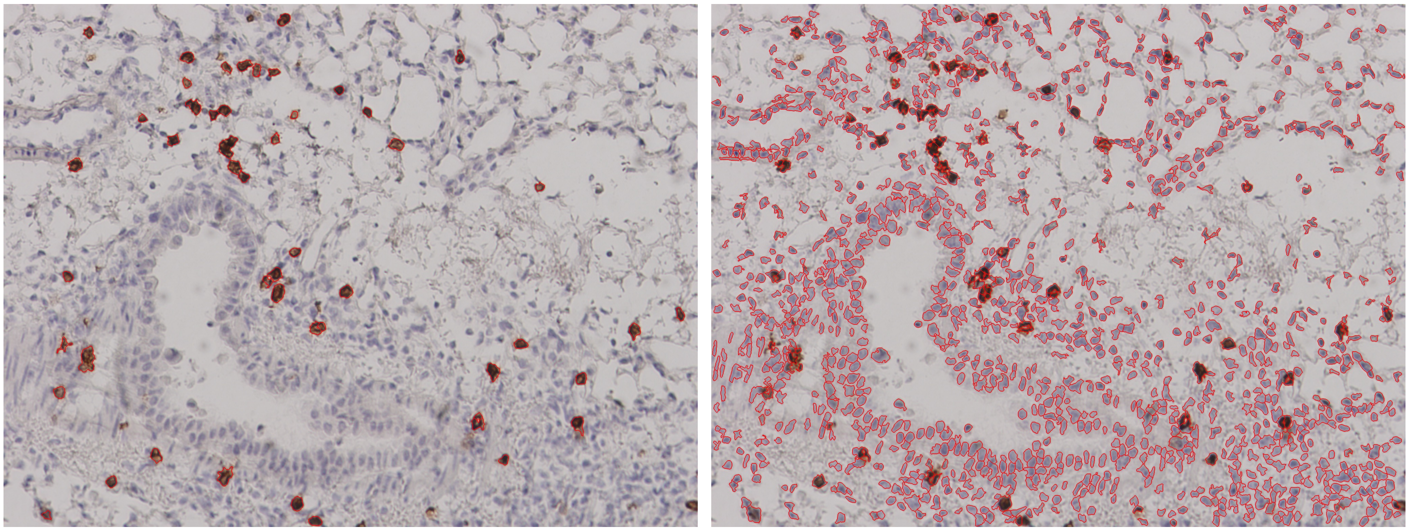

50 CD8$^+$ cells | 1208 nucleated cells

CD4$^+$

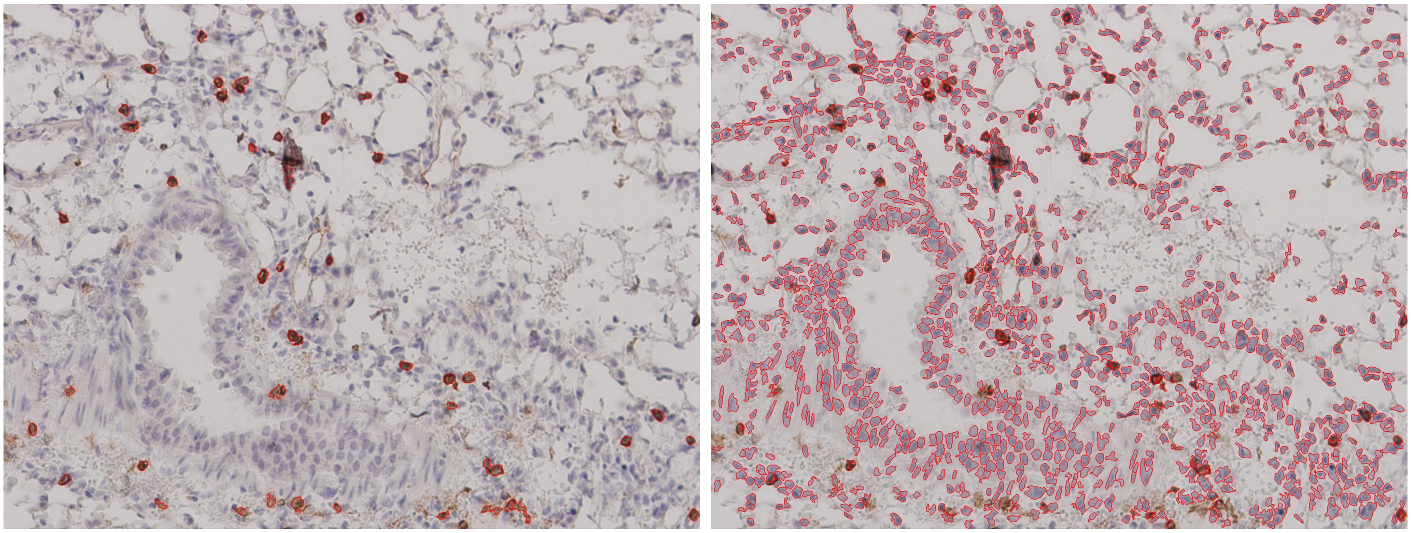

48 CD4$^+$ cells | 1205 nucleated cells

Supplemental Figure 7
Page 16/29

MIT-T-COVID Lung 1-1
7 dpi

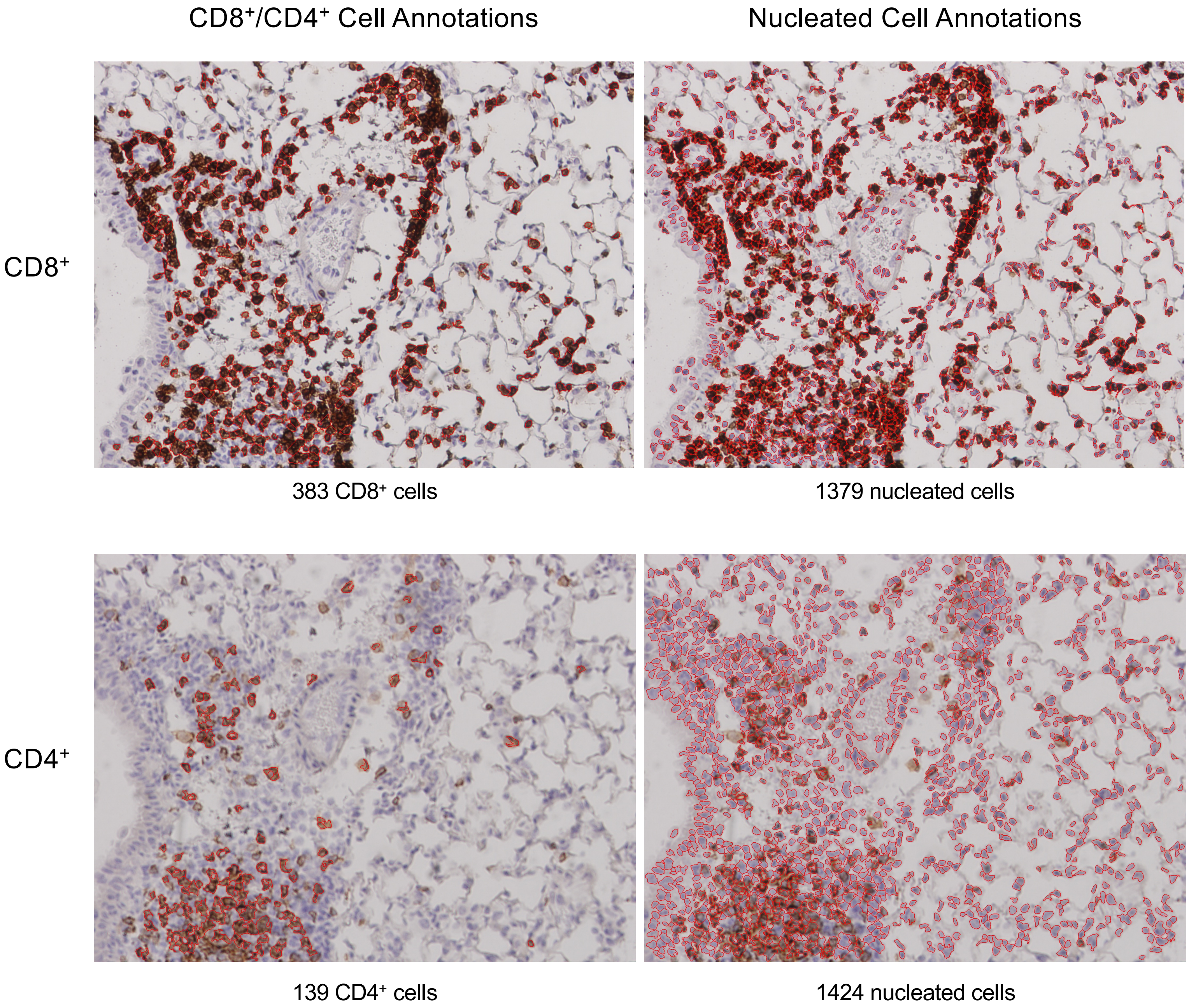

MIT-T-COVID Lung 1-2
7 dpi

CD8⁺/CD4⁺ Cell Annotations | Nucleated Cell Annotations

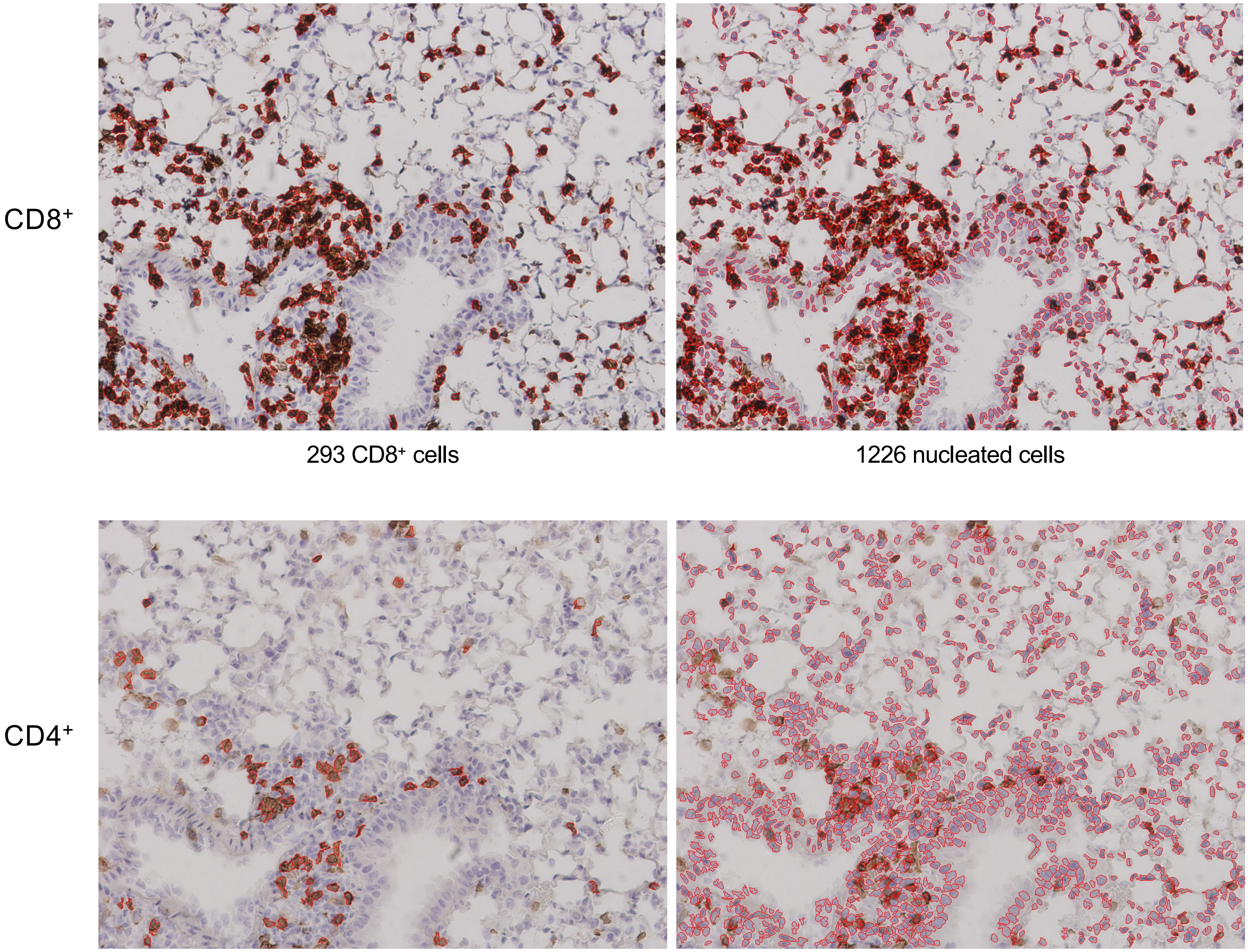


293 CD8+ cells | 1226 nucleated cells

78 CD4+ cells | 1209 nucleated cells

MIT-T-COVID Lung 2-1
7 dpi

$CD8^+$/$CD4^+$ Cell Annotations

Nucleated Cell Annotations

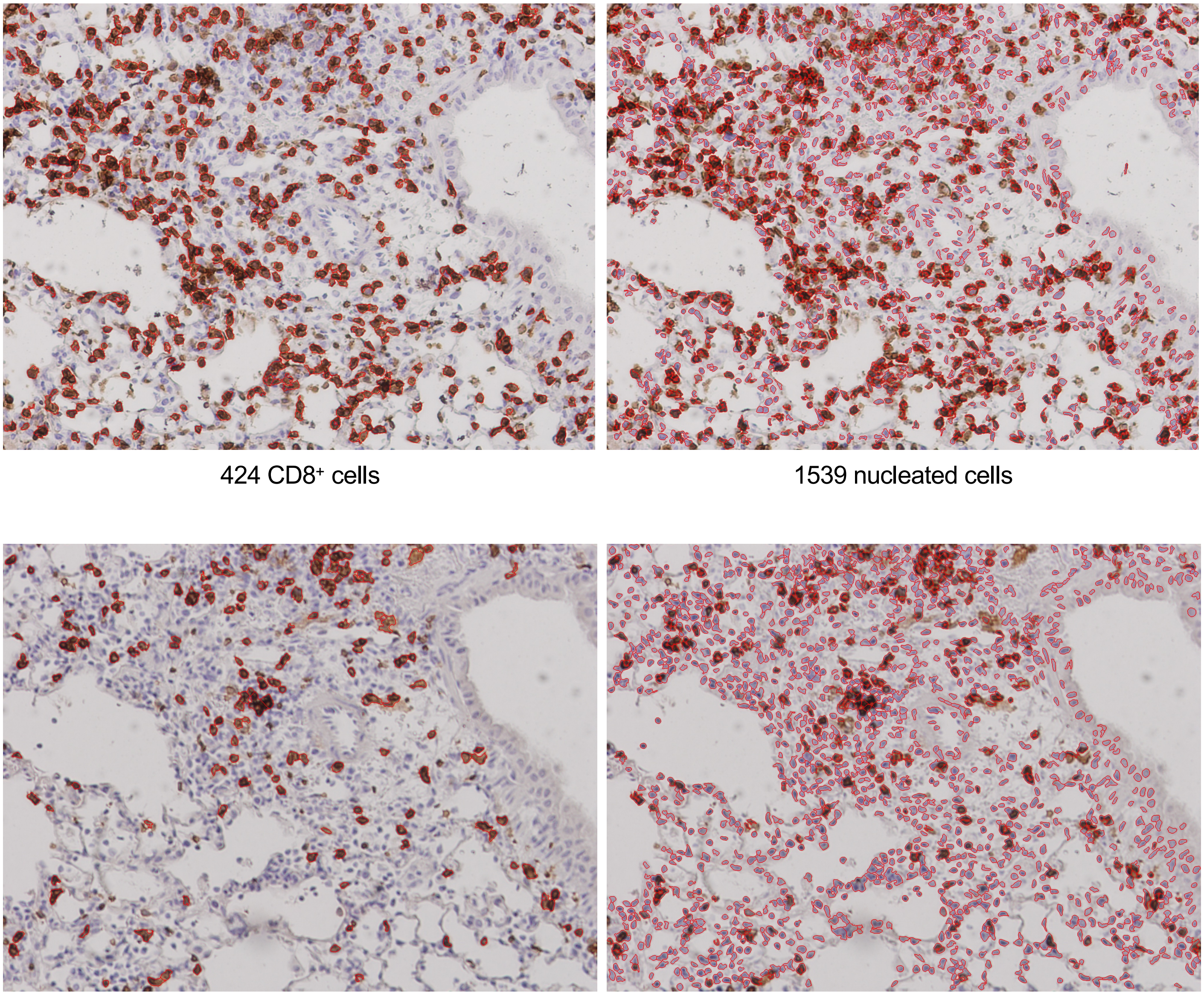

$CD8^+$

424 $CD8^+$ cells

1539 nucleated cells

$CD4^+$

152 $CD4^+$ cells

1446 nucleated cells

MIT-T-COVID Lung 2-2
7 dpi

CD8$^+$/CD4$^+$ Cell Annotations | Nucleated Cell Annotations

CD8$^+$

CD4$^+$

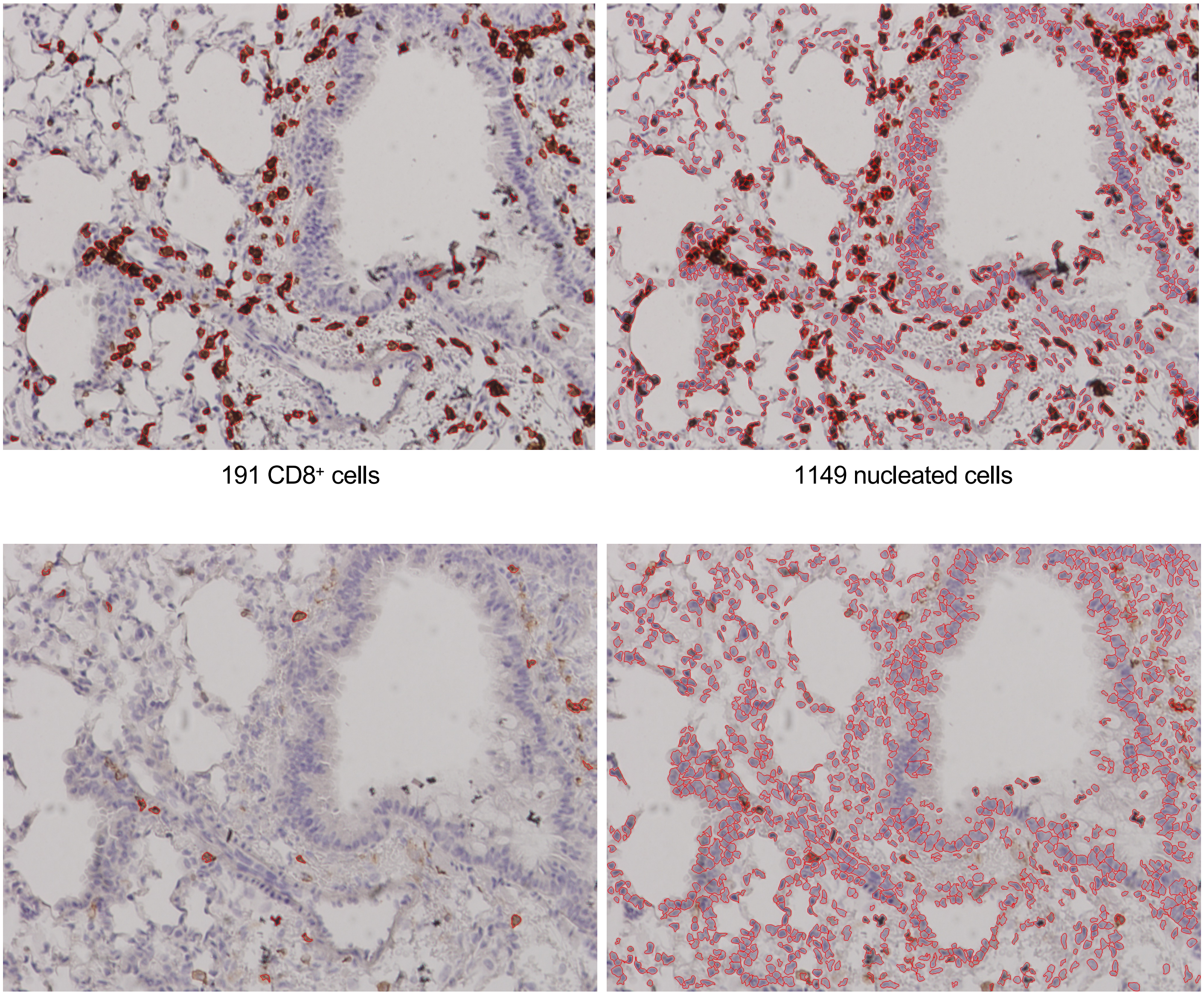

191 CD8$^+$ cells | 1149 nucleated cells

17 CD4$^+$ cells | 1052 nucleated cells

#### Supplemental Figure 7


##### Pfizer/BNT Lung 1-1
7 dpi

| | CD8$^+$/CD4$^+$ Cell Annotations | Nucleated Cell Annotations |
|---|---|---|

CD8$^+$

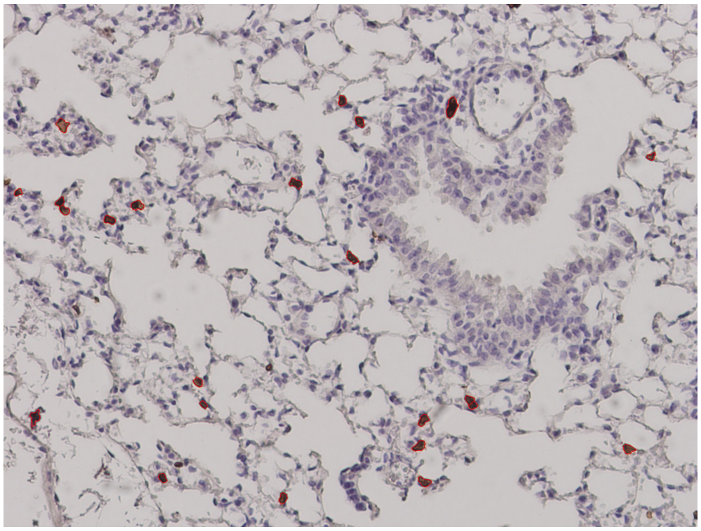

23 CD8$^+$ cells

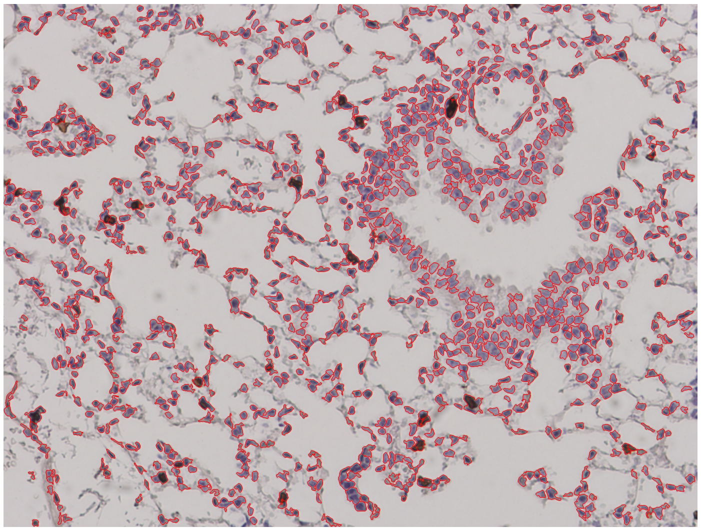

1093 nucleated cells

CD4$^+$

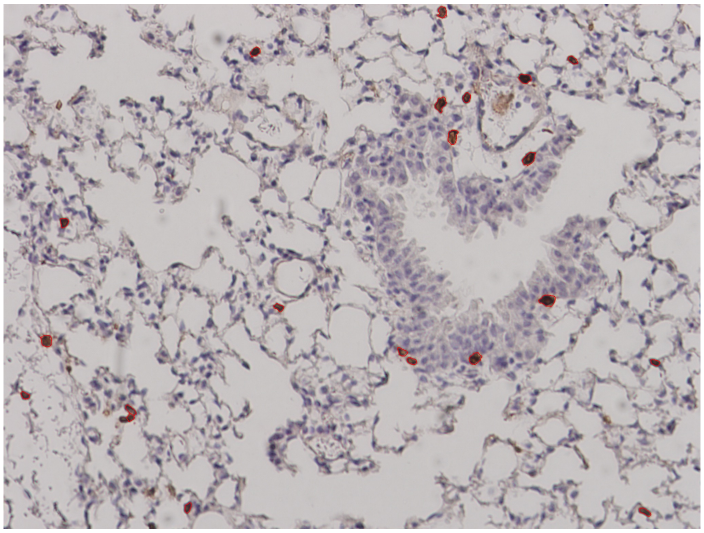

21 CD4$^+$ cells

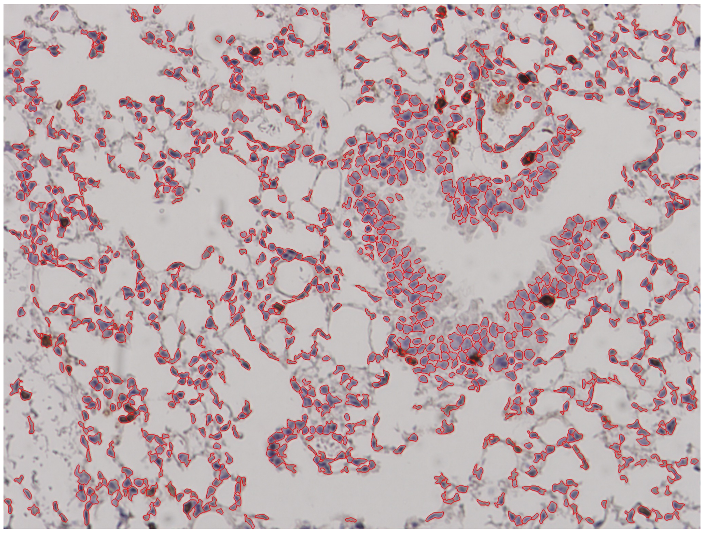

1119 nucleated cells

Supplemental Figure 7


Pfizer/BNT Lung 1-2
7 dpi

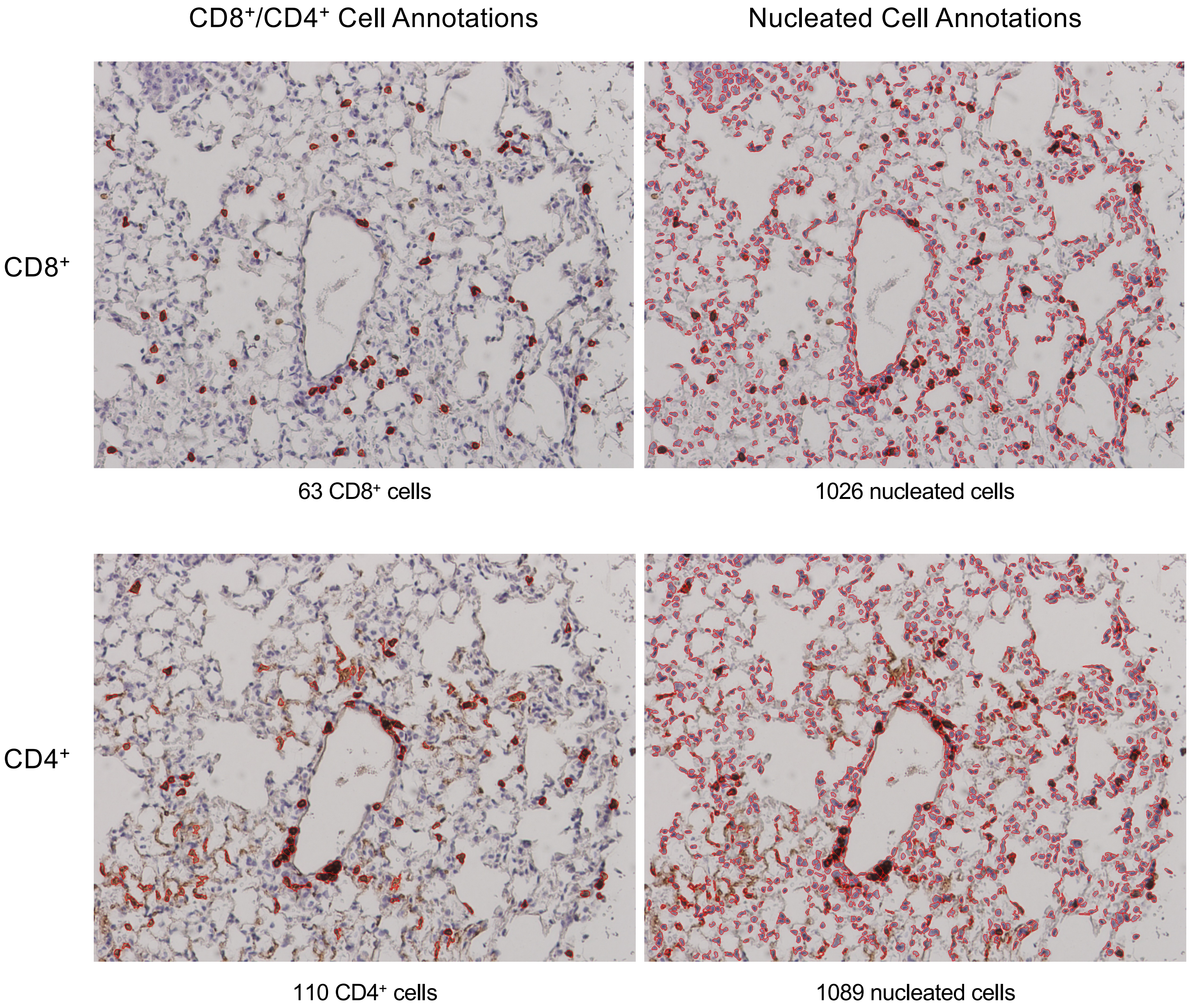

Supplemental Figure 7


Pfizer/BNT Lung 1-3
7 dpi

CD8$^+$/CD4$^+$ Cell Annotations | Nucleated Cell Annotations

CD8$^+$

CD4$^+$

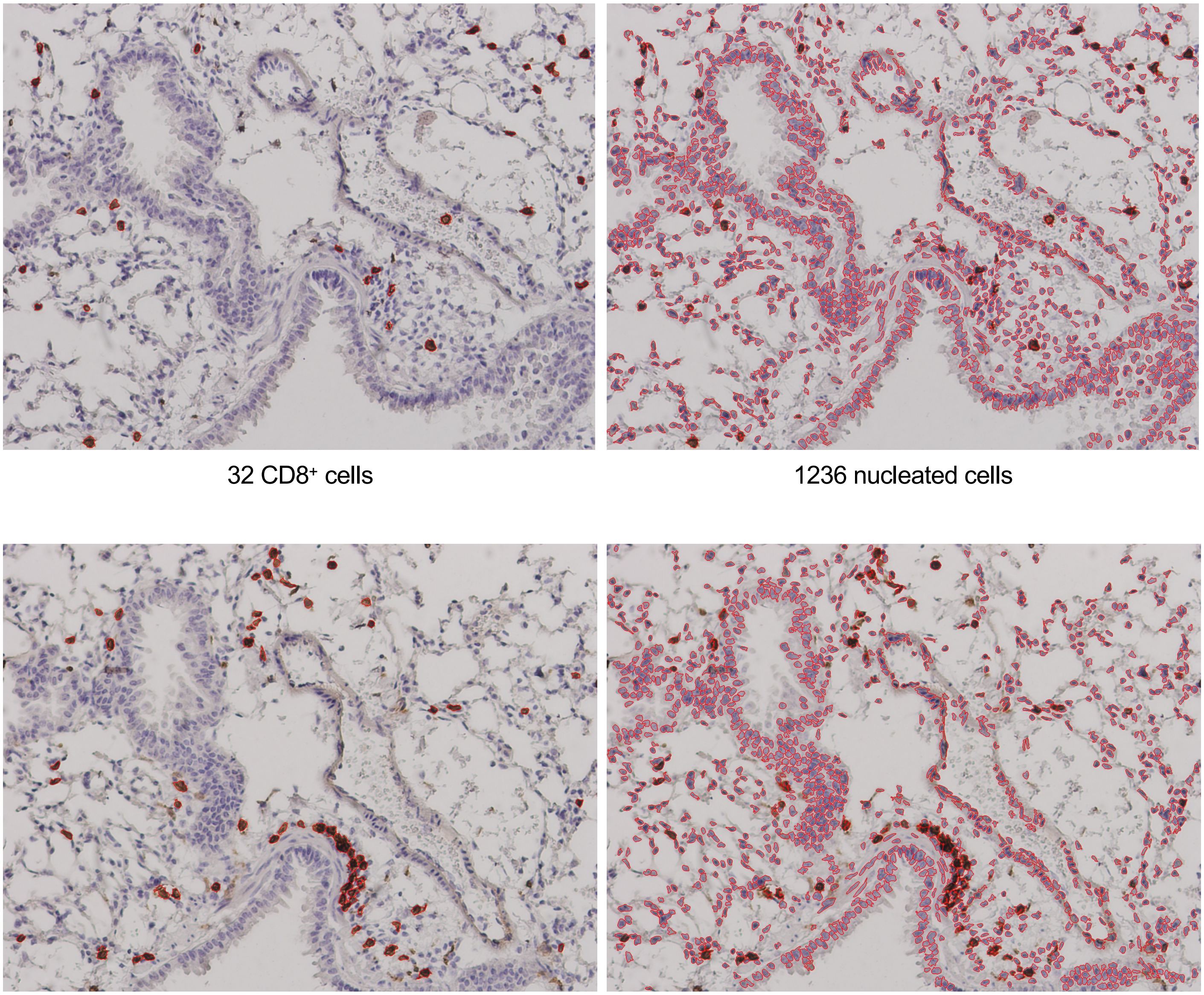

32 CD8$^+$ cells | 1236 nucleated cells

63 CD4$^+$ cells | 1153 nucleated cells

Supplemental Figure 7
Page 23/29

Pfizer/BNT Lung 2-1
7 dpi

CD8$^+$/CD4$^+$ Cell Annotations

Nucleated Cell Annotations

CD8$^+$

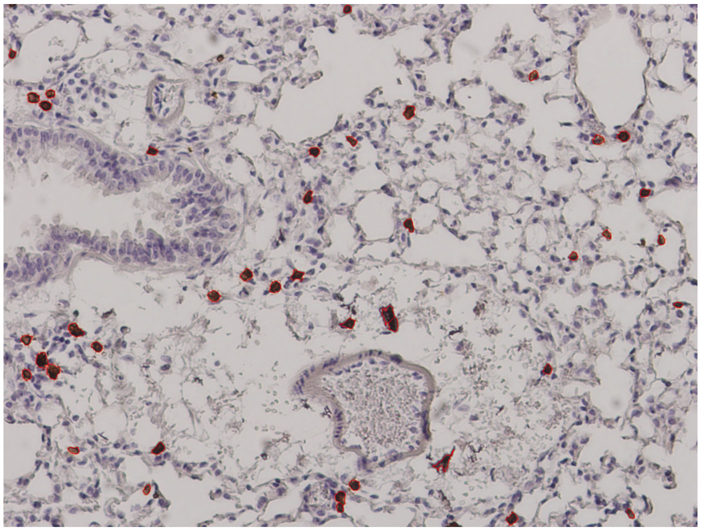

42 CD8$^+$ cells

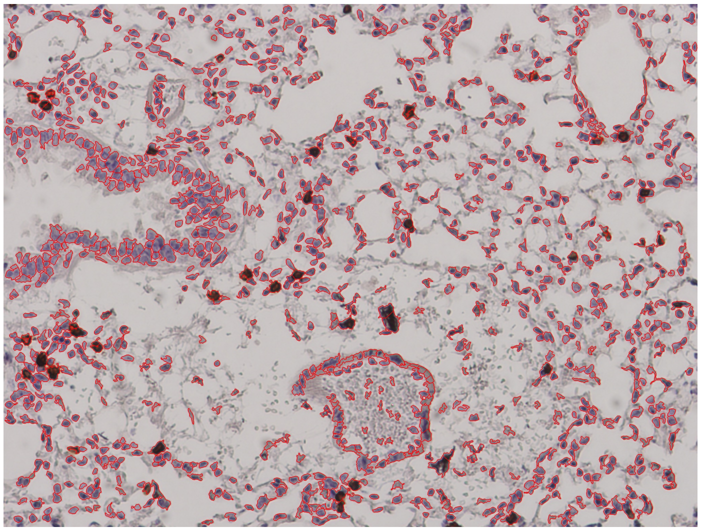

1108 nucleated cells

CD4$^+$

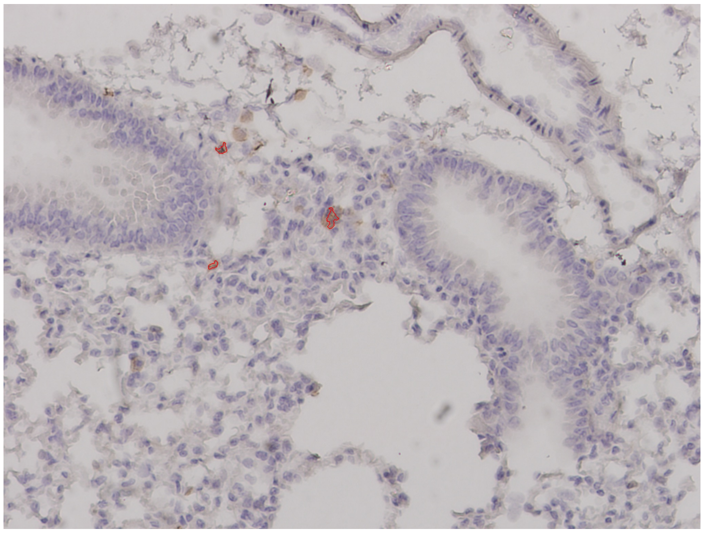

3 CD4$^+$ cells

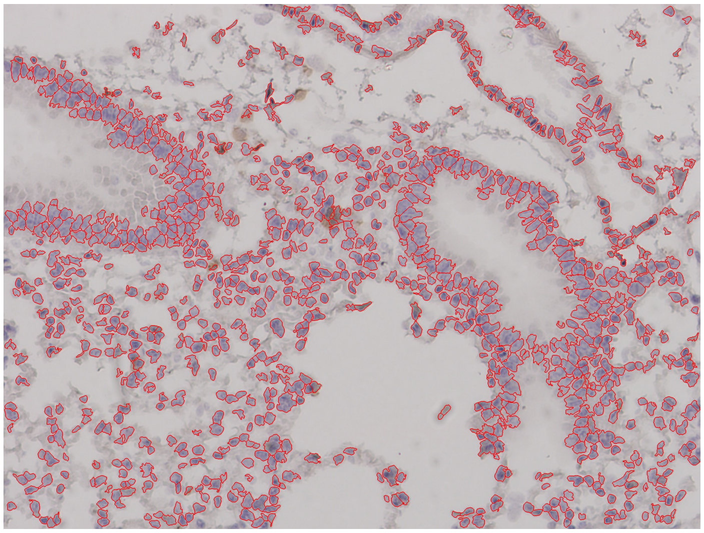

1036 nucleated cells

Supplemental Figure 7
Page 24/29

Pfizer/BNT Lung 3-1
7 dpi

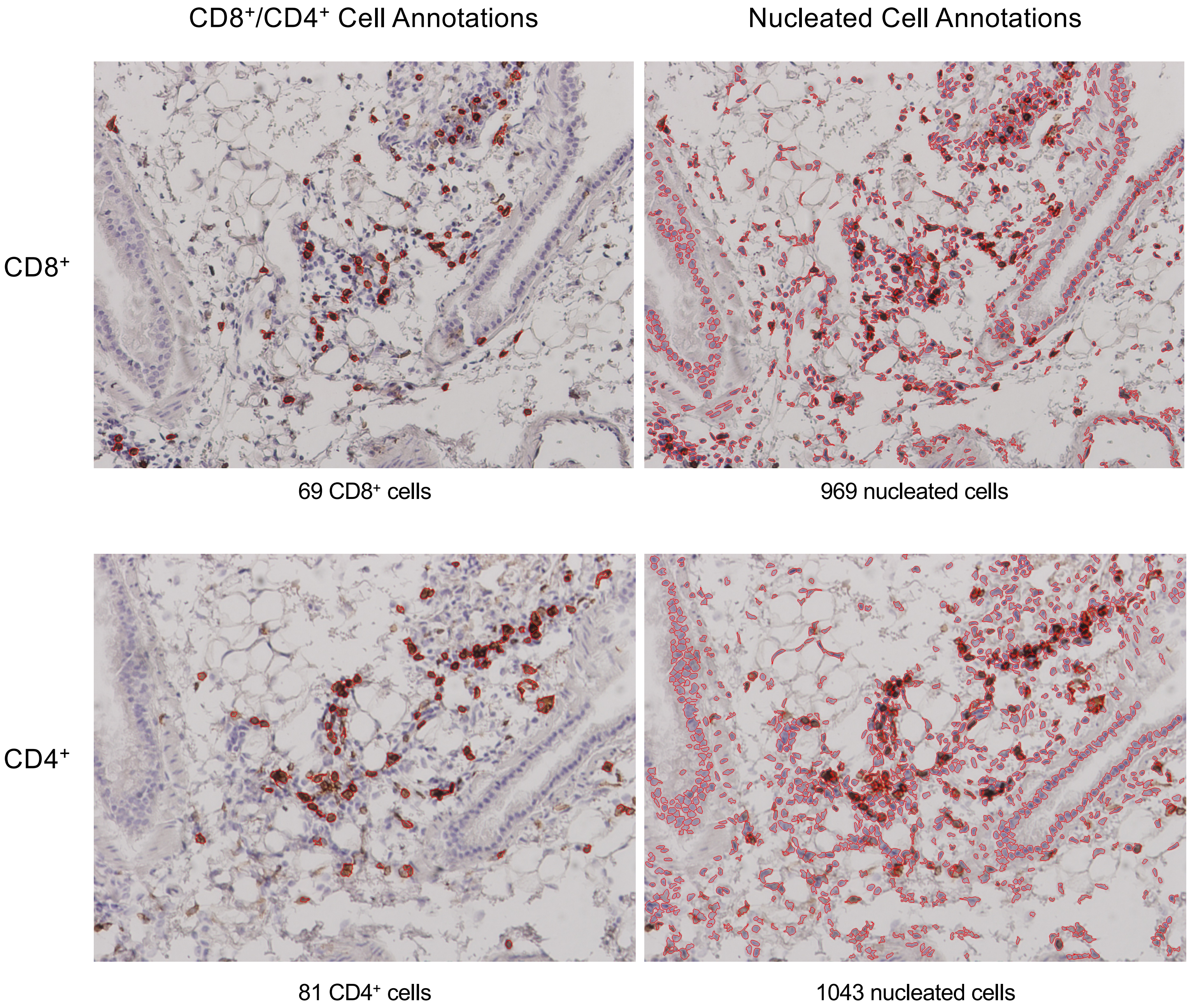

Supplemental Figure 7
Page 25/29

PBS Lung 1-1
7 dpi

CD8$^+$/CD4$^+$ Cell Annotations | Nucleated Cell Annotations

CD8$^+$

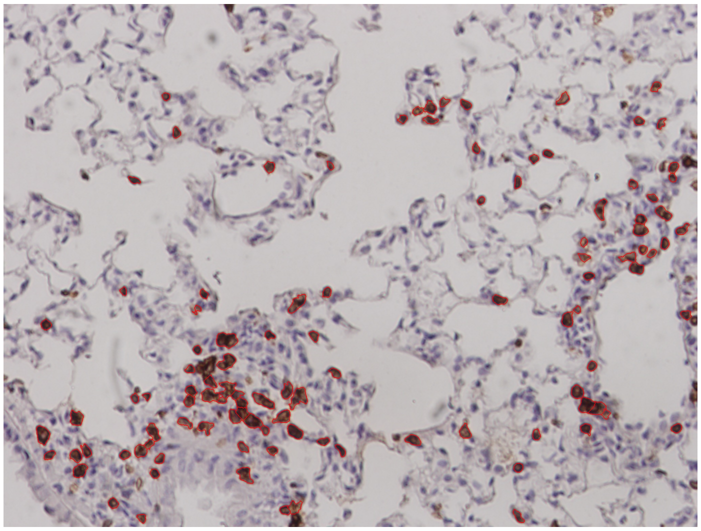

108 CD8$^+$ cells

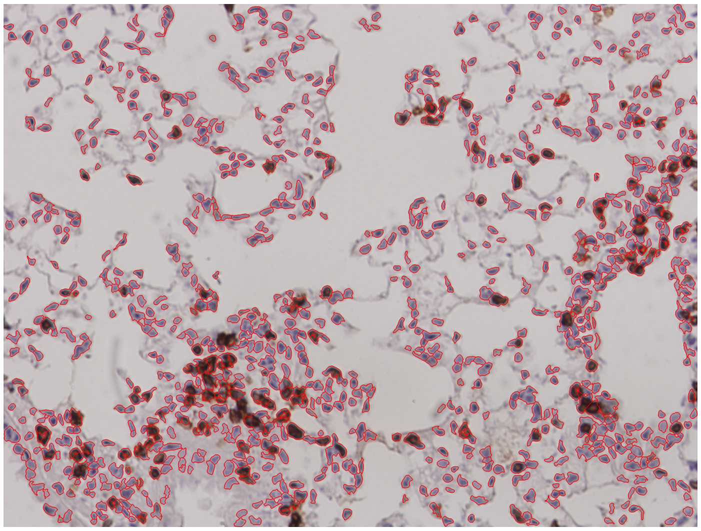

912 nucleated cells

CD4$^+$

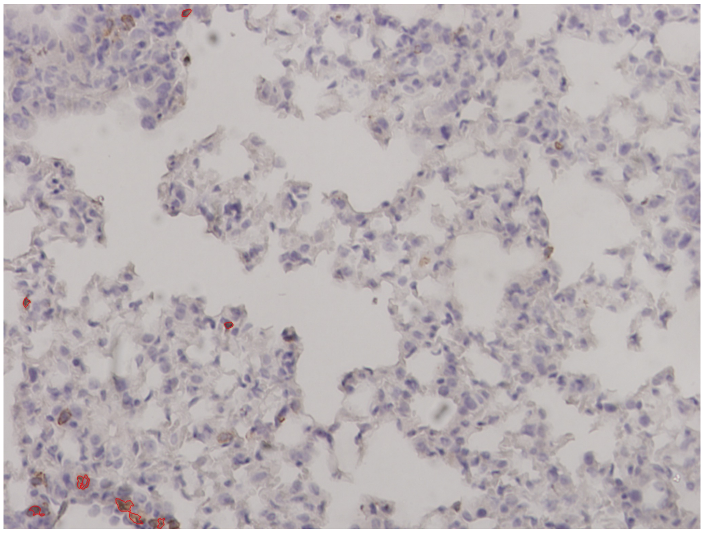

9 CD4$^+$ cells

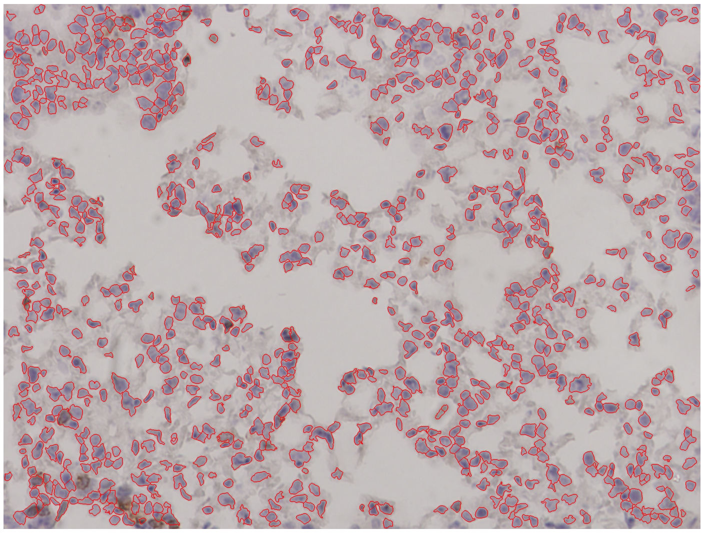

892 nucleated cells

Supplemental Figure 7
Page 26/29

PBS Lung 2-1
7 dpi

CD8⁺/CD4⁺ Cell Annotations | Nucleated Cell Annotations

CD8⁺

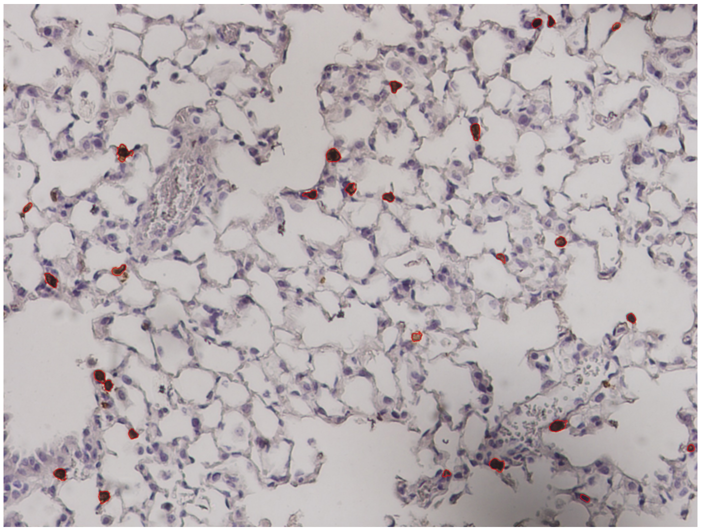

27 CD8⁺ cells

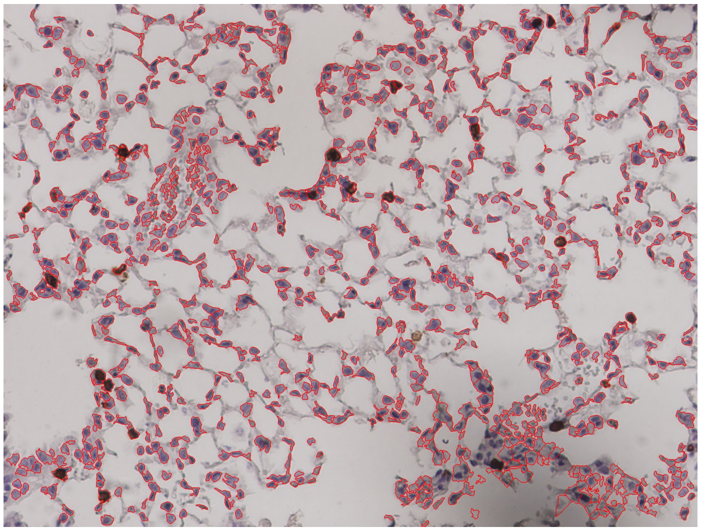

958 nucleated cells

CD4⁺

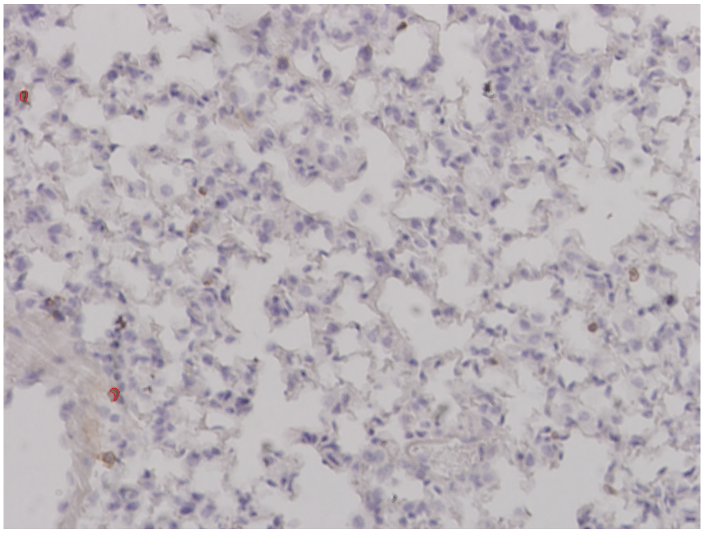

2 CD4⁺ cells

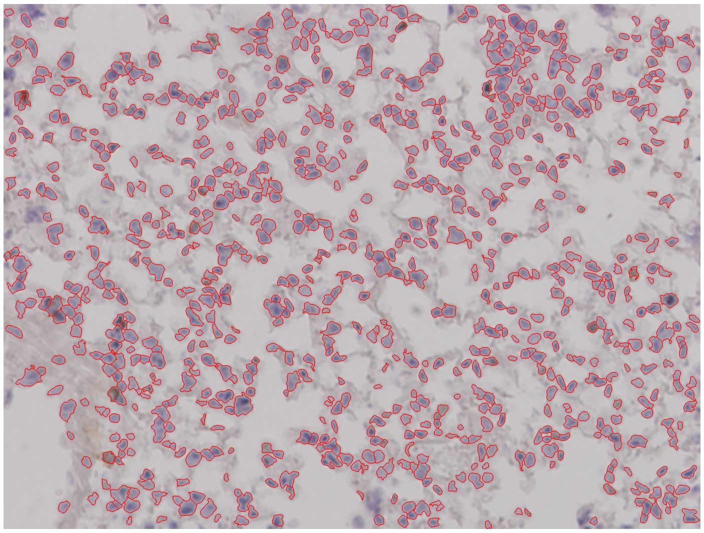

813 nucleated cells

Supplemental Figure 7


PBS Lung 2-2
7 dpi

CD8$^+$/CD4$^+$ Cell Annotations | Nucleated Cell Annotations

CD8$^+$

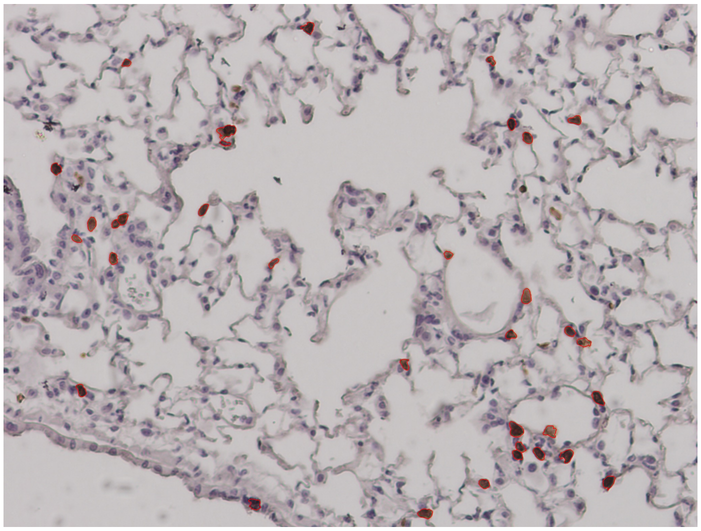

34 CD8$^+$ cells

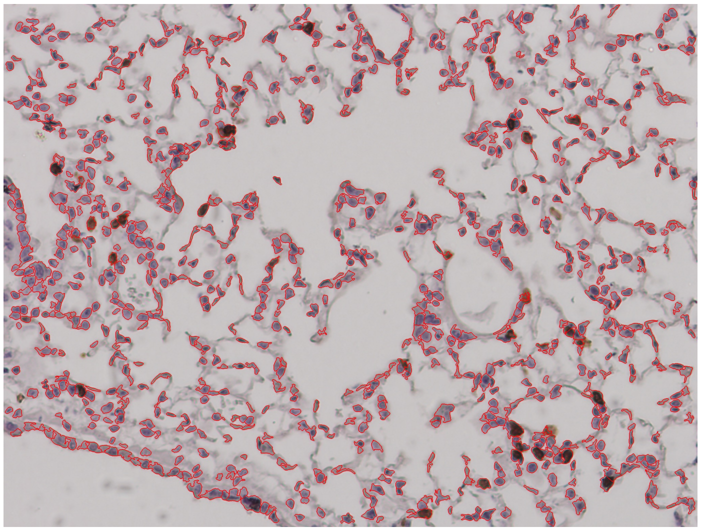

869 nucleated cells

CD4$^+$

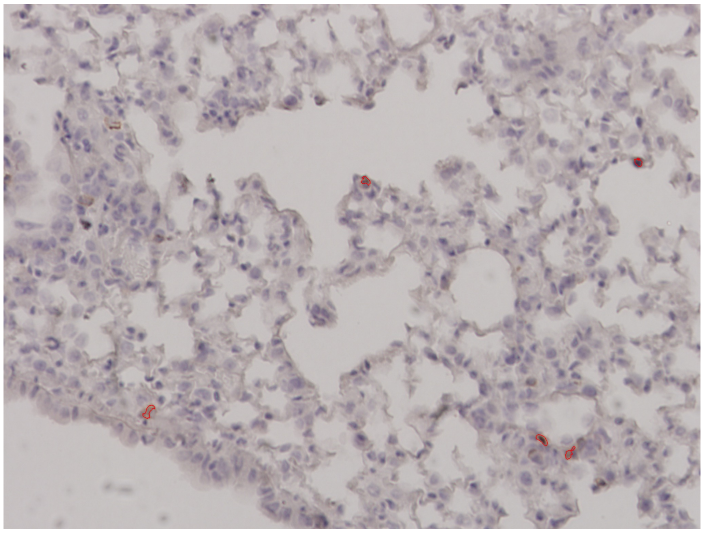

5 CD4$^+$ cells

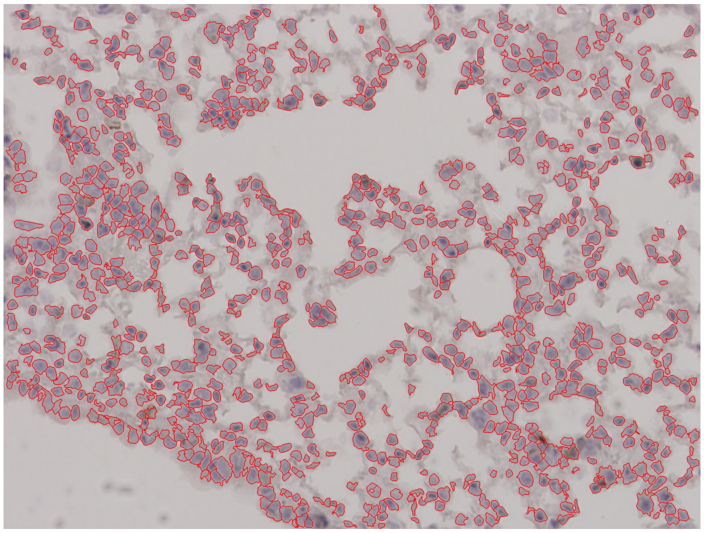

937 nucleated cells

Supplemental Figure 7


PBS Lung 4-1
7 dpi

CD8$^{+}$/CD4$^{+}$ Cell Annotations | Nucleated Cell Annotations

CD8$^{+}$

CD4$^{+}$

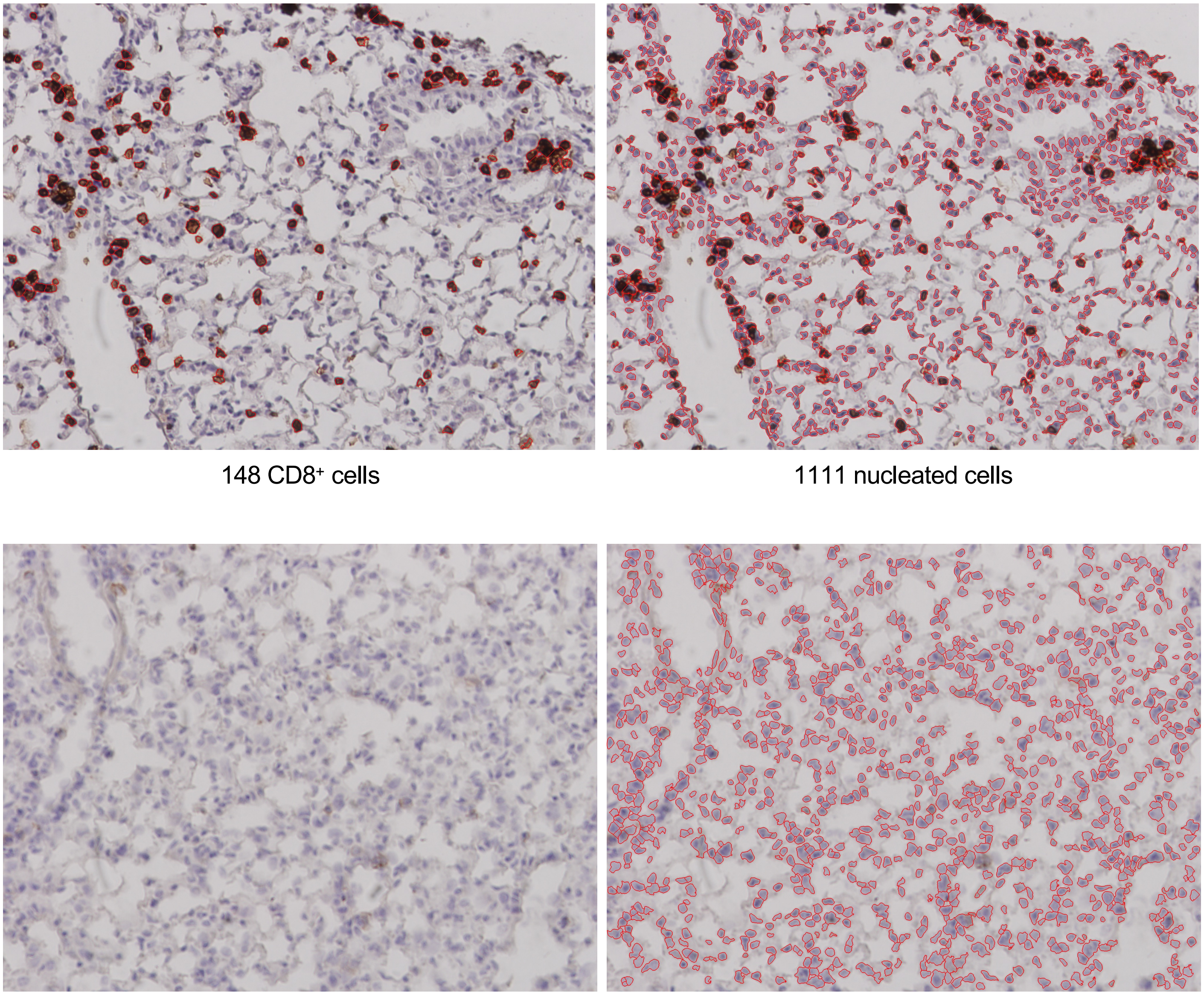

148 CD8$^{+}$ cells | 1111 nucleated cells

0 CD4$^{+}$ cells | 981 nucleated cells

Supplemental Figure 7


PBS Lung 5-1
7 dpi

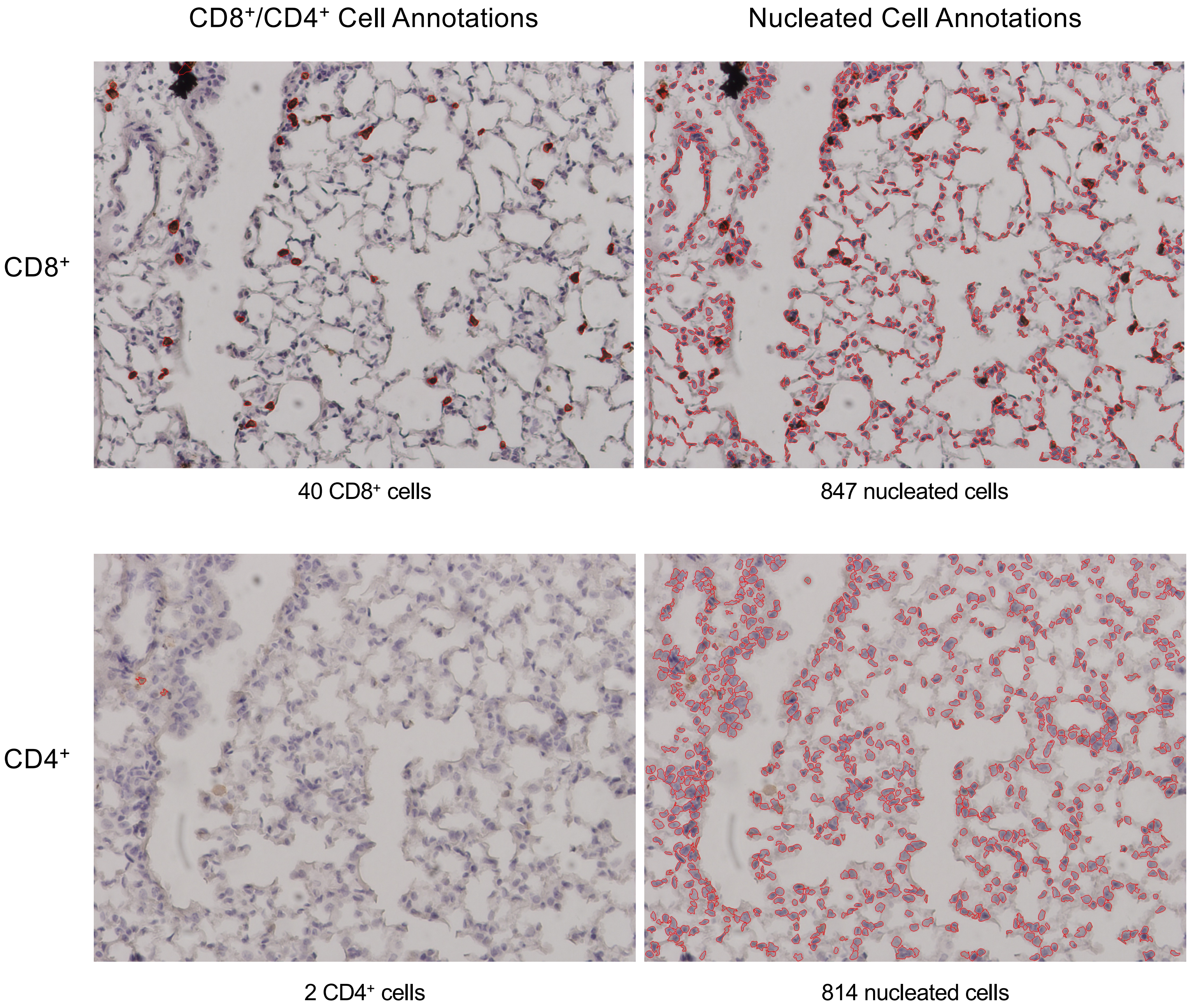
