## Supplemental Figure 11 for "A pan-variant mRNA-LNP T cell vaccine protects HLA transgenic mice from mortality after infection with SARS-CoV-2 Beta"


MIT-T-COVID Lung 1-1
Unchallenged, Female Cohort

| | CD8+/CD4+ Cell Annotations | Nucleated Cell Annotations |
|---|---|---|
| CD8+ | 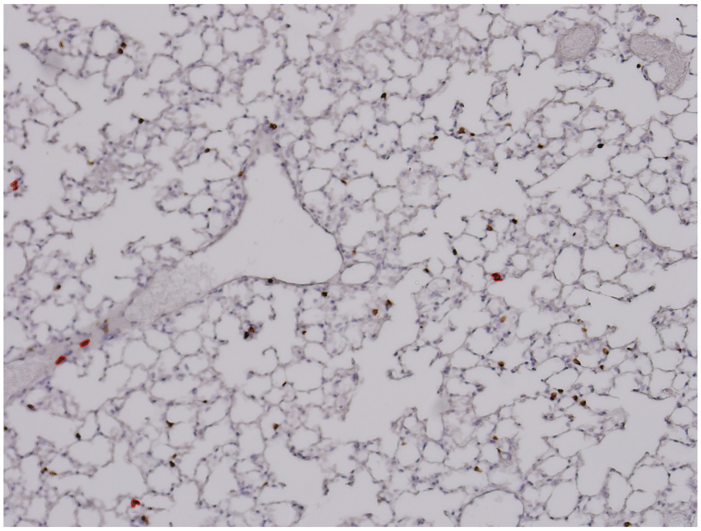 | 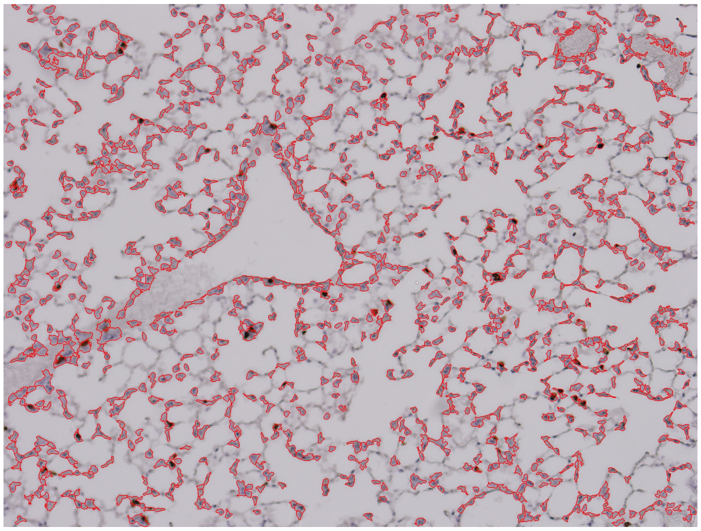 |
| | 5 CD8+ cells | 1104 nucleated cells |
| CD4+ | 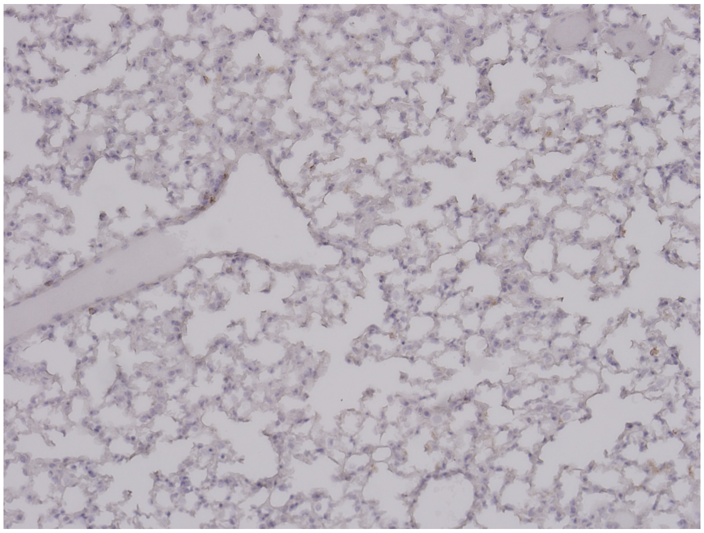 | 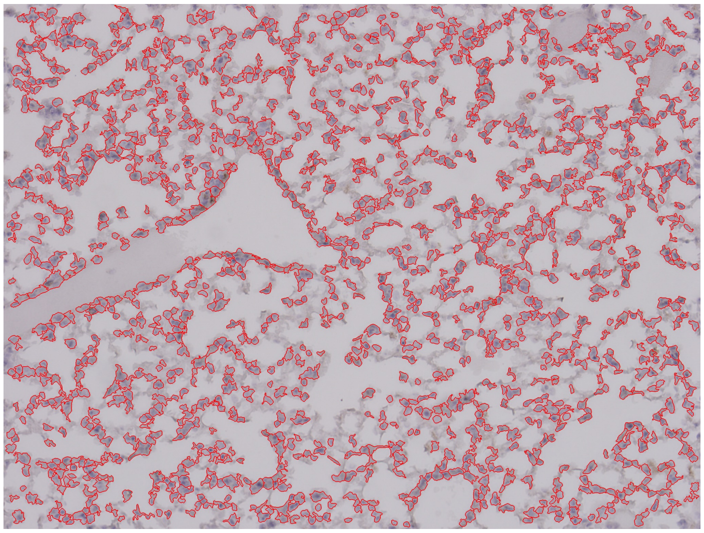 |
| | 0 CD4+ cells | 1388 nucleated cells |

#### MIT-T-COVID Lung 1-2
#### Unchallenged, Female Cohort

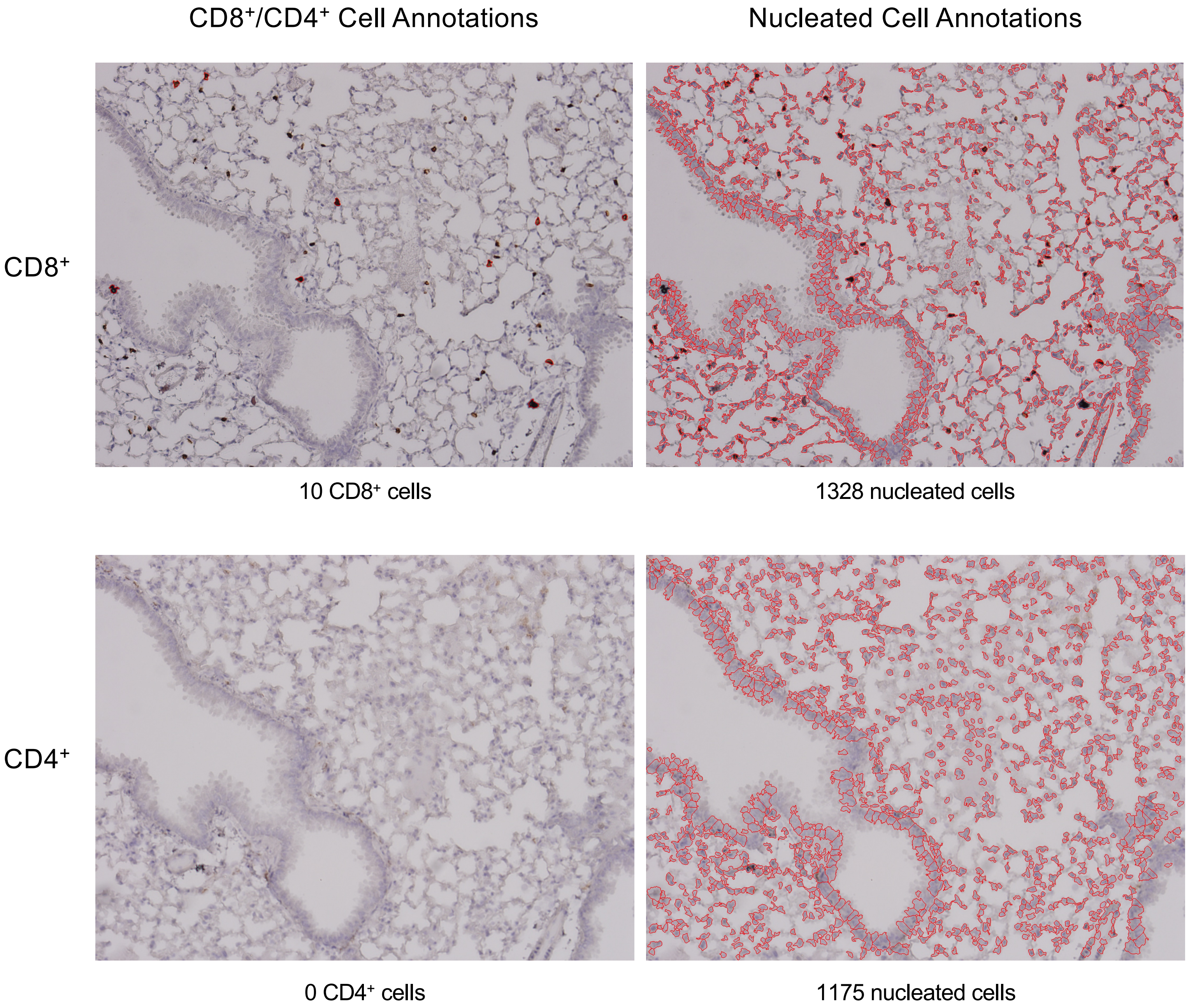

Supplemental Figure 11
Page 3/30

MIT-T-COVID Lung 2-1
Unchallenged, Female Cohort

CD8$^{+}$/CD4$^{+}$ Cell Annotations | Nucleated Cell Annotations

CD8$^{+}$

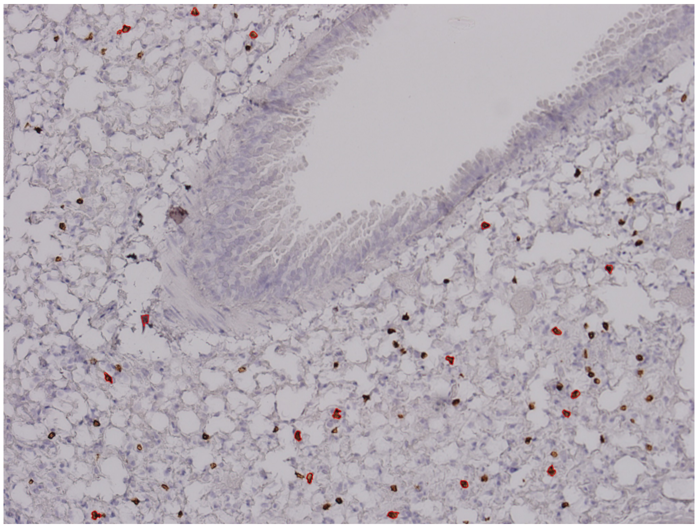

18 CD8$^{+}$ cells

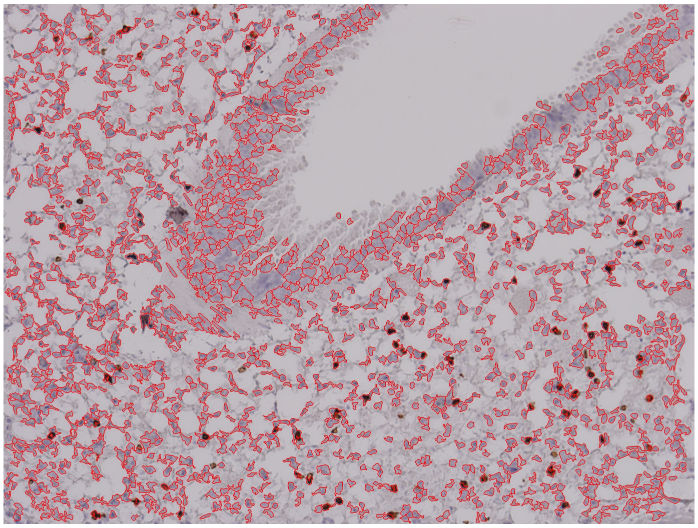

1481 nucleated cells

CD4$^{+}$

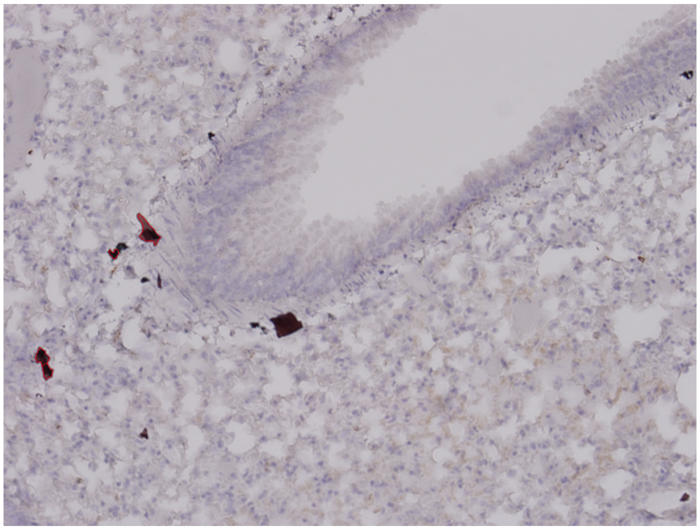

4 CD4$^{+}$ cells

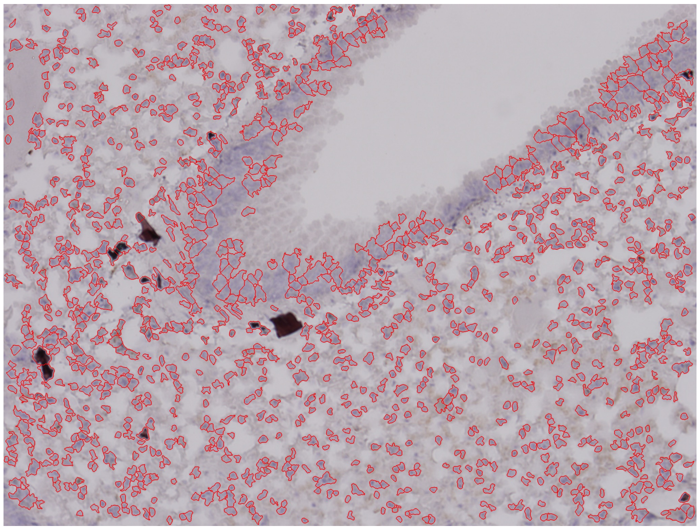

1190 nucleated cells

Supplemental Figure 11
Page 4/30

MIT-T-COVID Lung 2-2
Unchallenged, Female Cohort

CD8$^+$/CD4$^+$ Cell Annotations | Nucleated Cell Annotations

CD8$^+$

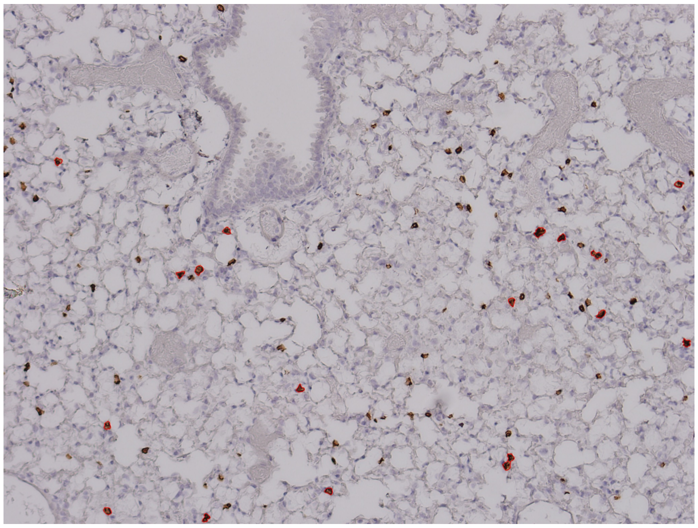

18 CD8$^+$ cells

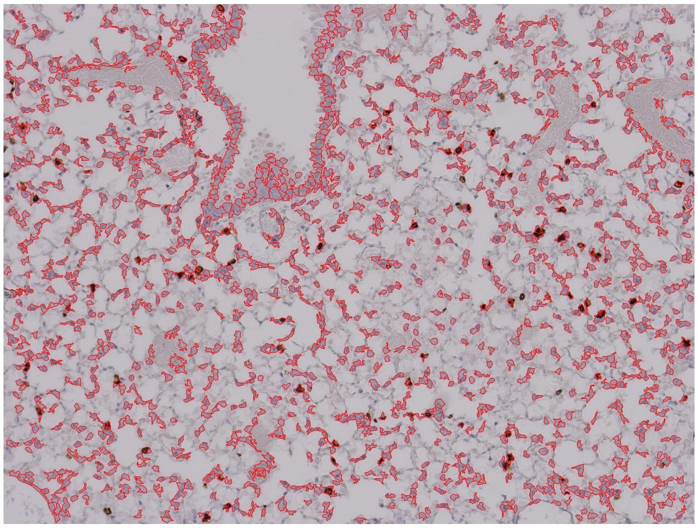

1291 nucleated cells

CD4$^+$

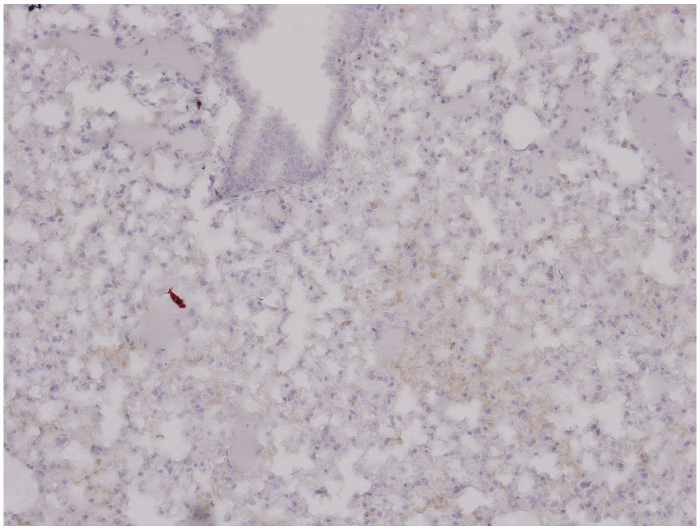

1 CD4$^+$ cells

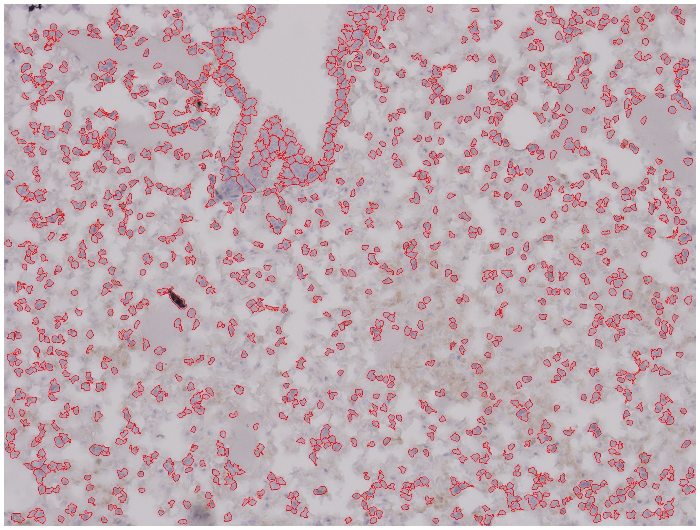

1132 nucleated cells

MIT-T-COVID Lung 2-3
Unchallenged, Female Cohort

| | CD8⁺/CD4⁺ Cell Annotations | Nucleated Cell Annotations |
|---|---|---|
| CD8⁺ | 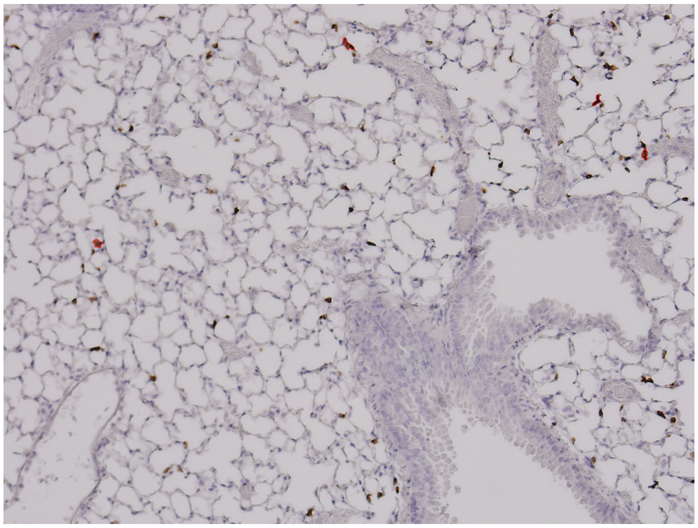 4 CD8⁺ cells | 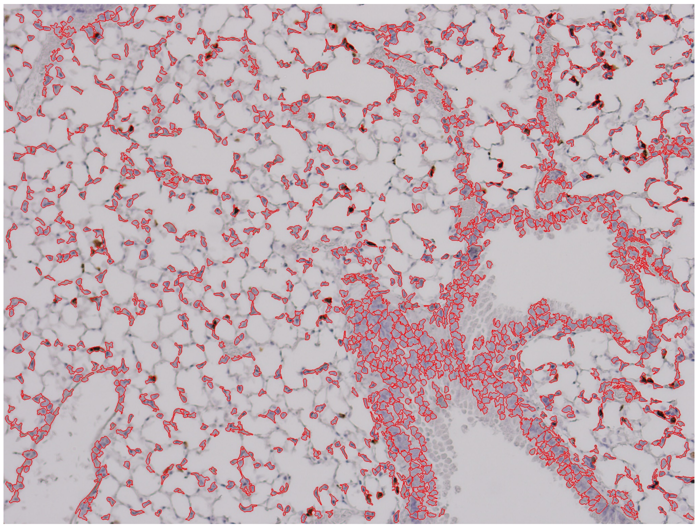 1284 nucleated cells |
| CD4⁺ | 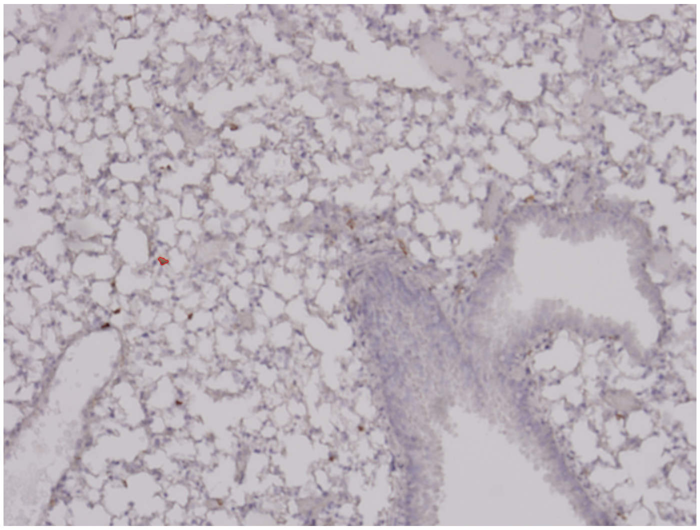 1 CD4⁺ cells | 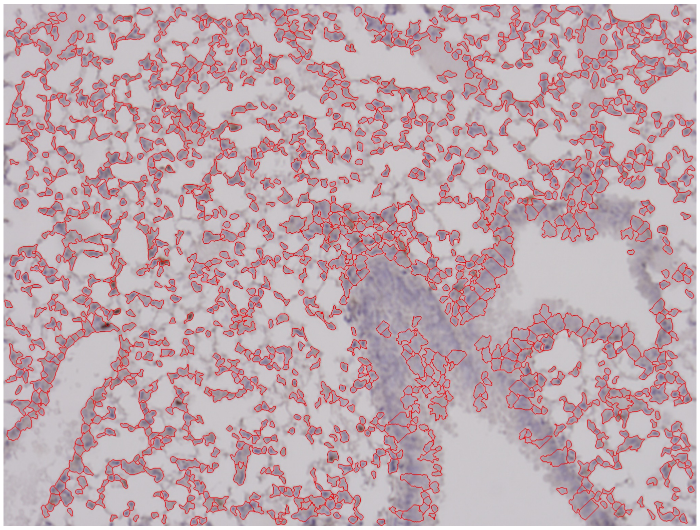 1368 nucleated cells |

MIT-T-COVID Lung 2-4
Unchallenged, Female Cohort

CD8$^+$/CD4$^+$ Cell Annotations | Nucleated Cell Annotations

CD8$^+$

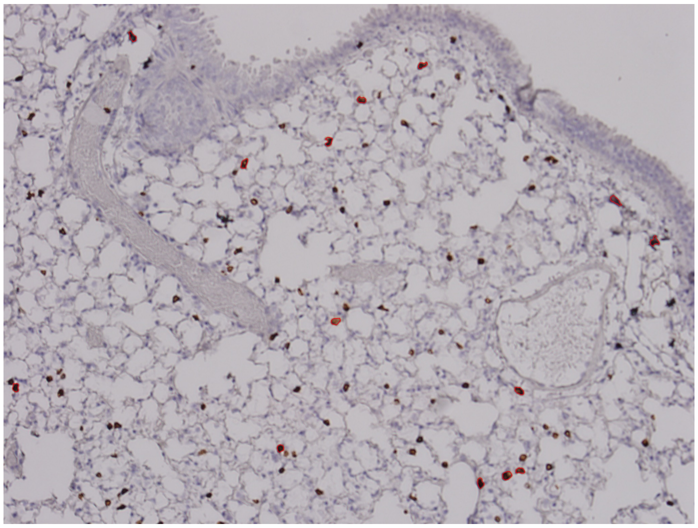

14 CD8$^+$ cells

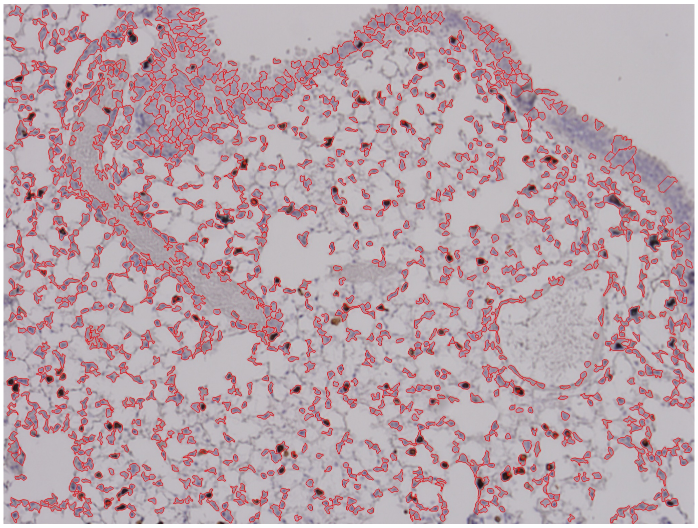

1328 nucleated cells

CD4$^+$

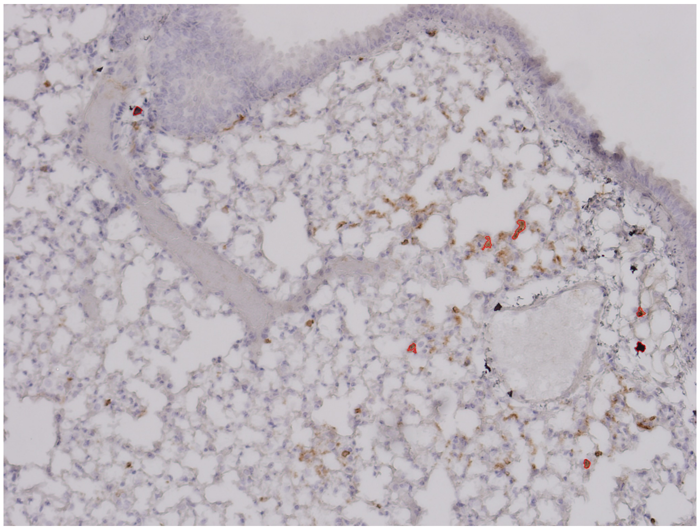

7 CD4$^+$ cells

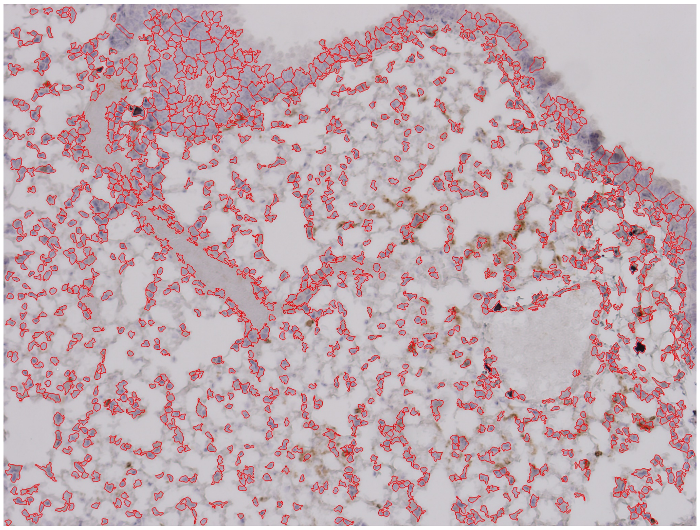

1297 nucleated cells

### MIT-T-COVID Lung 3-1
#### Unchallenged, Female Cohort

CD8$^+$/CD4$^+$ Cell Annotations | Nucleated Cell Annotations

CD8$^+$

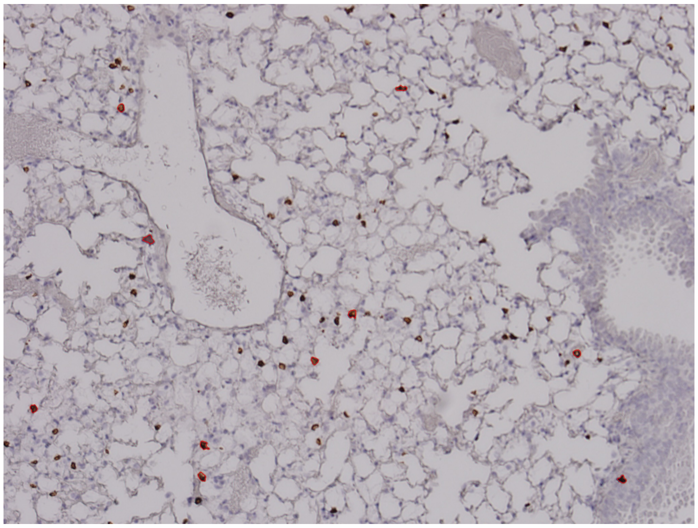

10 CD8$^+$ cells

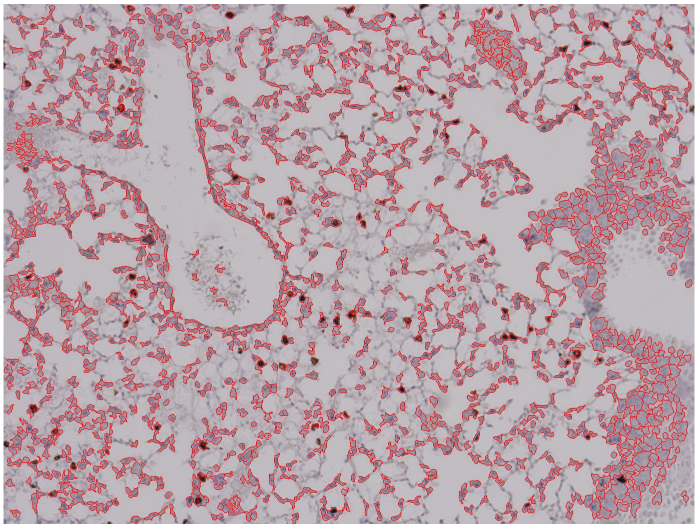

1458 nucleated cells

CD4$^+$

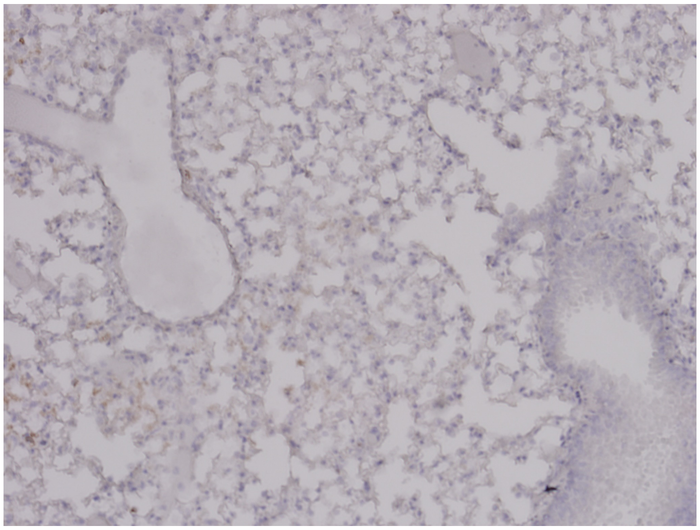

0 CD4$^+$ cells

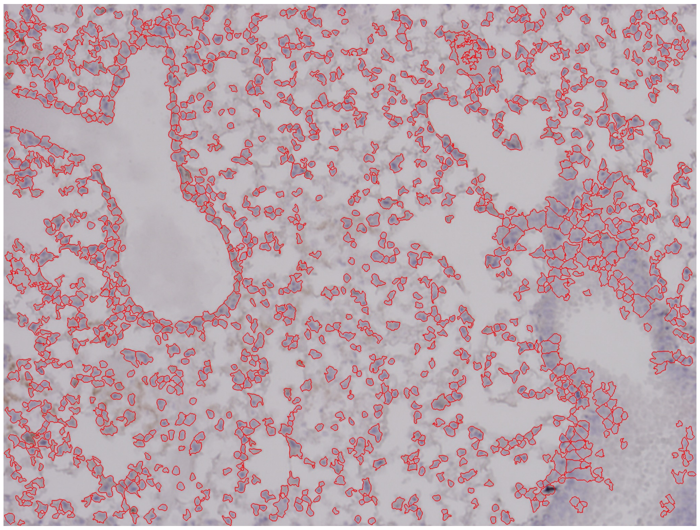

1272 nucleated cells

Supplemental Figure 11
Page 8/30

MIT-T-COVID Lung 3-2
Unchallenged, Female Cohort

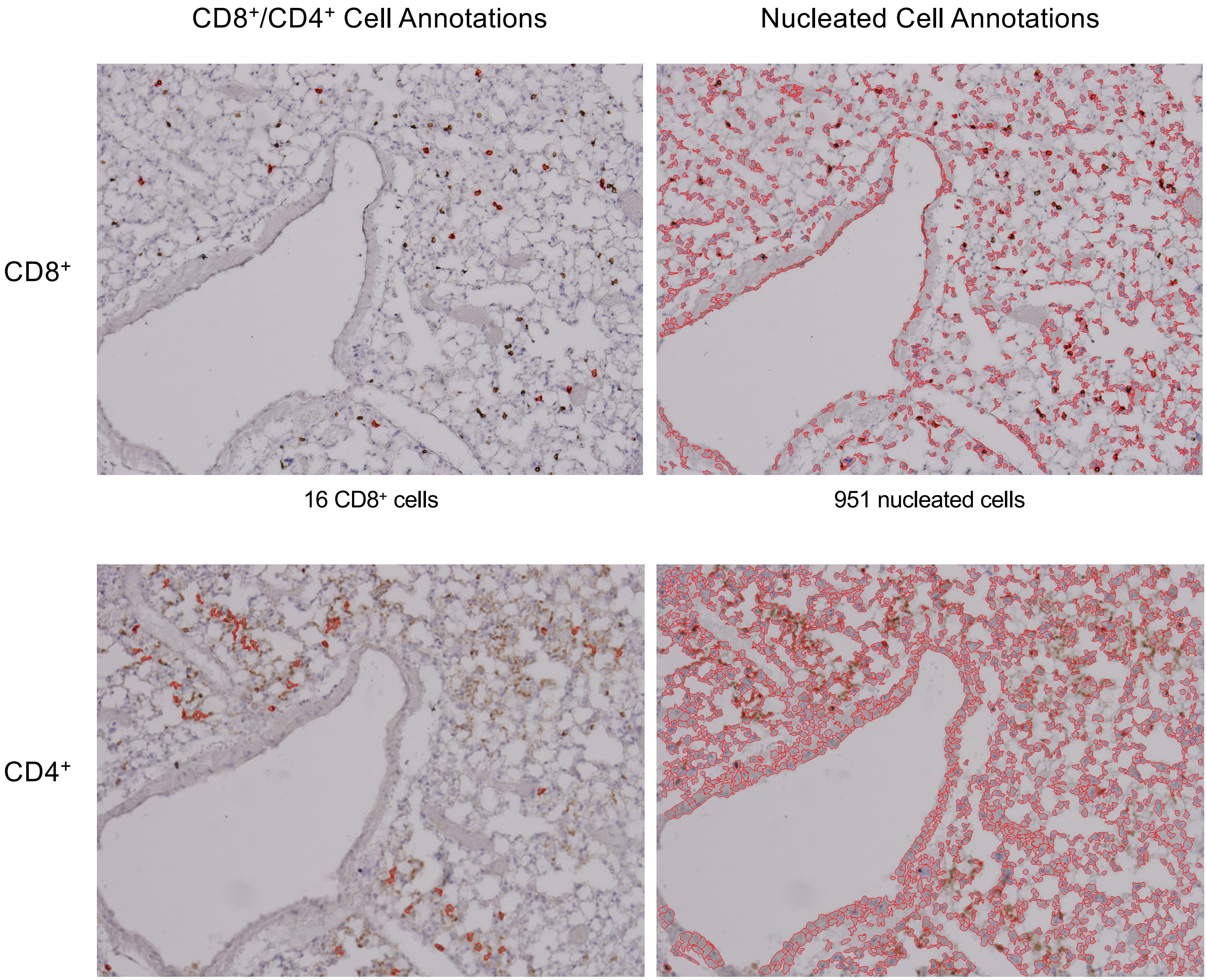


16 $CD8^+$ cells

951 nucleated cells

46 $CD4^+$ cells

1484 nucleated cells

#### MIT-T-COVID Lung 3-3
#### Unchallenged, Female Cohort

CD8$^+$/CD4$^+$ Cell Annotations | Nucleated Cell Annotations

CD8$^+$

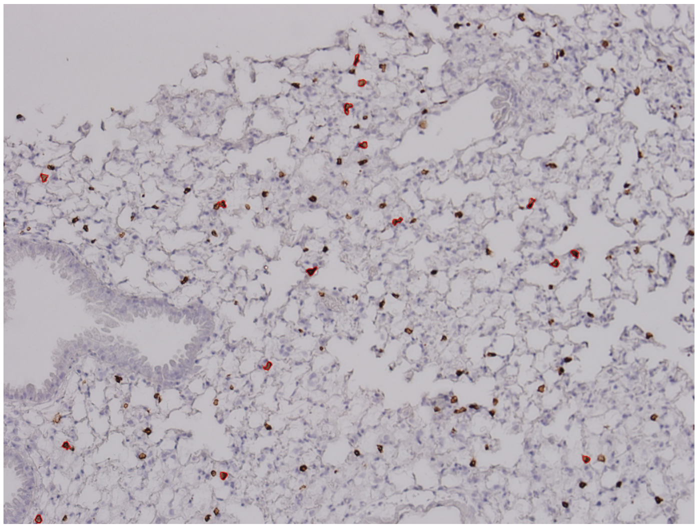

16 CD8$^+$ cells

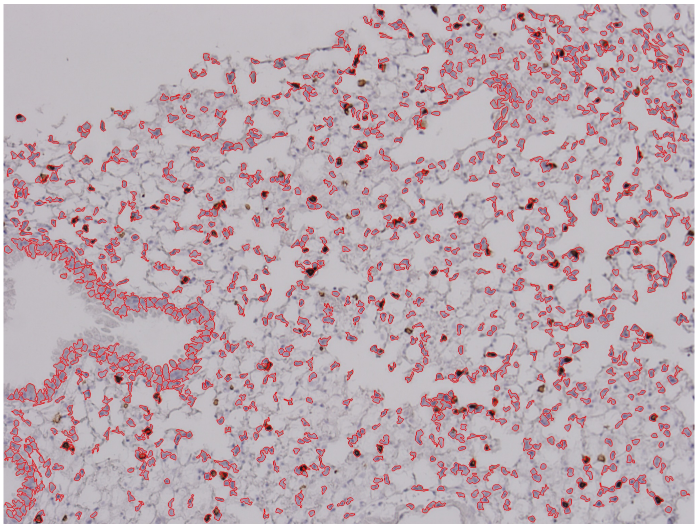

1115 nucleated cells

CD4$^+$

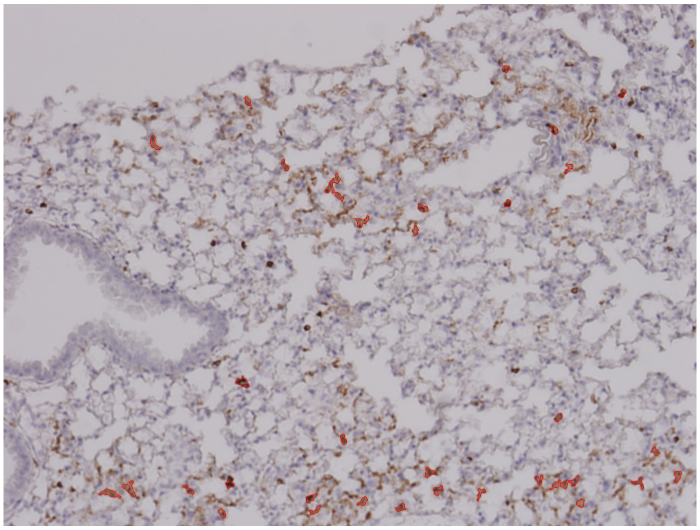

37 CD4$^+$ cells

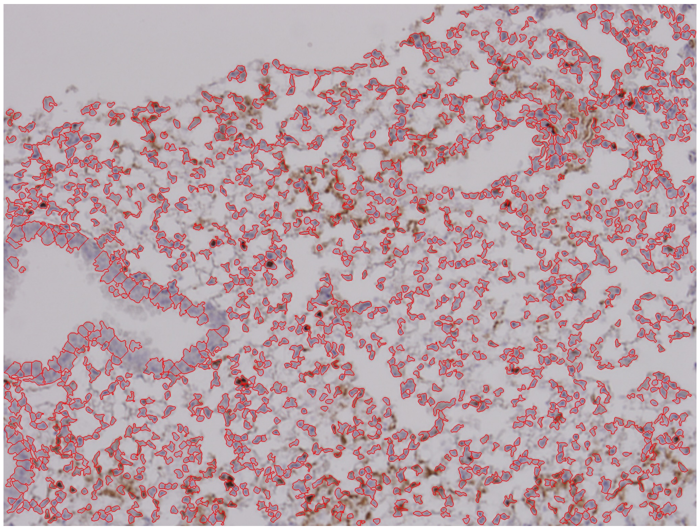

1499 nucleated cells

Supplemental Figure 11
Page 10/30

MIT-T-COVID Lung 3-4
Unchallenged, Female Cohort

CD8$^+$/CD4$^+$ Cell Annotations

Nucleated Cell Annotations

CD8$^+$

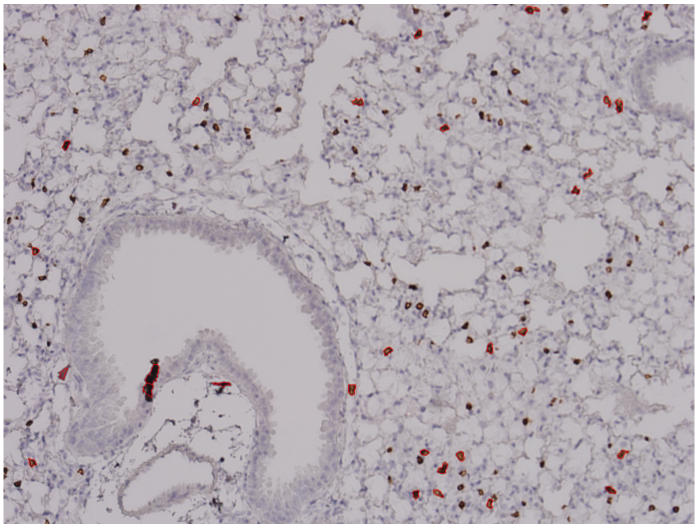

29 CD8$^+$ cells

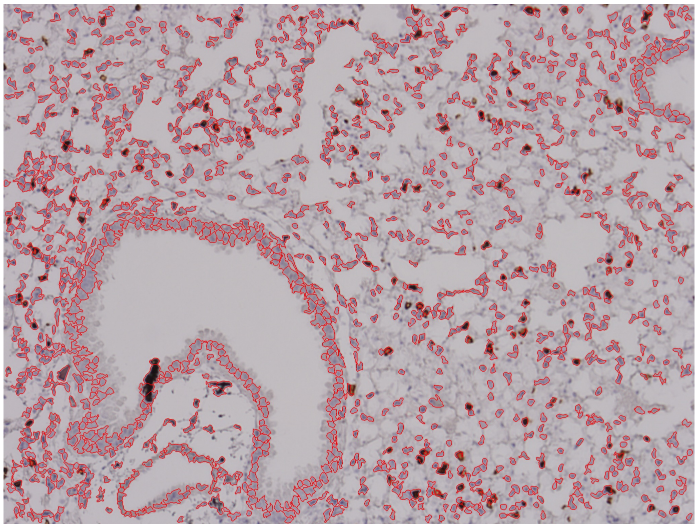

1303 nucleated cells

CD4$^+$

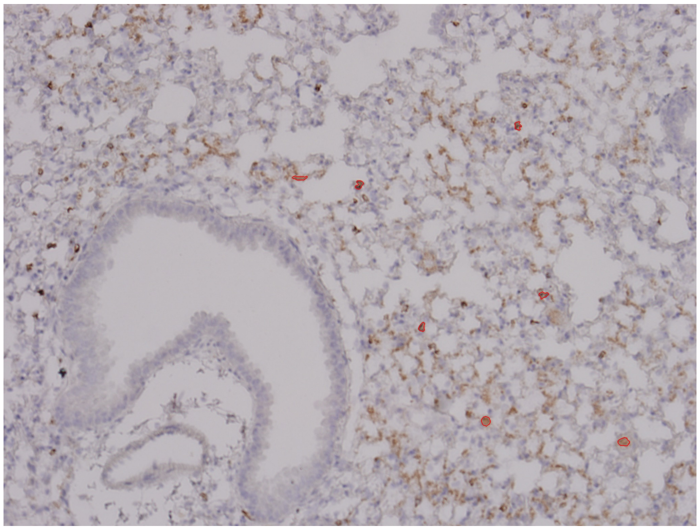

7 CD4$^+$ cells

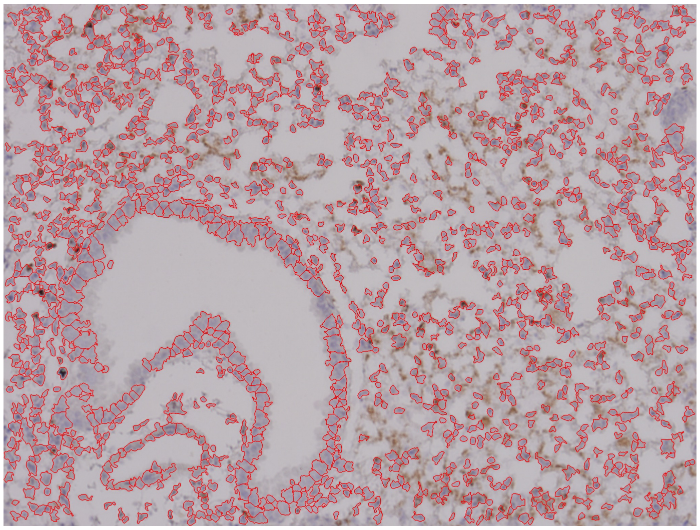

1412 nucleated cells

Pfizer/BNT Lung 1-1
Unchallenged, Female Cohort

CD8⁺/CD4⁺ Cell Annotations | Nucleated Cell Annotations

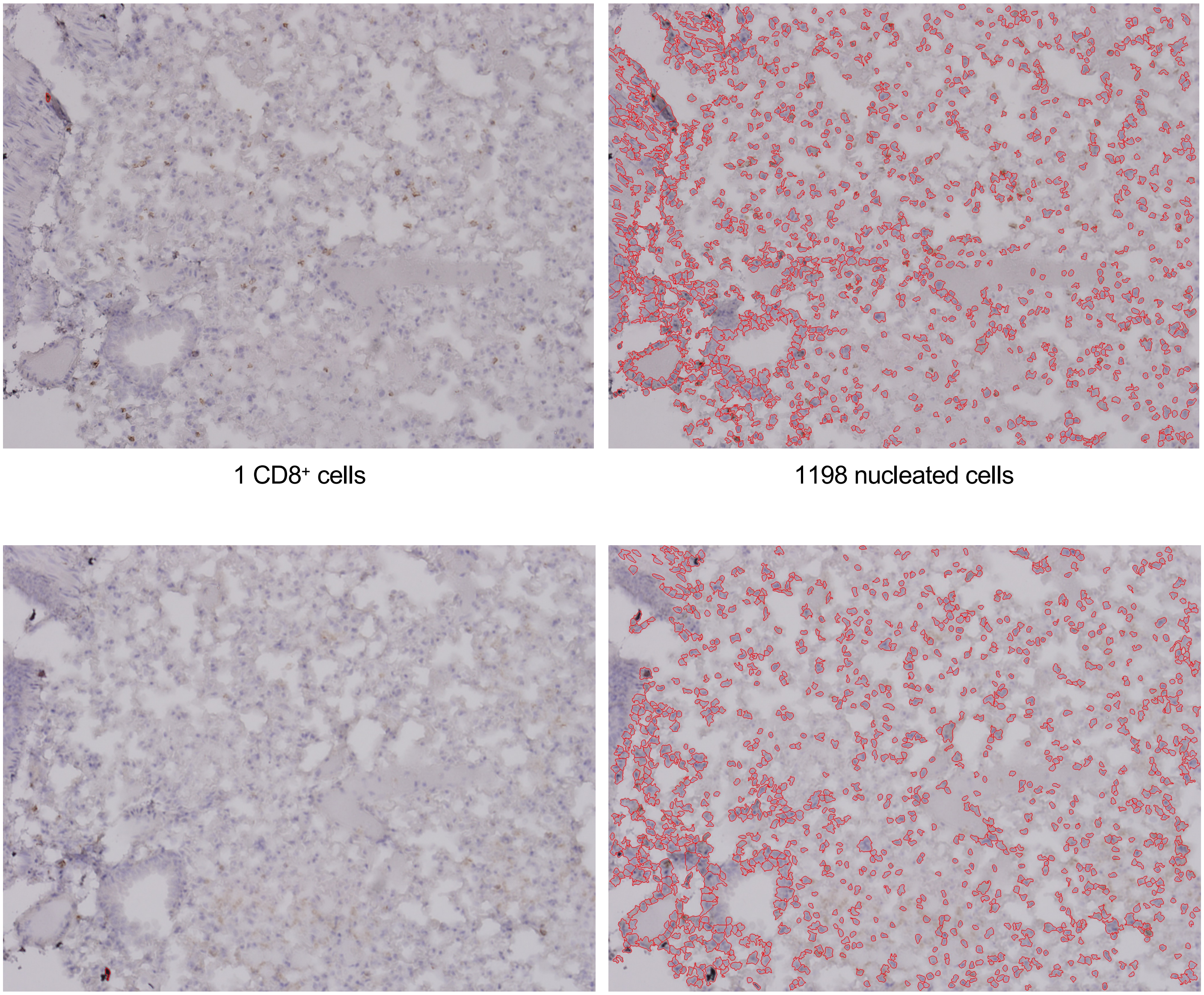

CD8⁺

1 CD8⁺ cells

1198 nucleated cells

CD4⁺

1 CD4⁺ cells

1125 nucleated cells

#### Supplemental Figure 11



##### Pfizer/BNT Lung 1-2
##### Unchallenged, Female Cohort

CD8$^+$/CD4$^+$ Cell Annotations | Nucleated Cell Annotations

CD8$^+$

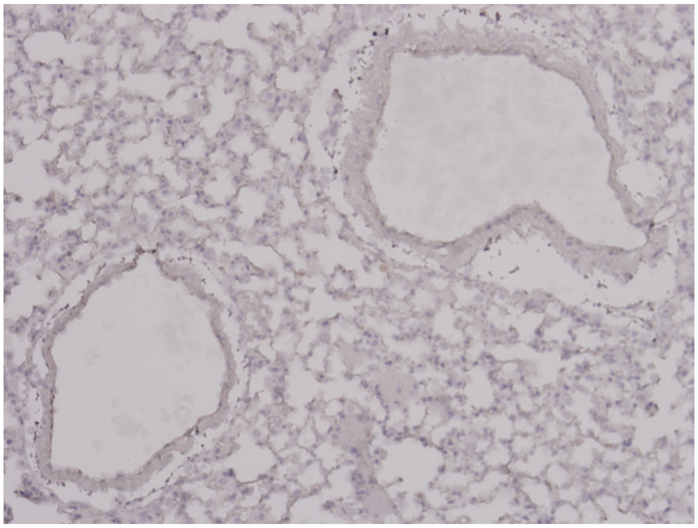

0 CD8$^+$ cells

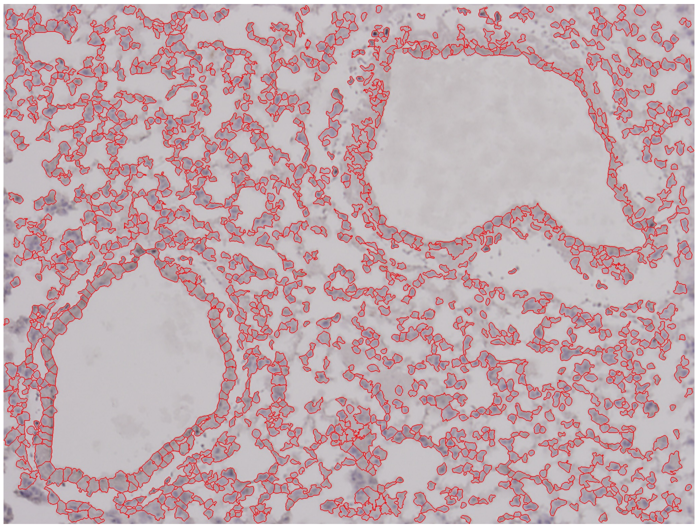

1396 nucleated cells

CD4$^+$

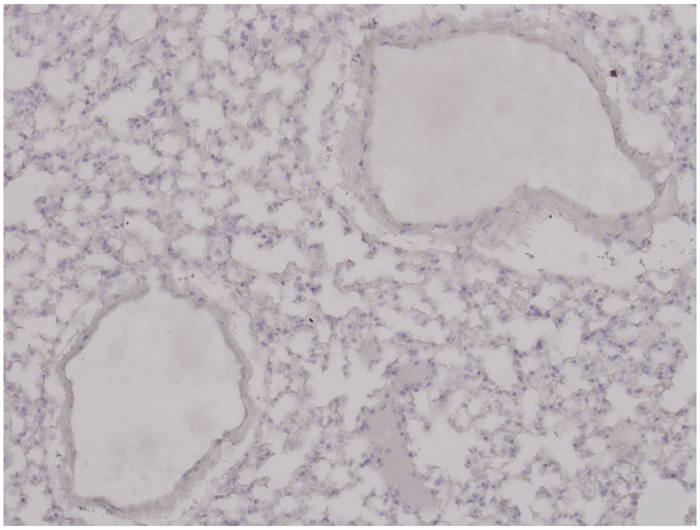

0 CD4$^+$ cells

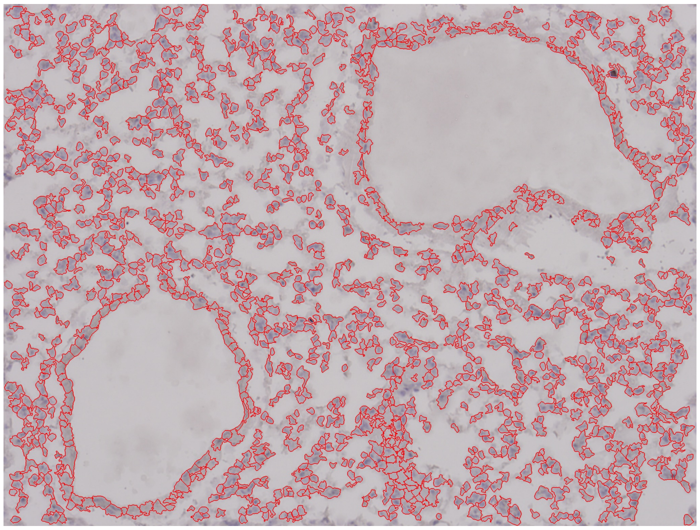

1355 nucleated cells

Supplemental Figure 11
Page 13/30

Pfizer/BNT Lung 2-1
Unchallenged, Female Cohort

CD8$^+$/CD4$^+$ Cell Annotations | Nucleated Cell Annotations

CD8$^+$

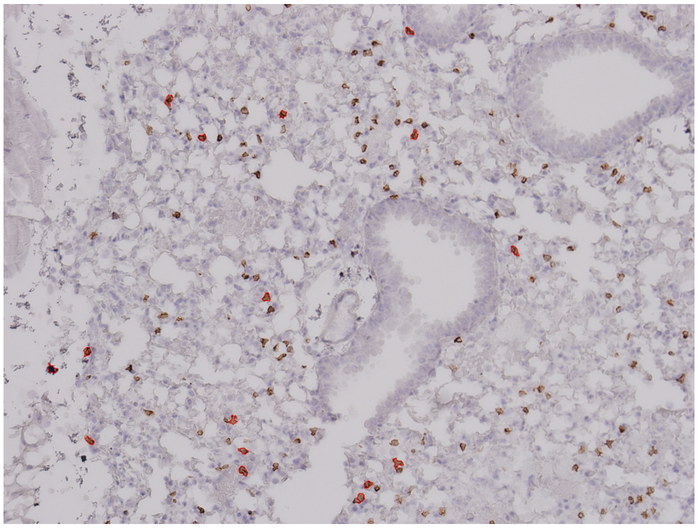

16 CD8$^+$ cells

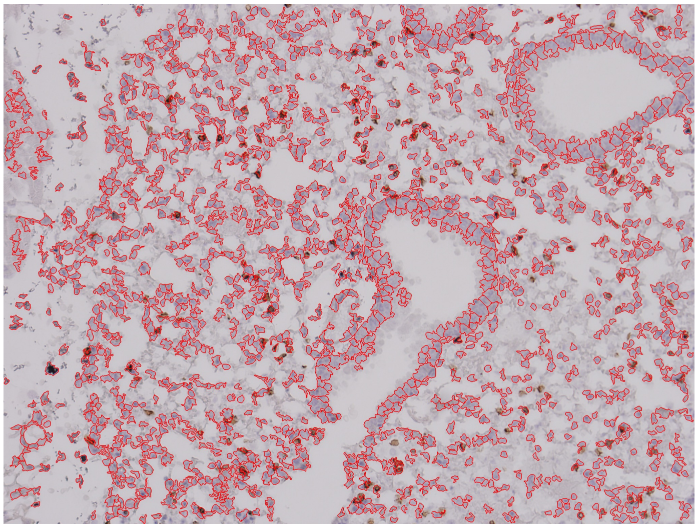

1544 nucleated cells

CD4$^+$

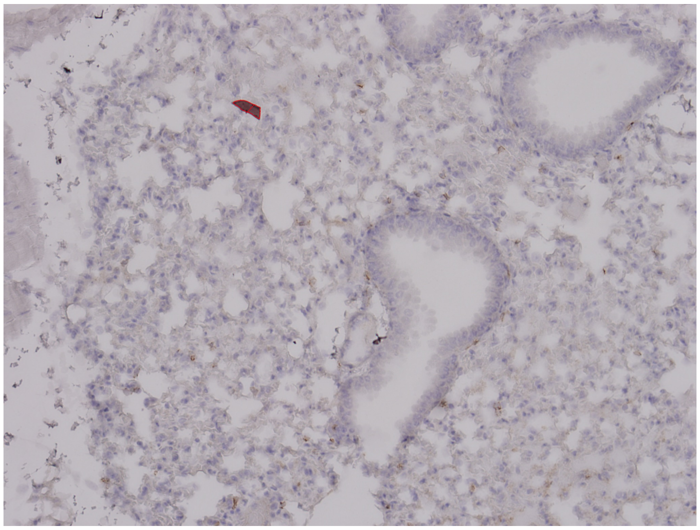

2 CD4$^+$ cells

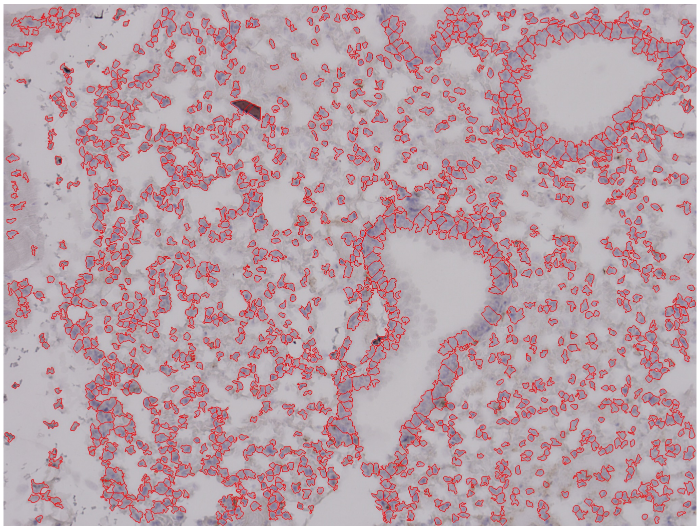

1447 nucleated cells

Pfizer/BNT Lung 2-2
Unchallenged, Female Cohort

CD8$^+$/CD4$^+$ Cell Annotations | Nucleated Cell Annotations

CD8$^+$

CD4$^+$

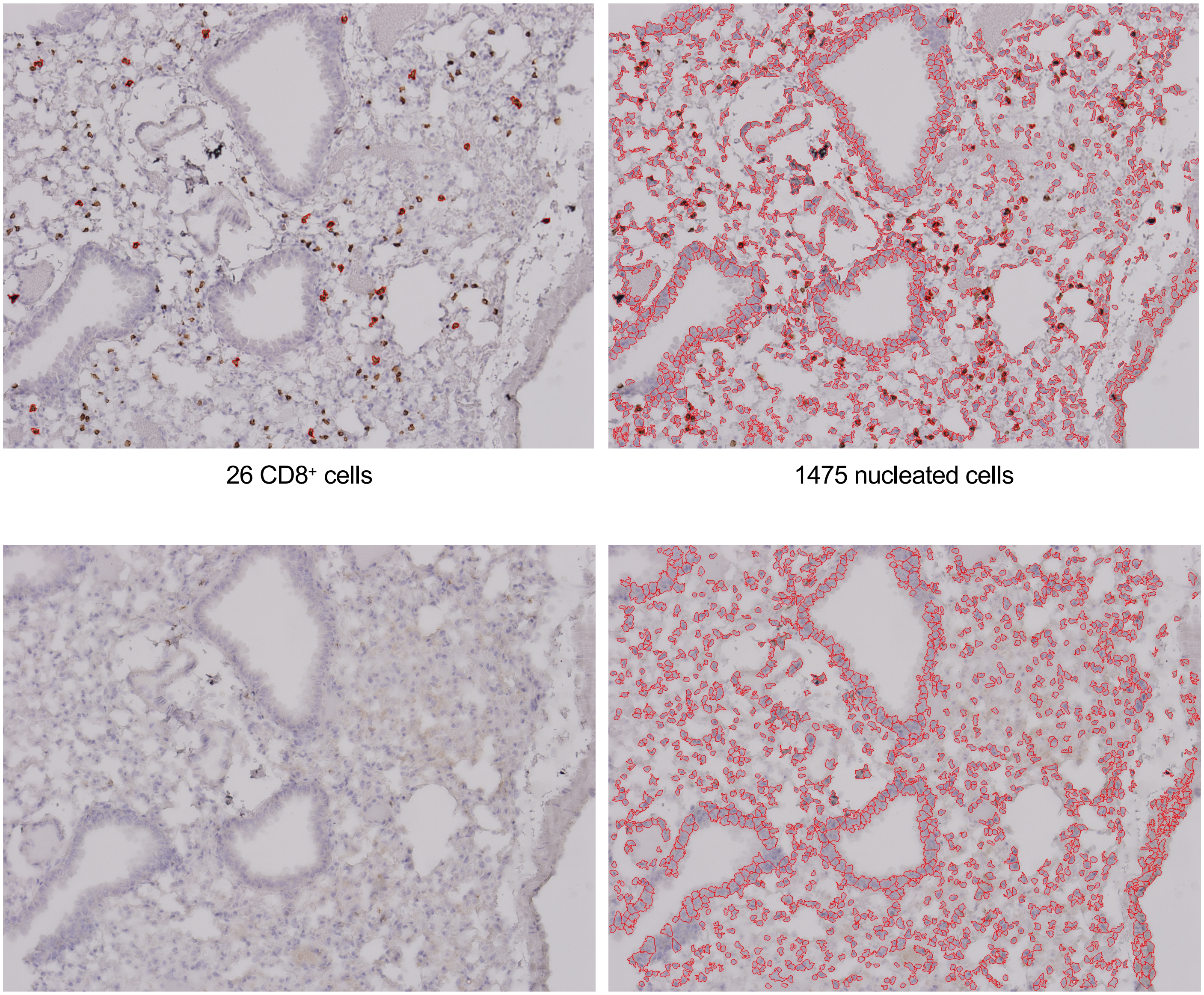

26 CD8$^+$ cells

1475 nucleated cells

0 CD4$^+$ cells

1376 nucleated cells

Supplemental Figure 11


Pfizer/BNT Lung 2-3
Unchallenged, Female Cohort

| | CD8⁺/CD4⁺ Cell Annotations | Nucleated Cell Annotations |
|---|---|---|
| CD8⁺ | 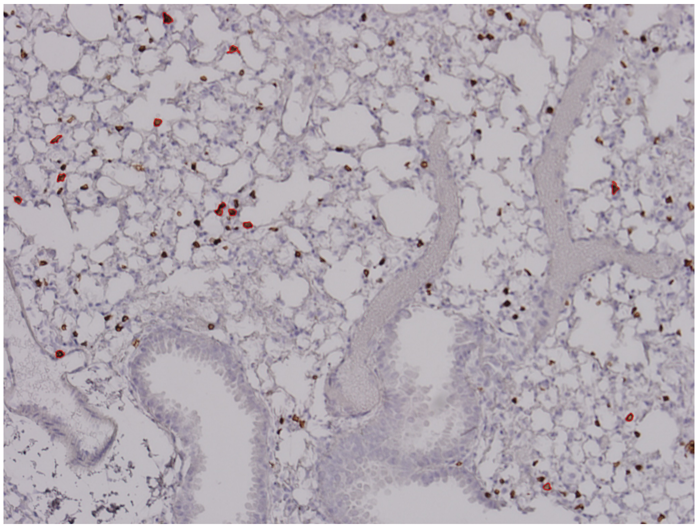 13 CD8⁺ cells | 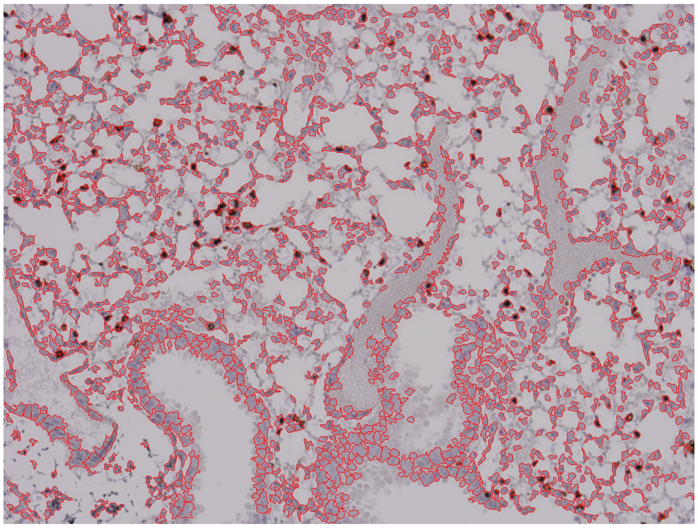 1582 nucleated cells |
| CD4⁺ | 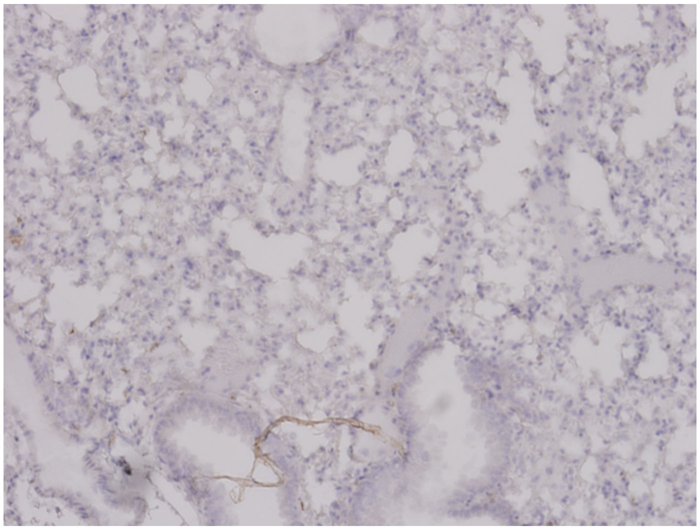 0 CD4⁺ cells | 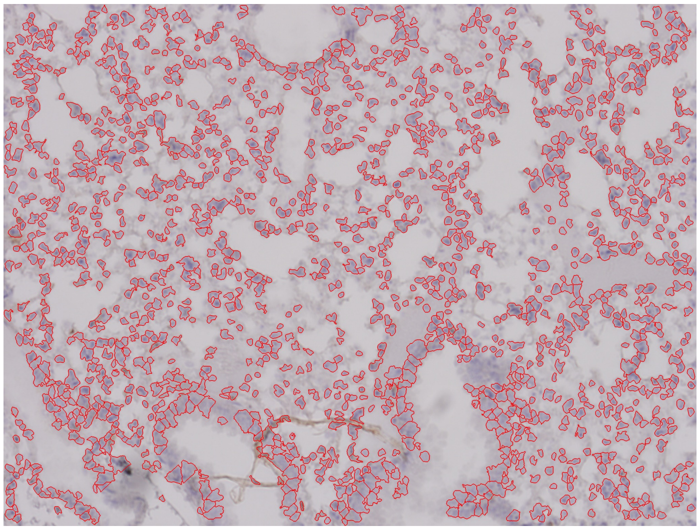 1440 nucleated cells |

Supplemental Figure 11
Page 16/30

Pfizer/BNT Lung 2-4
Unchallenged, Female Cohort

CD8$^+$/CD4$^+$ Cell Annotations

Nucleated Cell Annotations

CD8$^+$

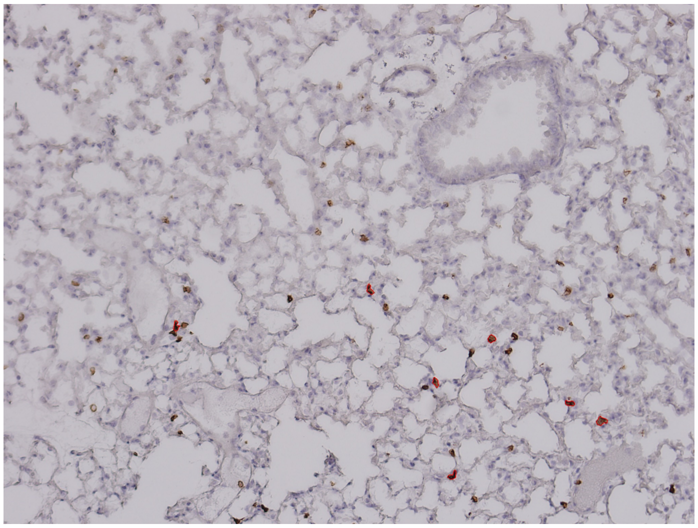

7 CD8$^+$ cells

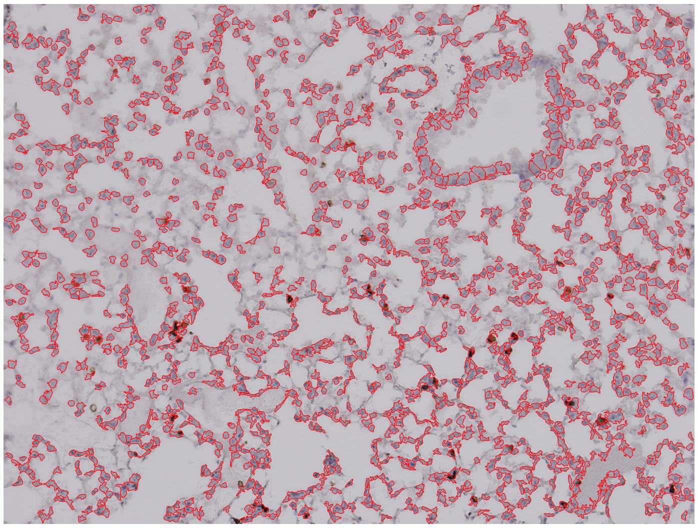

1389 nucleated cells

CD4$^+$

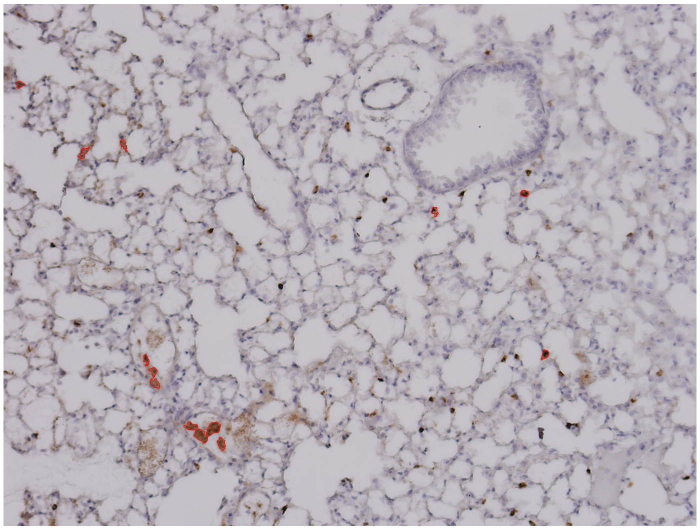

13 CD4$^+$ cells

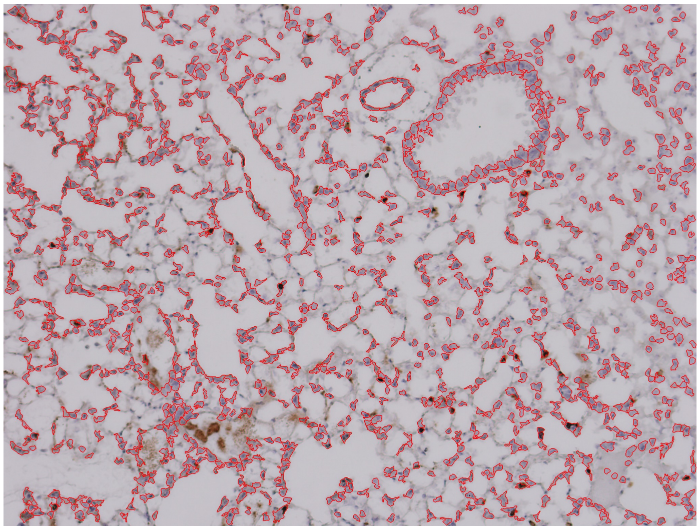

1197 nucleated cells

Supplemental Figure 11
Page 17/30

Pfizer/BNT Lung 3-1
Unchallenged, Female Cohort

CD8$^+$/CD4$^+$ Cell Annotations | Nucleated Cell Annotations

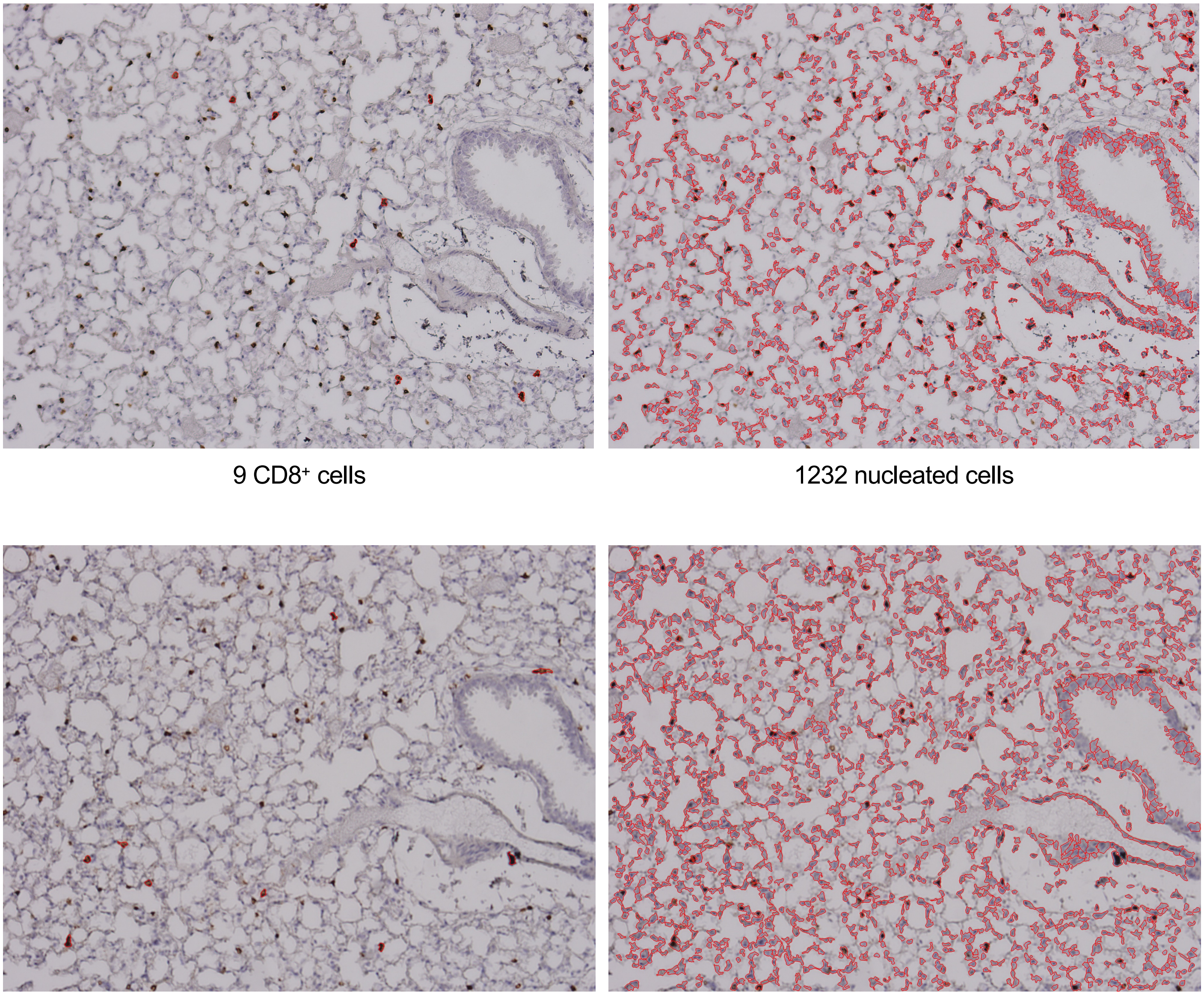

Supplemental Figure 11
Page 18/30

Pfizer/BNT Lung 3-2
Unchallenged, Female Cohort

CD8$^+$/CD4$^+$ Cell Annotations | Nucleated Cell Annotations

CD8$^+$

CD4$^+$

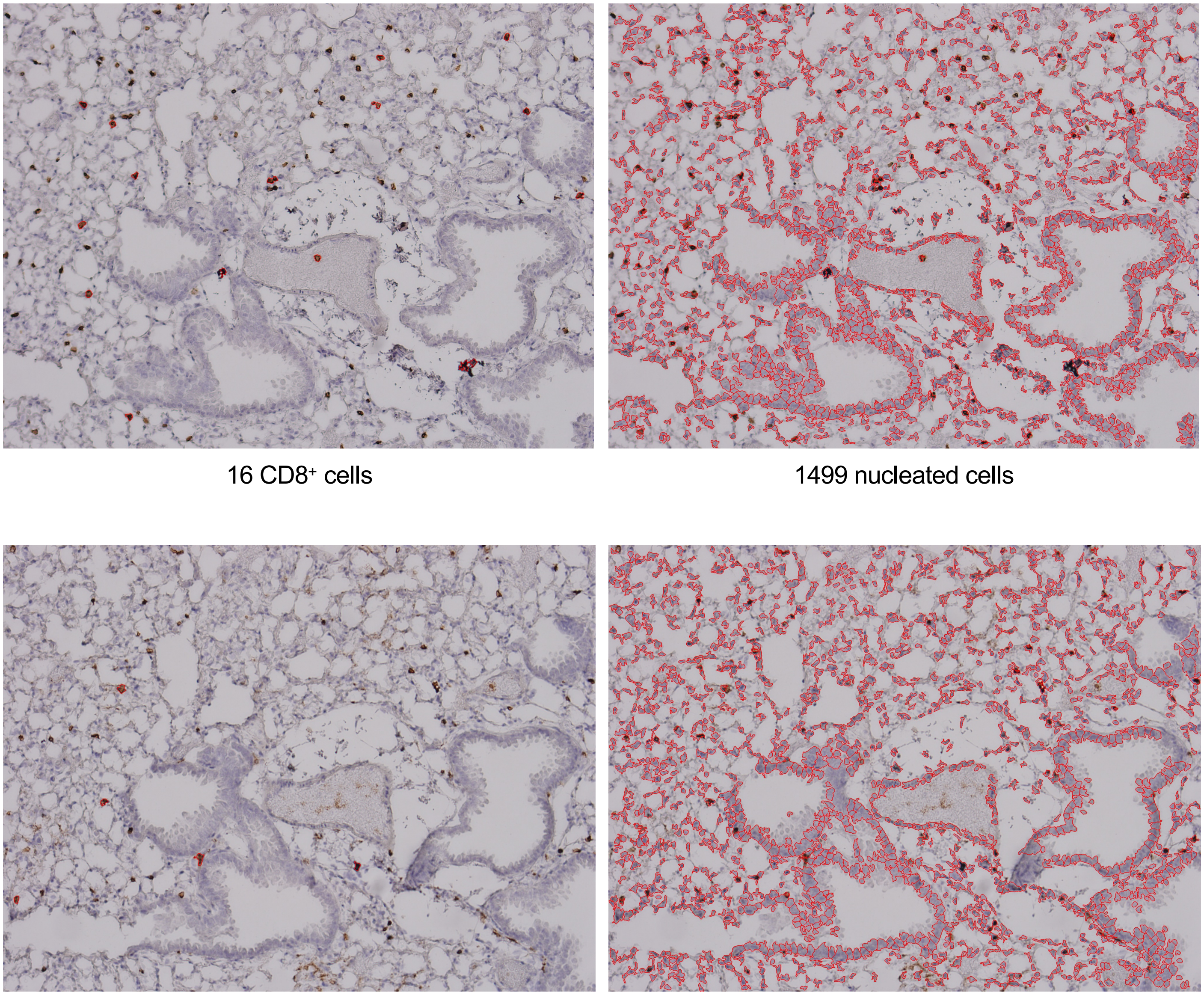

16 CD8$^+$ cells | 1499 nucleated cells

5 CD4$^+$ cells | 1464 nucleated cells

Supplemental Figure 11
Page 19/30

Pfizer/BNT Lung 3-3
Unchallenged, Female Cohort

CD8$^+$/CD4$^+$ Cell Annotations | Nucleated Cell Annotations

CD8$^+$

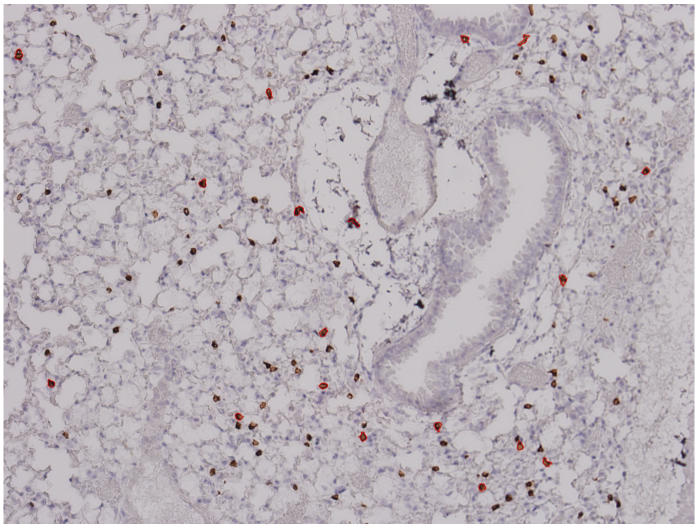

18 CD8$^+$ cells

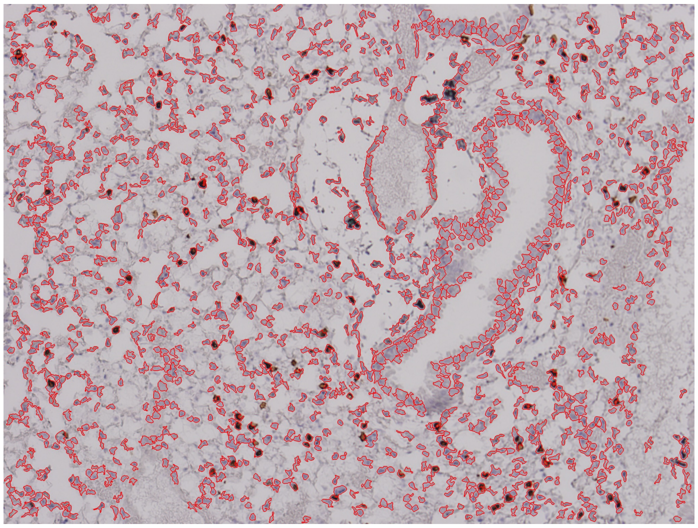

1365 nucleated cells

CD4$^+$

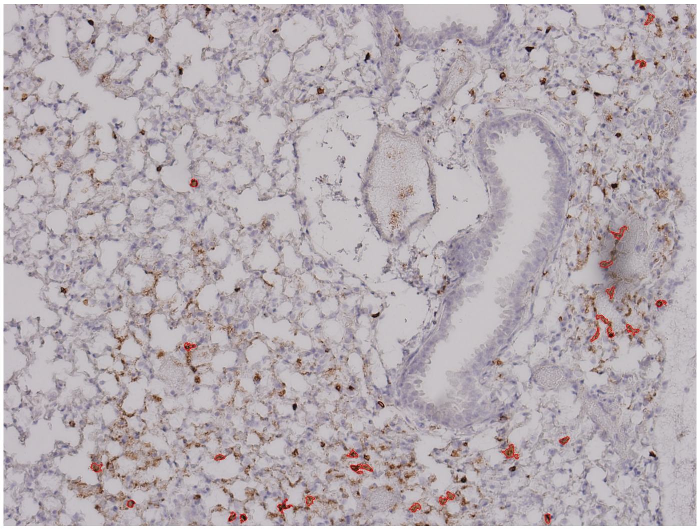

28 CD4$^+$ cells

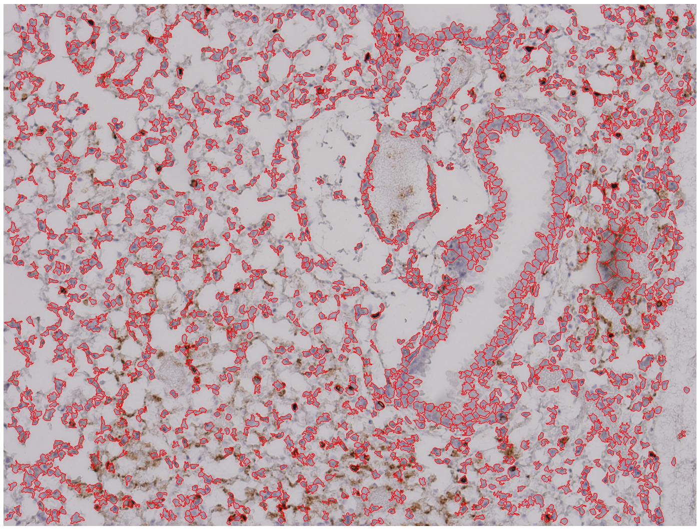

1541 nucleated cells

Supplemental Figure 11
Page 20/30

Pfizer/BNT Lung 3-4
Unchallenged, Female Cohort

CD8$^+$/CD4$^+$ Cell Annotations

Nucleated Cell Annotations

CD8$^+$

CD4$^+$

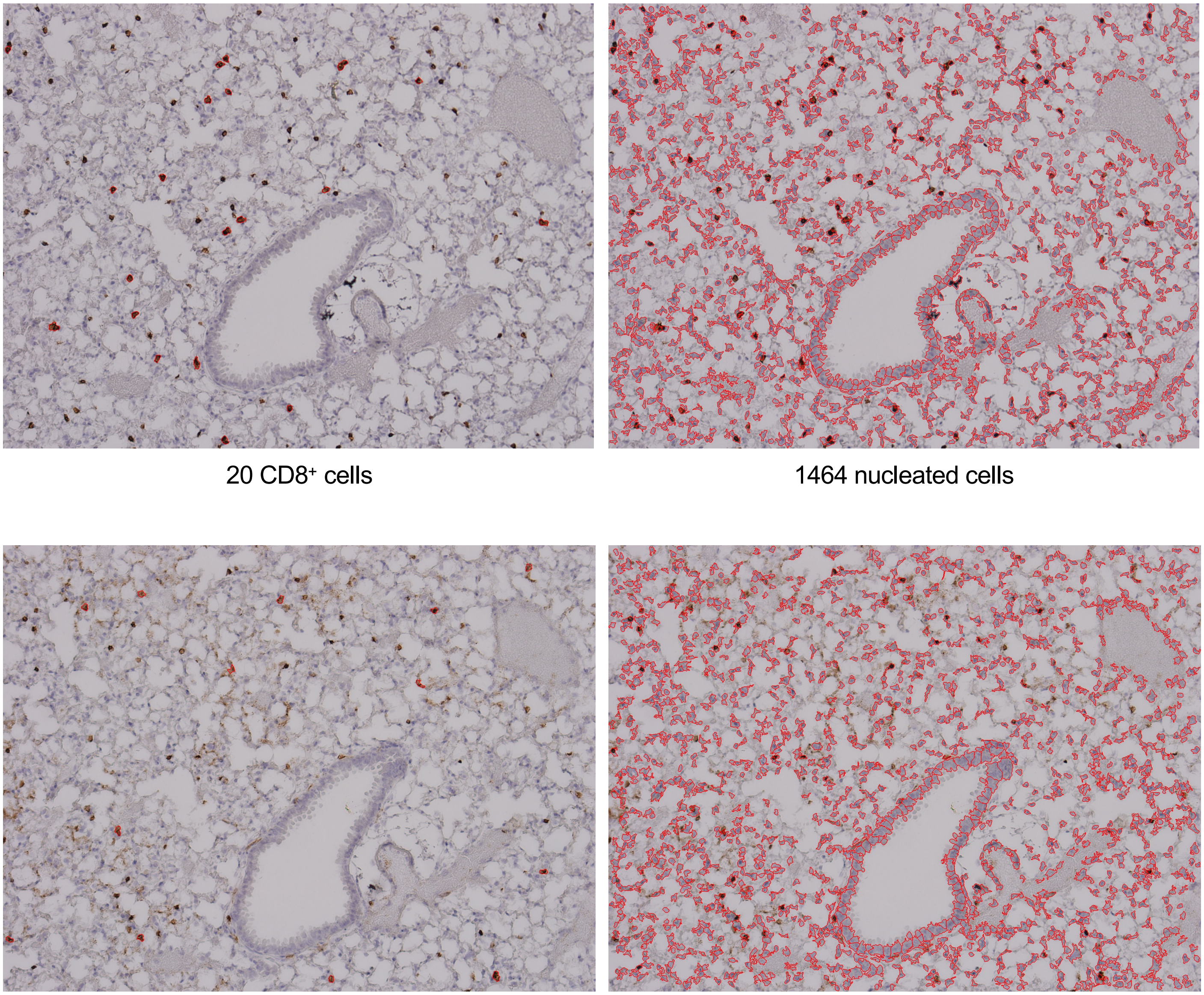

20 CD8$^+$ cells

1464 nucleated cells

13 CD4$^+$ cells

1400 nucleated cells

Supplemental Figure 11


PBS Lung 1-1
Unchallenged, Female Cohort

CD8⁺/CD4⁺ Cell Annotations — Nucleated Cell Annotations

CD8⁺

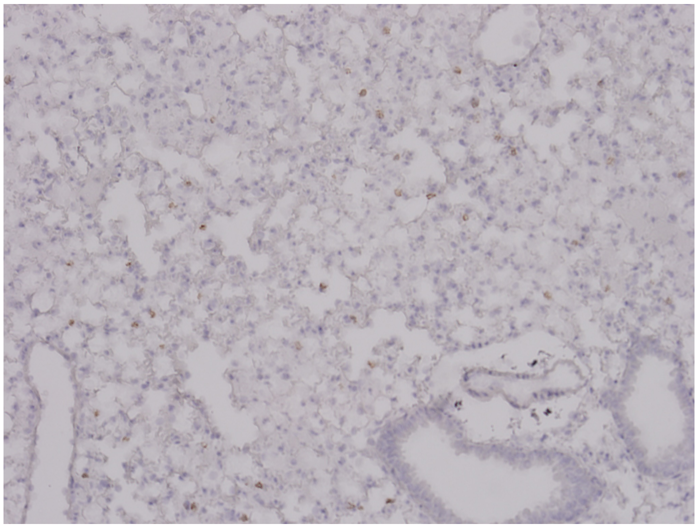

0 CD8⁺ cells

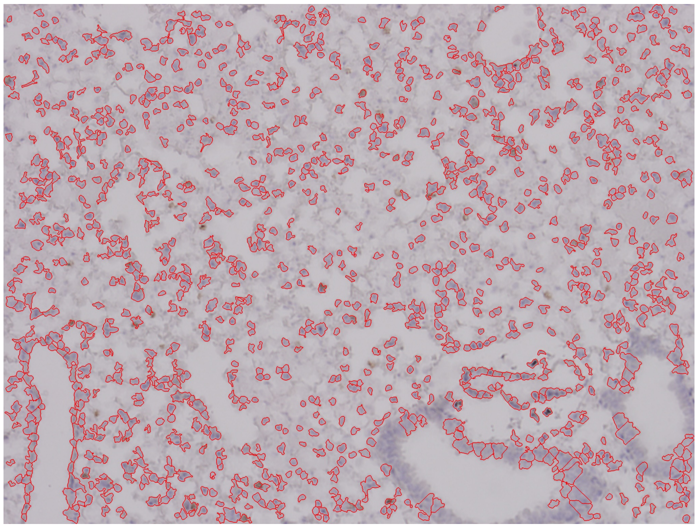

1136 nucleated cells

CD4⁺

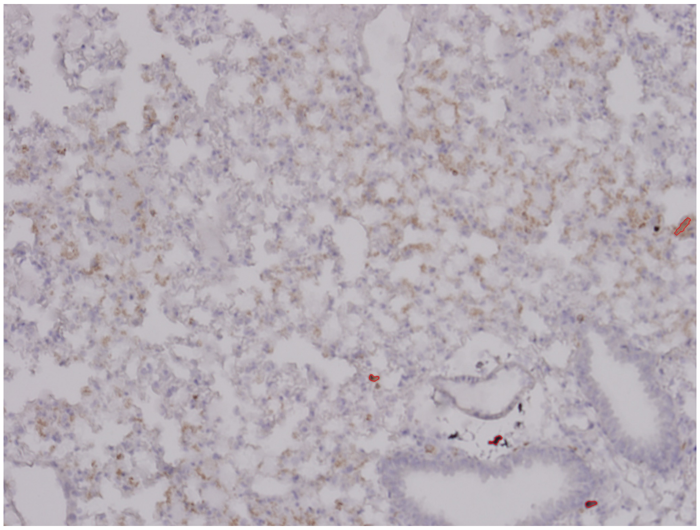

4 CD4⁺ cells

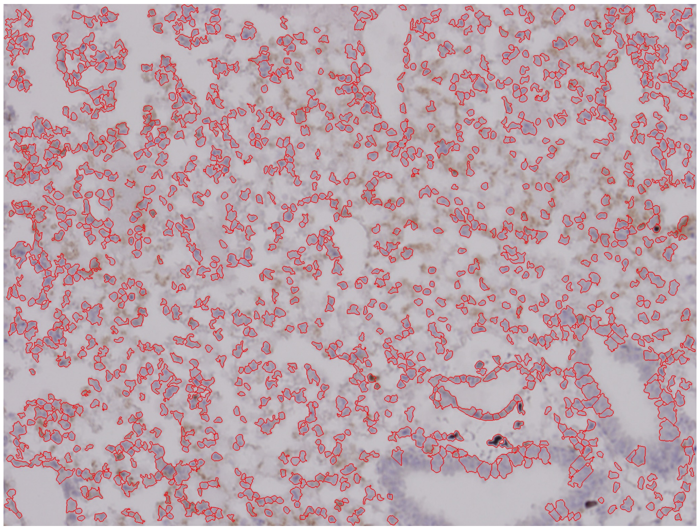

1299 nucleated cells

### Supplemental Figure 11


#### PBS Lung 1-2
#### Unchallenged, Female Cohort

CD8$^+$/CD4$^+$ Cell Annotations | Nucleated Cell Annotations

CD8$^+$

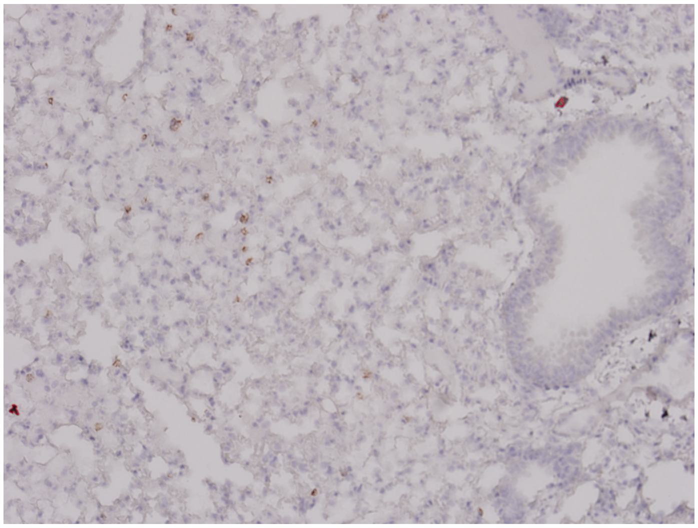

2 CD8$^+$ cells

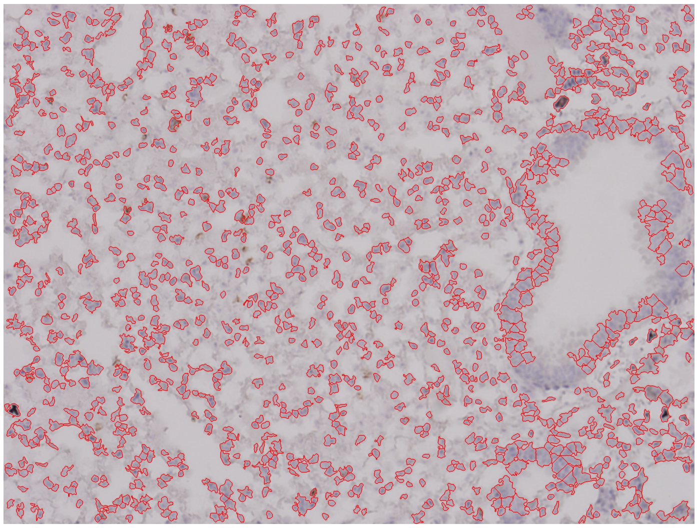

1190 nucleated cells

CD4$^+$

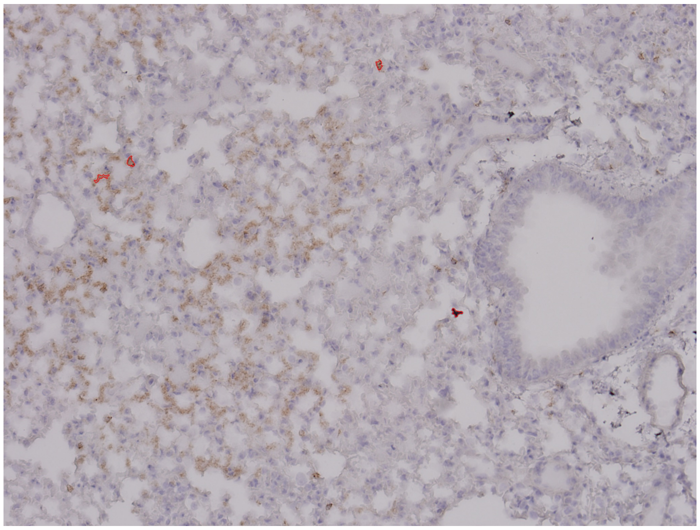

4 CD4$^+$ cells

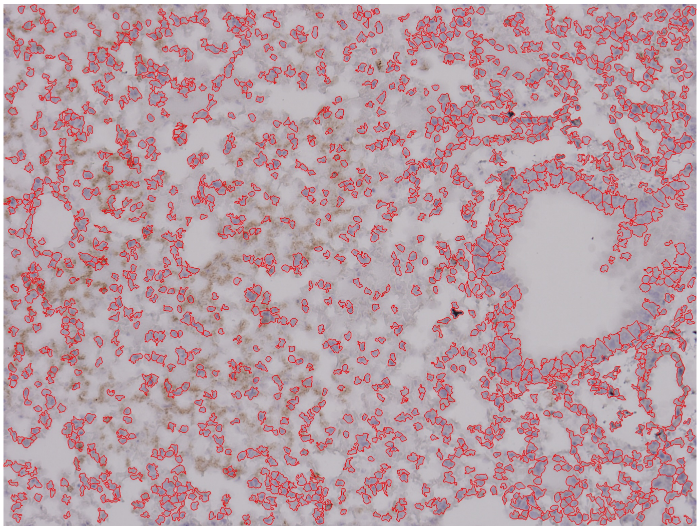

1497 nucleated cells

Supplemental Figure 11


PBS Lung 2-1
Unchallenged, Female Cohort

| | CD8⁺/CD4⁺ Cell Annotations | Nucleated Cell Annotations |
|---|---|---|

CD8⁺/CD4⁺ Cell Annotations

Nucleated Cell Annotations

$CD8^+$

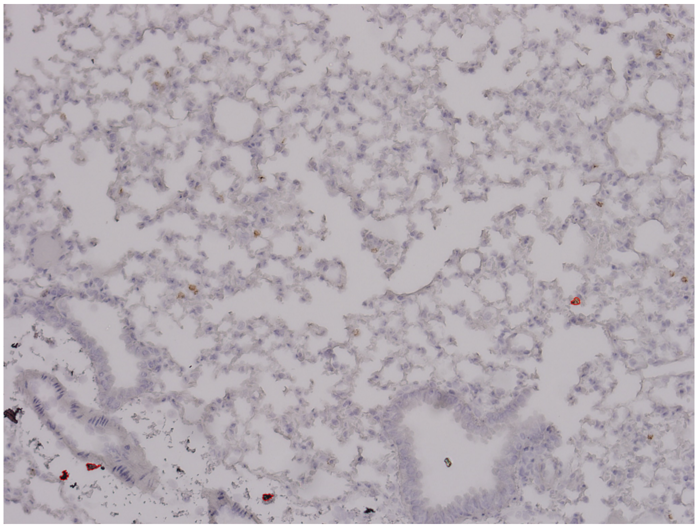

4 $CD8^+$ cells

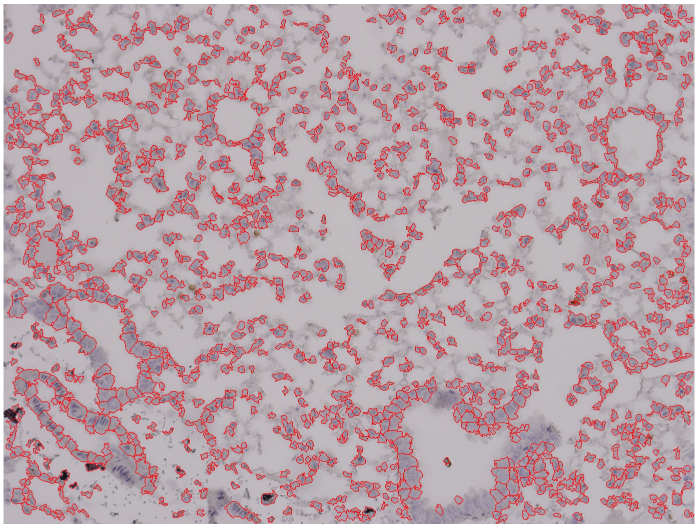

1307 nucleated cells

$CD4^+$

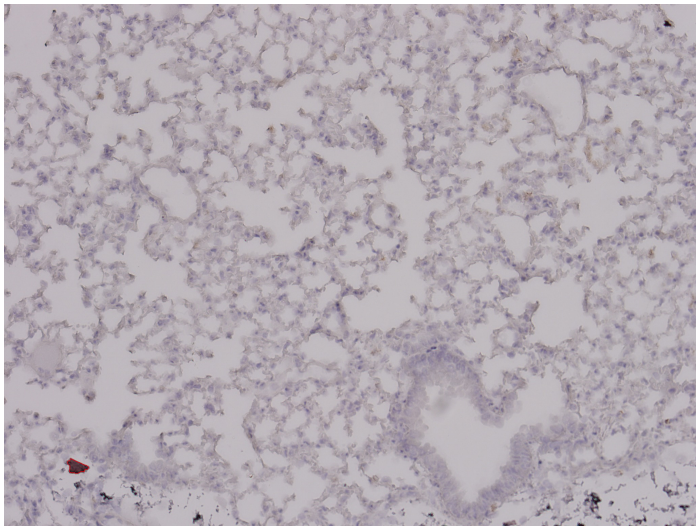

1 $CD4^+$ cells

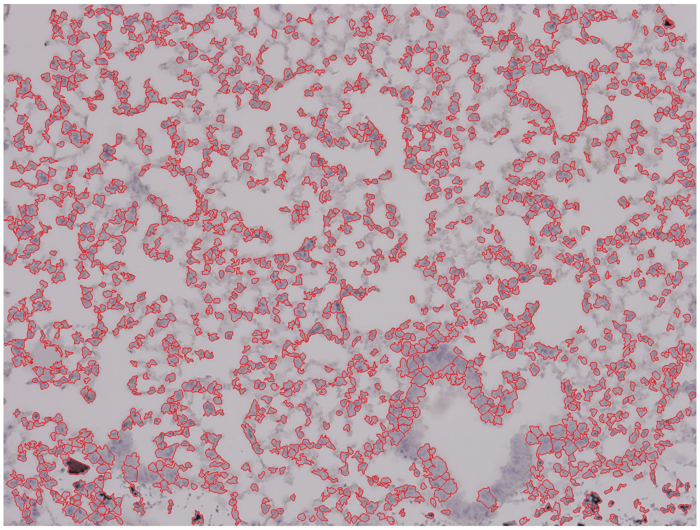

1452 nucleated cells

Supplemental Figure 11


PBS Lung 2-2
Unchallenged, Female Cohort

CD8$^+$/CD4$^+$ Cell Annotations | Nucleated Cell Annotations

CD8$^+$

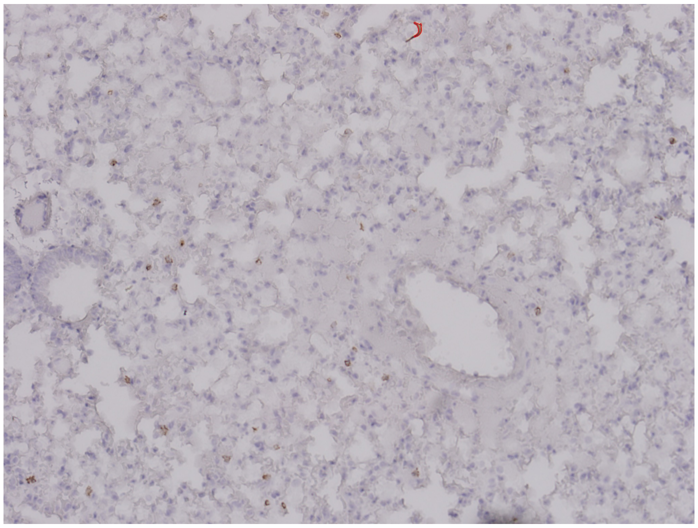

1 CD8$^+$ cells

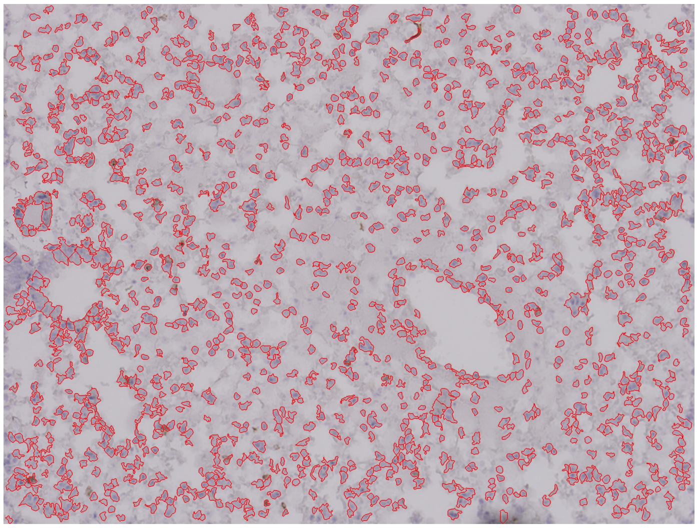

1316 nucleated cells

CD4$^+$

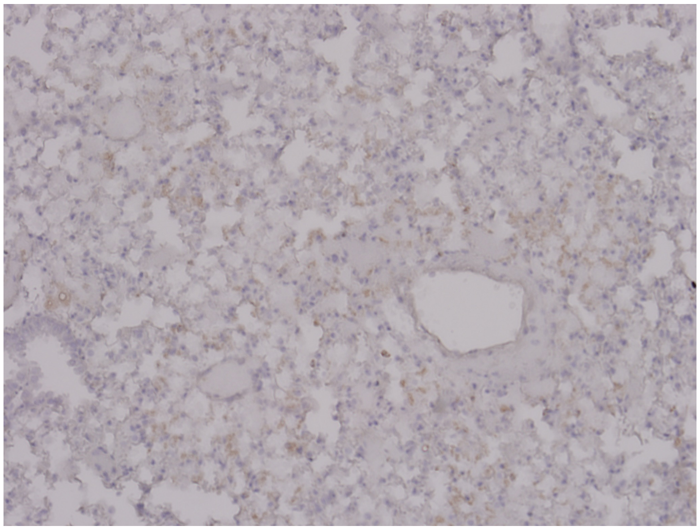

0 CD4$^+$ cells

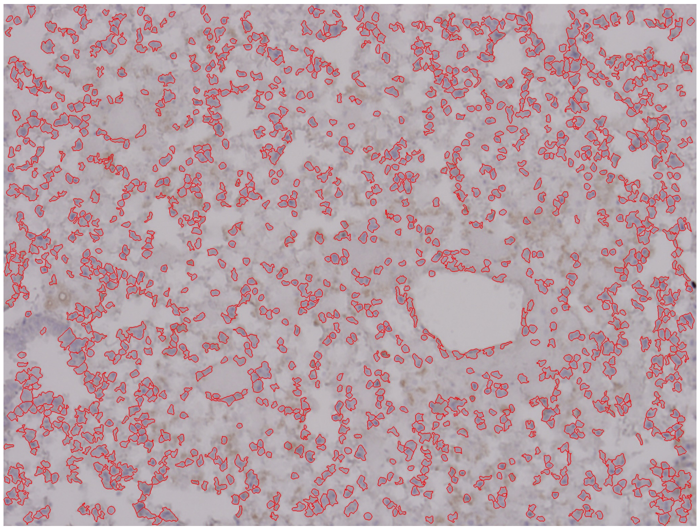

1304 nucleated cells

Supplemental Figure 11


PBS Lung 2-3
Unchallenged, Female Cohort

| | CD8$^+$/CD4$^+$ Cell Annotations | Nucleated Cell Annotations |
| --- | --- | --- |
| CD8$^+$ | 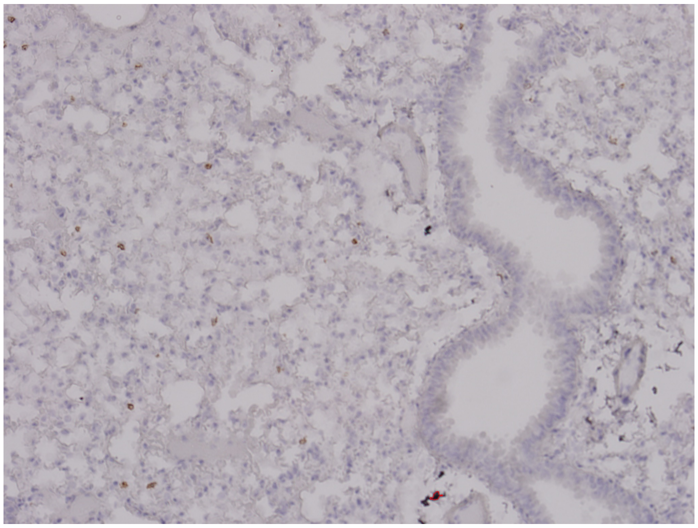 1 CD8$^+$ cells | 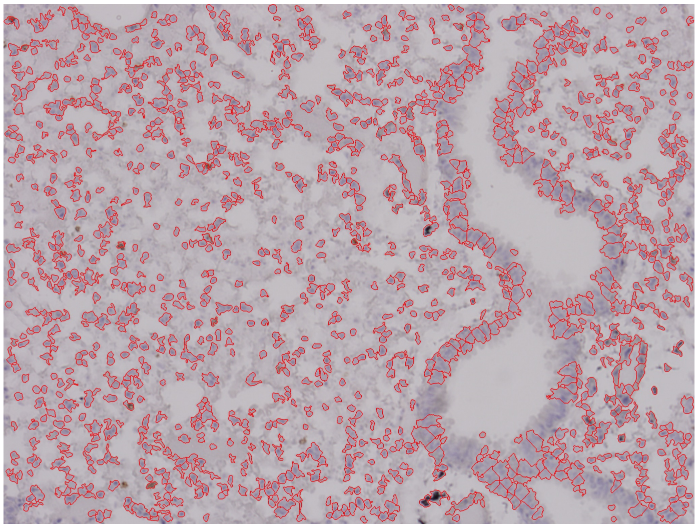 1275 nucleated cells |
| CD4$^+$ | 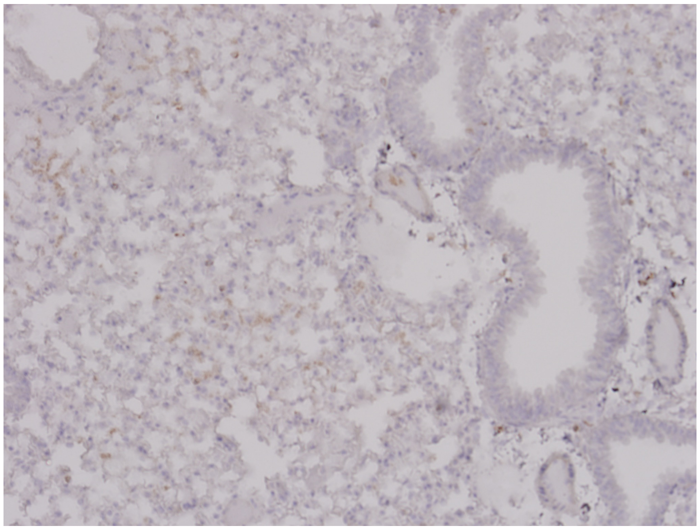 0 CD4$^+$ cells | 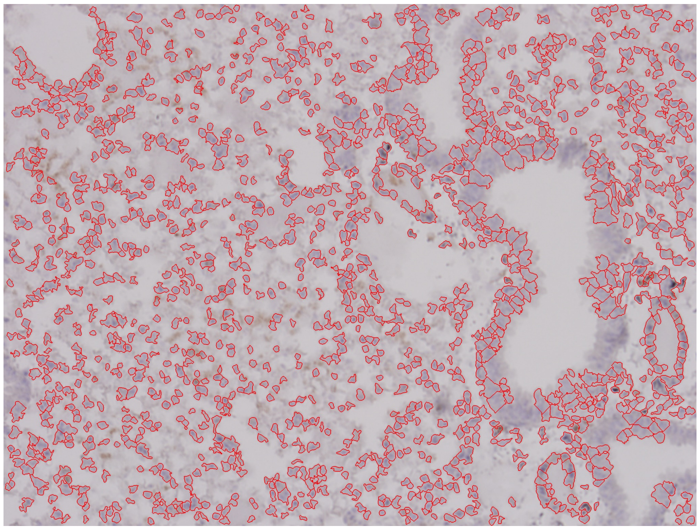 1337 nucleated cells |

Supplemental Figure 11
Page 26/30

PBS Lung 2-4
Unchallenged, Female Cohort

CD8$^+$/CD4$^+$ Cell Annotations | Nucleated Cell Annotations

CD8$^+$

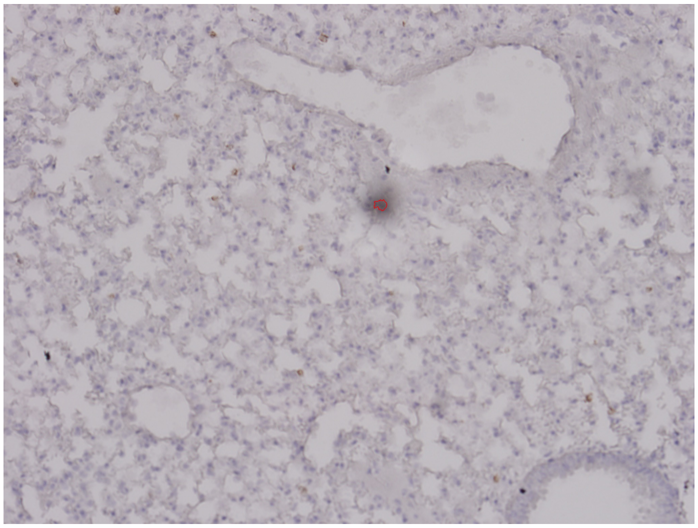

1 CD8$^+$ cells

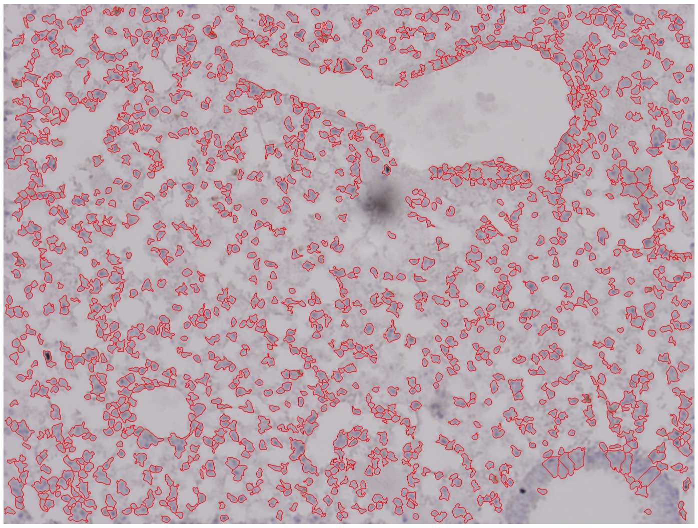

1308 nucleated cells

CD4$^+$

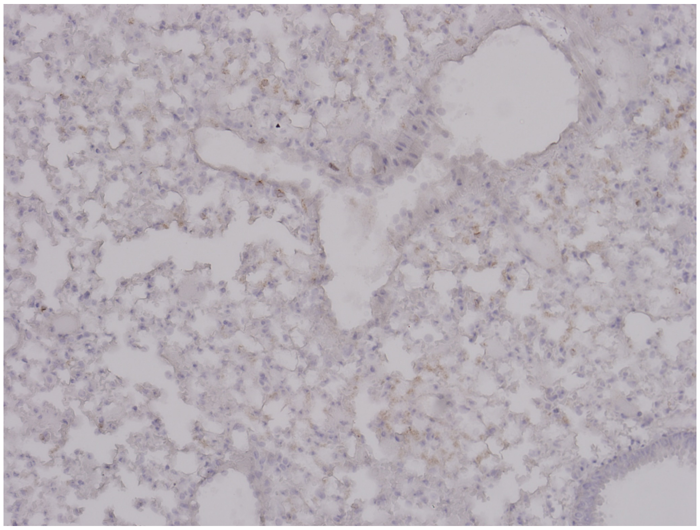

0 CD4$^+$ cells

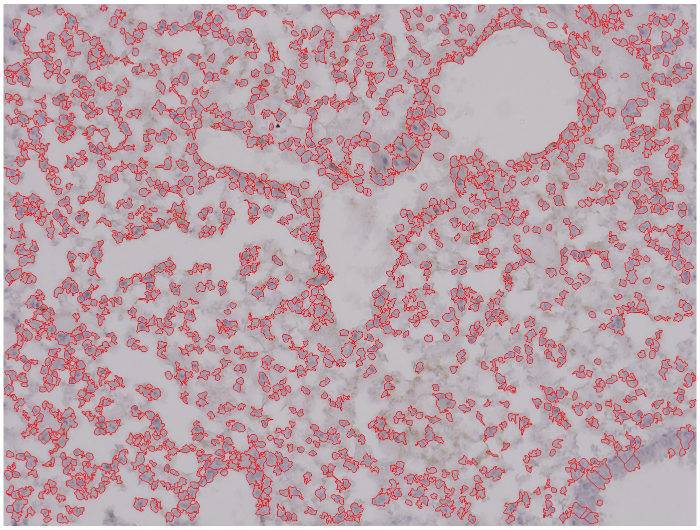

1427 nucleated cells

### Supplemental Figure 11



#### PBS Lung 3-1
#### Unchallenged, Female Cohort

CD8$^+$/CD4$^+$ Cell Annotations | Nucleated Cell Annotations

CD8$^+$

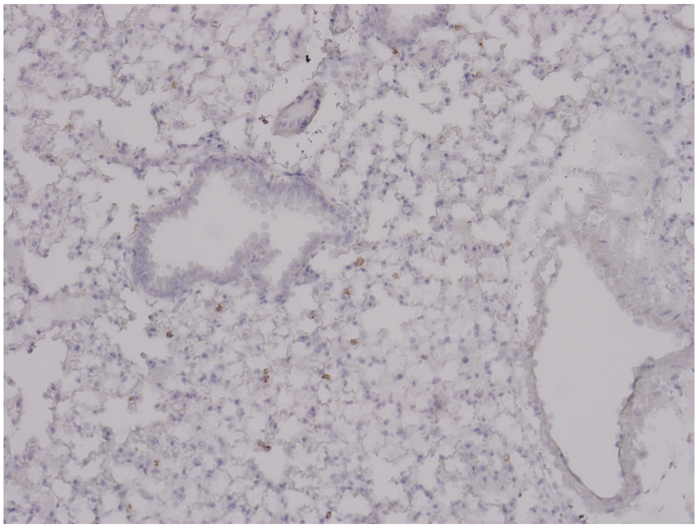

0 CD8$^+$ cells

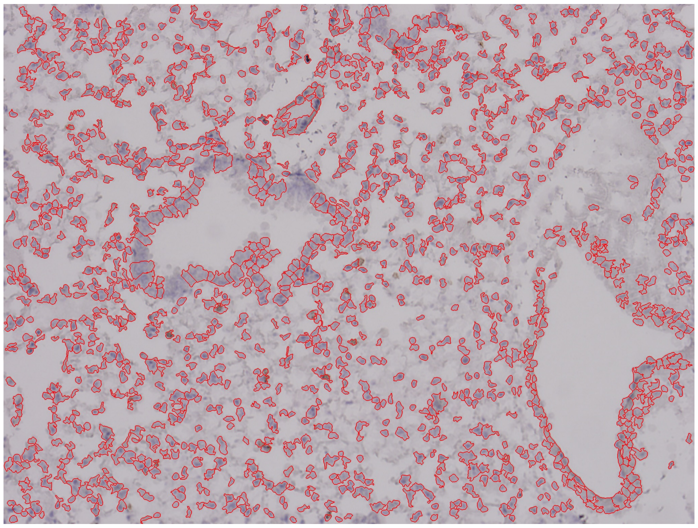

1322 nucleated cells

CD4$^+$

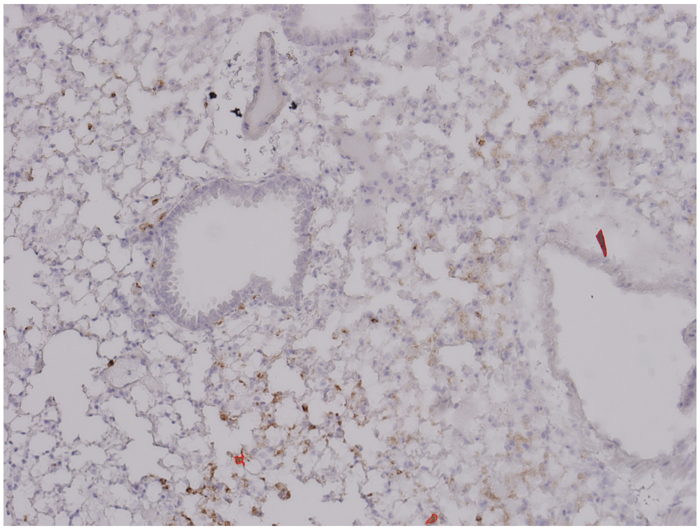

3 CD4$^+$ cells

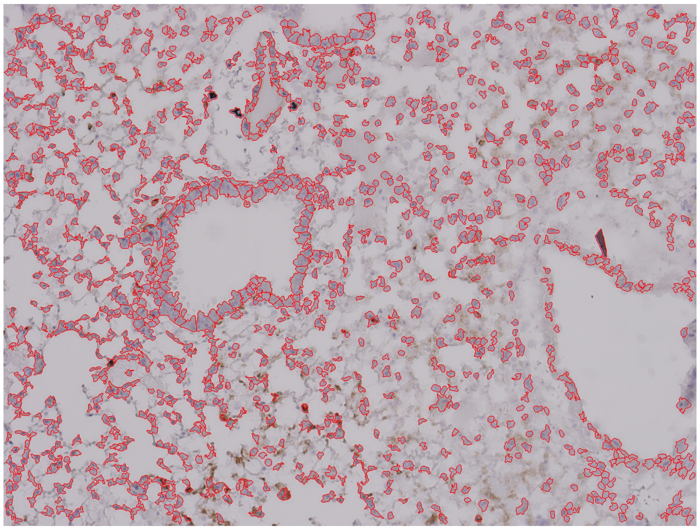

1207 nucleated cells

Supplemental Figure 11
Page 28/30

PBS Lung 3-2
Unchallenged, Female Cohort

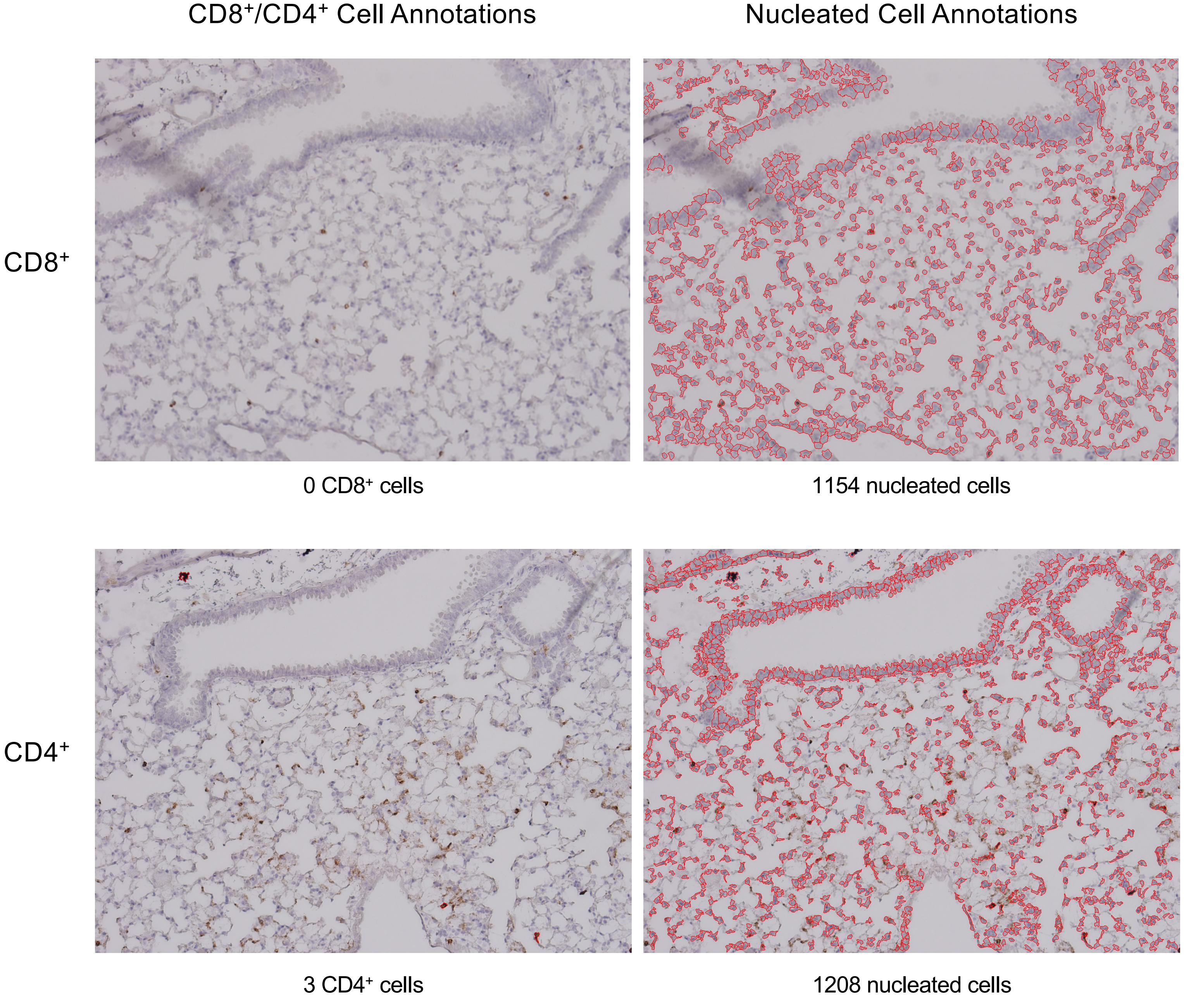

Supplemental Figure 11
Page 29/30

PBS Lung 3-3
Unchallenged, Female Cohort

CD8$^+$/CD4$^+$ Cell Annotations

Nucleated Cell Annotations

CD8$^+$

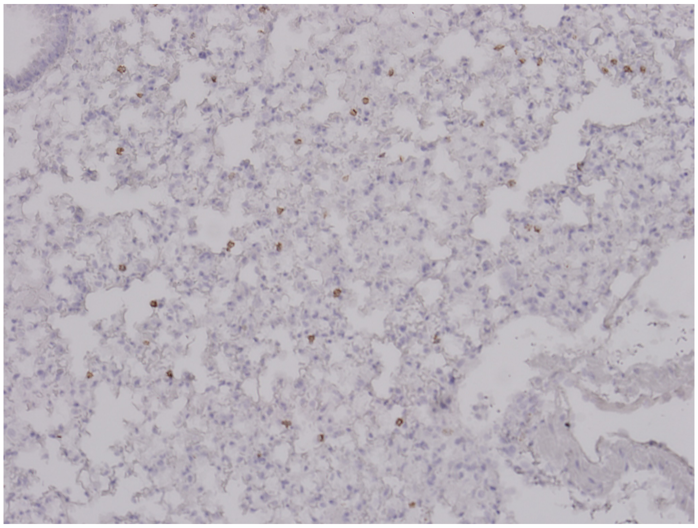

0 CD8$^+$ cells

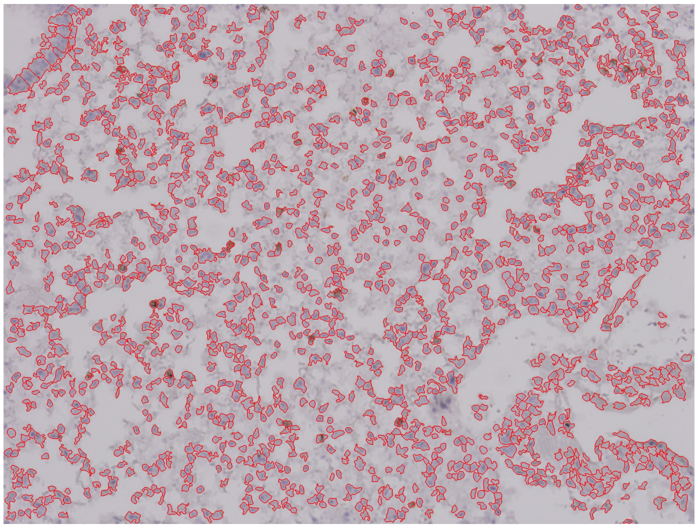

1509 nucleated cells

CD4$^+$

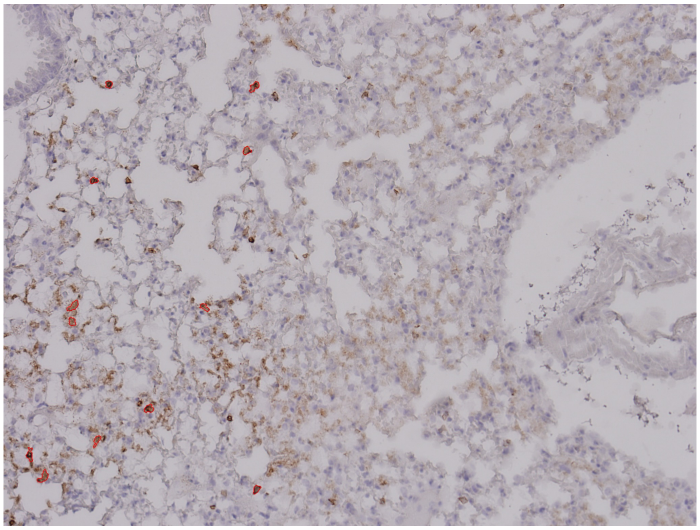

12 CD4$^+$ cells

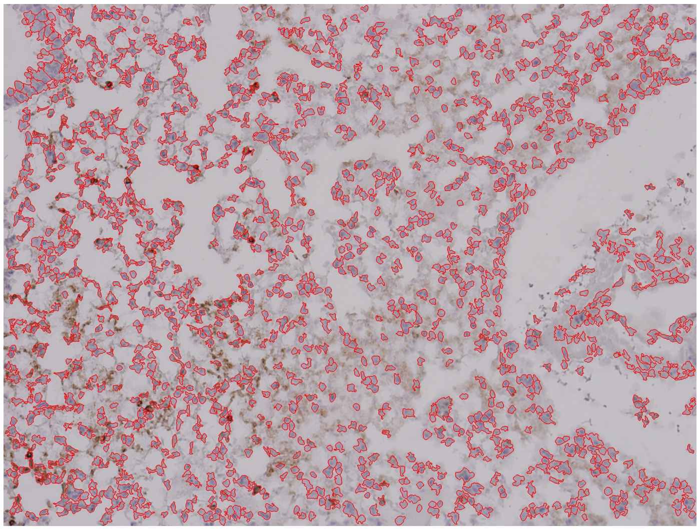

1402 nucleated cells

Supplemental Figure 11
Page 30/30

PBS Lung 3-4
Unchallenged, Female Cohort

CD8$^+$/CD4$^+$ Cell Annotations

Nucleated Cell Annotations

CD8$^+$

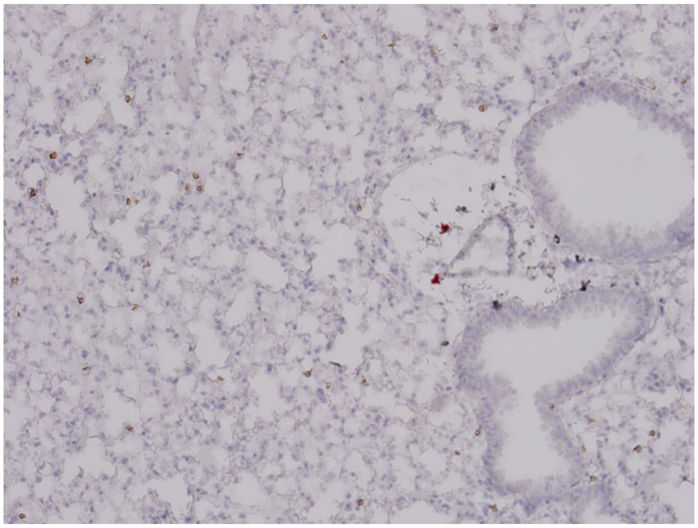

2 CD8$^+$ cells

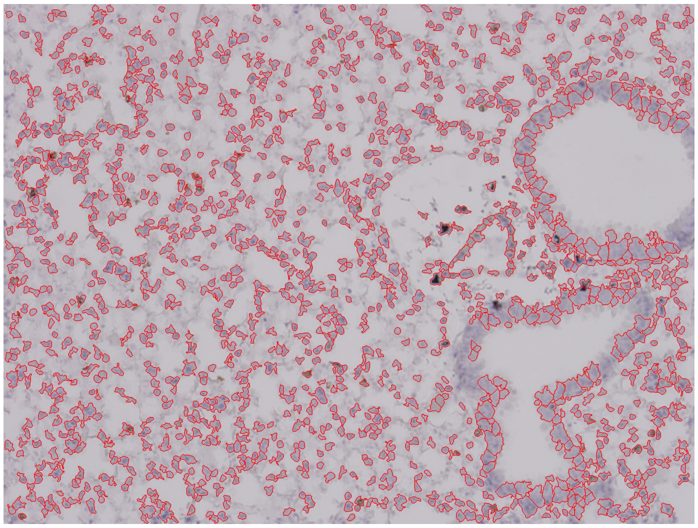

1349 nucleated cells

CD4$^+$

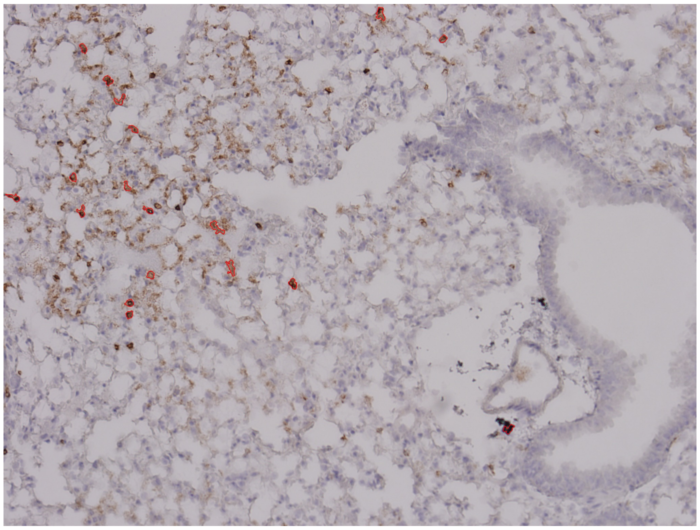

18 CD4$^+$ cells

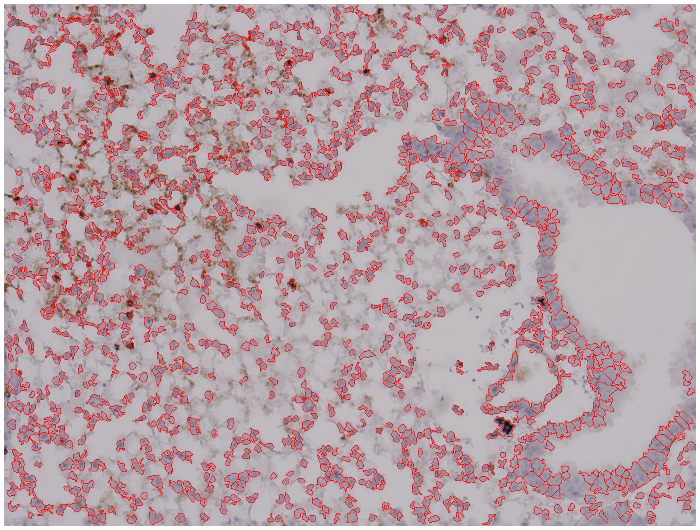

1316 nucleated cells
